# An aerosolised dual-action Autotaxin inhibitor-PPARγ agonist for the treatment of pulmonary fibrosis

**DOI:** 10.1101/2025.11.12.687859

**Authors:** Alexios N. Matralis, Elli-Anna Stylianaki, Eleni Ladopoulou, Paraskevi Kanellopoulou, Stefanos Smyrniotis, Christiana Magkrioti, Konstantinos D. Papavasileiou, Sabine Willems, Juan Pablo Rincon Pabon, Dimitris Nastos, Alexandros Galaras, Skarlatos G. Dedos, Eleanna Kaffe, Pantelis Hatzis, Daniel Merk, Argyris Politis, Antreas Afantitis, Ioulia Tseti, Katerina M. Antoniou, Athol U. Wells, Vassilis Aidinis

## Abstract

Fibrosis is a significant mortality factor and health concern, promoting organ malfunction as well as immune and chemical resistance. Among the different fibroproliferative diseases, idiopathic pulmonary fibrosis (IPF) is a fatal fibrotic interstitial lung disease (ILD) with limited therapeutic options. Autotaxin (ATX), an established therapeutic target in IPF, is a secreted lysophospholipase D that catalyses the extracellular production of lysophosphatidic acid (LPA), a growth factor-like signalling phospholipid. The many pathologic effects of LPA in the lung include the suppression of peroxisome proliferator-activated receptor γ (PPARγ), a therapeutic target in metabolic disorders, which are frequent comorbidities of IPF associated with unfavourable prognosis. In this report, we introduce EL244, the first-in-class dual ATX inhibitor and PPARγ agonist, which is endowed with drug-like properties. Developed through chemoinformatic repositioning, innovative rational design, targeted synthesis and pharmacological characterization, EL244 exhibited favourable ADMET and PK/PD profiles. Remarkably, EL244 inhalation, which alleviates systemic toxicity concerns, decreased pulmonary LPA levels and related effects in pulmonary cells, and attenuated bleomycin (BLM)-induced pulmonary fibrosis, restoring respiratory functions. Therefore, EL244 emerges as a promising candidate for the inhaled treatment of IPF and ILDs.

## INTRODUCTION

Fibrosis, the excessive deposition of collagen and other components of extracellular matrix, is a pathologic feature of many fibroproliferative diseases in all organs, including cancer, accounting for ∼45% of global deaths ^1^. Pulmonary fibroproliferative disorders, collectively referred to as interstitial lung diseases (ILDs), encompass a complex array with diverse prognoses and clinical behaviours, including multisystem conditions, such as systemic sclerosis (SSc-ILD) and rheumatoid arthritis (RA-ILD)^2,3^. Idiopathic pulmonary fibrosis (IPF), the most common and fatal ILD, is a chronic, progressive disease of unknown cause, with a prognosis worse than many types of cancer ^4^. While the current standard of care treatments for IPF and some ILDs, pirfenidone and nintedanib, slow disease progression and reduce mortality in patients who can tolerate them ^5^, they often cause troublesome side effects and have not been shown to improve quality of life in pivotal trials ^6,7^. Therefore, the treatment of IPF/ILD remains an unmet medical need.

Autotaxin (ATX) is a secreted lysophospholipase D that catalyses the extracellular production of lysophosphatidic acid (LPA), a bioactive growth-factor-like phospholipid^8^. Increased ATX expression has been detected in many fibroproliferative diseases ^8^, including IPF ^9,10^. Its genetic deletion from epithelial cells and macrophages attenuated the development of bleomycin (BLM)-induced pulmonary fibrosis in mice ^9^, indicating a pathologic role for ATX in pulmonary fibrosis and providing the proof of principle for pharmacologic targeting ^10^. Accordingly, pharmacologic ATX inhibition attenuated pulmonary fibrosis in animal models ^9,11–13^, thus establishing ATX as a therapeutic target in IPF and spurring the development of different ATX inhibitors ^14,15^. Although clinical trials with early ATX inhibitors have been discontinued ^16,17^, ongoing clinical trials continue to explore the therapeutic potential of ATX inhibition, using compounds with better target engagement and improved physicochemical properties ^18,19^.

LPA activates its cognate GPCR receptors, widely expressed in most pulmonary cell types, to stimulate TGF-β activation, vascular leak and fibroblast accumulation ^20,21^. Moreover, LPA has been suggested to inactivate and/or reduce the transcription of the nuclear receptor peroxisome proliferator-activated receptor γ (PPARγ) ^22^, which regulates the expression of genes involved in lipid metabolism and glucose homeostasis ^23^. PPARγ agonists (Tro-, pio-, rosi-glitazones) have been used as a first-line medication for type II diabetes and dyslipidaemia, widespread comorbidities of IPF associated with an unfavourable prognosis ^24^. Genetic deletion of PPARγ exacerbated fibrotic responses in a murine model of sarcoidosis ^25^, and PPARγ agonists attenuated BLM-induced pulmonary fibrosis ^26^, suggesting a therapeutic benefit for PPARγ agonism in IPF.

In this report, we present EL244, a first-in-class type IV ATX inhibitor and PPARγ agonist, developed through the application of chemoinformatics, molecular modelling, rational design, and targeted synthesis.

## RESULTS

### Repurposing TGZ as an ATX inhibitor

The repurposing of existing drugs for novel therapeutic indications has attracted substantial interest due to its capacity to expedite drug development timelines and mitigate associated costs ^27^. In this context, and to potentially repurpose FDA-approved drugs as ATX inhibitors, molecular docking was employed to virtually screen the Prestwick chemical library, which contains 1520 off-patent drugs with known bioavailability and safety profiles in humans. The computational virtual screening was performed with the Enalos Asclepios KNIME drug discovery pipeline ^28^, using RxDock for High Throughput Virtual Screening (HTVS)^29^. HTVS (Fig. 1A) was based on the docking score and the binding mode of each compound in the active site of ATX, including orientation, interactions, and size-similarity with the crystallographic ATX inhibitor HA-155 (PDB ID: 2XRG), a boronic acid-based potent ATX inhibitor ^30^. HVTS steps include the preparation of 3D ligand and protein structures, docking cavity mapping, ligand virtual screening and selection of promising binders (Fig. 1A).

**Fig. 1.**
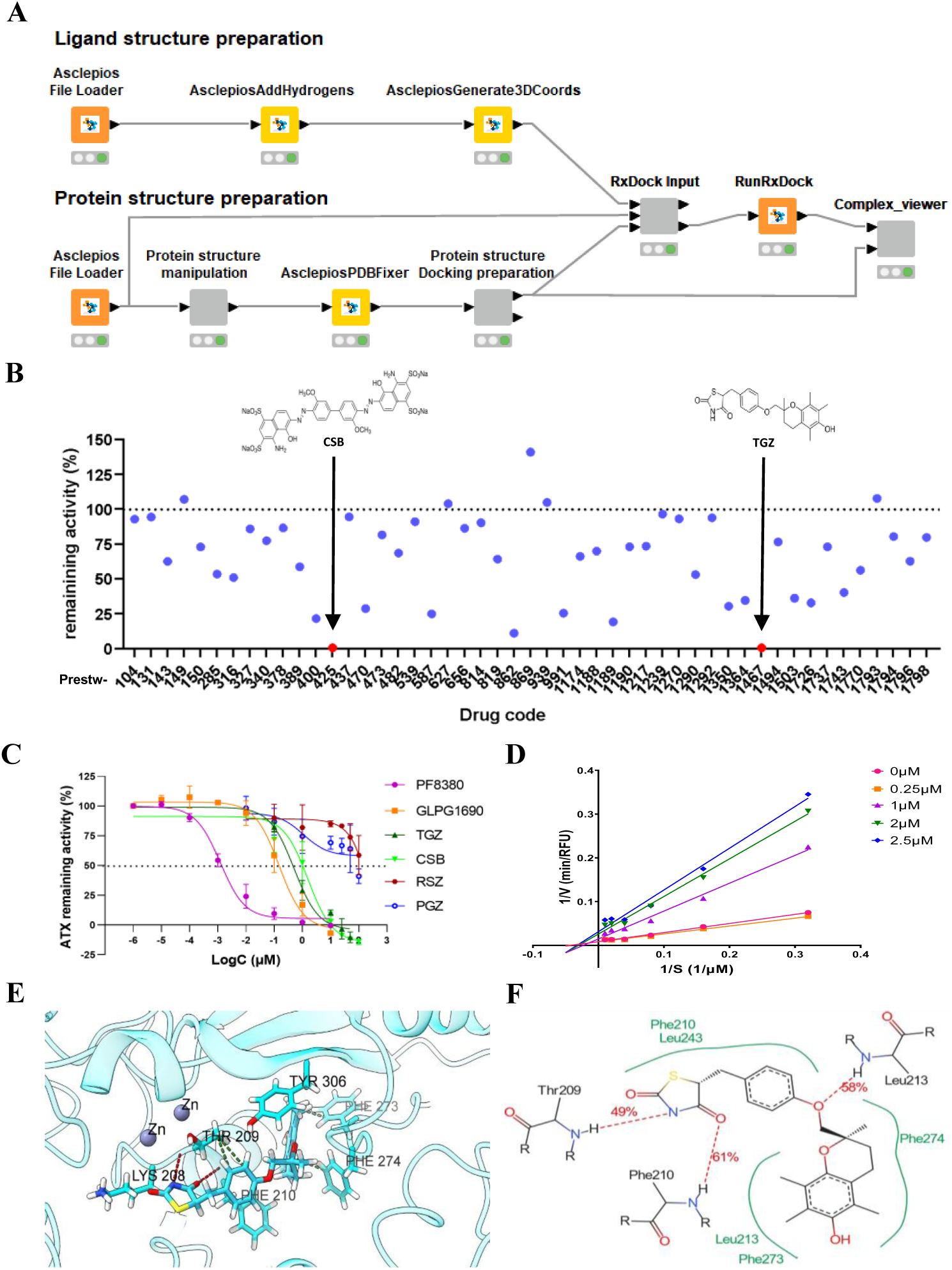
Repositioning TGZ as an ATX inhibitor. (**A**) Virtual screening workflow for the repurposing of FDA-approved drugs as potential ATX inhibitors. (**B**) *In vitro* screening of the *in silico* top-ranked compounds (at 50 μΜ) for their inhibitory activity against ATX. (**C**) Dose-response curves for the top “hits” identified, CSB and TGZ, in comparison with rosiglitazone (RGZ) and pioglitazone (PGZ), and the known ATX inhibitors PF-8380 and GLPG1690. (**D**) Mode of ATX inhibition by TGZ. (**E**) Three- and (**F**) two-dimensional MD simulation representations of TGZ_1, the most favourable TGZ isomer, in complex with ATX.

The top-ranked forty-nine compounds (Table S1) were subsequently tested *in vitro* for their ATX inhibitory activity using the Amplex-Red assay (Fig. 1B), as previously reported ^28,31,32^, and as recently published in detail ^33^. Surprisingly, the most potent ATX inhibitors were Chicago Sky Blue 6B (CSB, Prestw-425, Table S1), a diazo-dye with reported anti-inflammatory properties ^34^, and Troglitazone (TGZ, Prestw-1467, Table S1), a withdrawn first-line antidiabetic drug ^35^. Both compounds fully inhibited ATX activity at the maximum concentration tested (100 μΜ; Fig. 1B, Table S1).

Dose-response curves confirmed the significant inhibitory effect against ATX (IC_50_ values of 1.60 and 0.53 μΜ for CSB and TGZ, respectively; Fig. 1C), in comparison with PF-8380, the most potent *in vitro* ATX inhibitor reported ^36^, and GLPG1690, the first-in-class ATX inhibitor that entered clinical trials ^16,37^. Focusing on TGZ, mode of inhibition analysis with Lineweaver-Burk plots further revealed that TGZ is a non-competitive ATX inhibitor, suggesting that it does not bind to the catalytic site of the enzyme (Fig. 1D). In agreement, Molecular Dynamics (MD) simulations root mean square deviation (RMSD) analysis (Fig. S1) and Molecular Mechanics Generalised Born and Surface Area continuum solvation (MM/GBSA; Table S2)^38^, suggested that TGZ binds within the deep and hydrophobic pocket next to the catalytic site of ATX (Fig. 1E), in a conformation classified as a type-II inhibitor. Binding free energy calculations were performed with *RR* (TGZ_1) and *SS* (TGZ_4) TGZ diastereoisomers, exhibiting the best (and very similar) calculated total binding energies to ATX (ΔG*_bind_*, Table S3). In the first case (*RR*, TGZ_1), the lipophilic 5,7,8-trimethyl-benzopyran-6-ol segment is buried close to hydrophobic residues Phe273, Phe274, and Leu213, whose backbone amino-atoms form a hydrogen bond with the oxygen of anisole moiety, while the 2,4-thiazolidinedione group is participating in the formation of hydrogen bonds with the amino backbone of Thr209 and Phe210 (Fig. 1E), the latter of which is calculated to be the most favourable ATX amino acid towards binding (Table S2). In the other case, TGZ_4 (*SS*) positions its chromanol group near Phe274 (the most favourable residue towards binding; Table S2) and Leu213, forming a hydrogen bond. The oxygen of the anisole group is engaged in a hydrogen bond with the backbone amine of Trp275, while the 2,4-thiazolidinedione moiety forms a transient hydrogen bond with Arg244 (Fig. S2).

### Design, synthesis and pharmacological activity of the first-in-class dual ATX inhibitor/PPARγ agonist

TGZ is a synthetic ligand for PPARγ, the first in the thiazolidinedione (TZD) class of oral hypoglycaemic drugs ^23^. TGZ had been approved for the treatment of type II diabetes ^35^, but it was soon discontinued on account of hepatic toxicity ^39^. Several factors have been proposed for TGZ-induced hepatotoxicity, with the formation of reactive metabolites, especially the oxidation of the chromane moiety of the side chain of the drug to *o*-quinone methide and quinone, being the dominant one (Fig. 2)^40^. To limit hepatotoxicity, two other TZDs were developed, rosiglitazone (RGZ) and pioglitazone (PGZ), which also exhibit potent PPARγ agonism but have no hepatotoxicity ^41^. However, both RGZ and PGZ were found not to inhibit ATX enzymatic activity (Table 1, Fig. 1C). The *in vitro* experimental findings, together with the cheminformatic results, which suggest the accommodation of the chromane substituent of TGZ in the hydrophobic pocket of ATX (Figs. 1E,F, S2), highlight the key role of this lipophilic moiety in endowing TGZ with ATX inhibitory activity. Accordingly, the lack of potency by both RGZ and PGZ against ATX could be attributed to their polar, compared to TGZ, tail (2-methylpyridine and 5-ethylpyridine, respectively), rendering them non-tolerated in the hydrophobic pocket of ATX.

**Figure 2.**
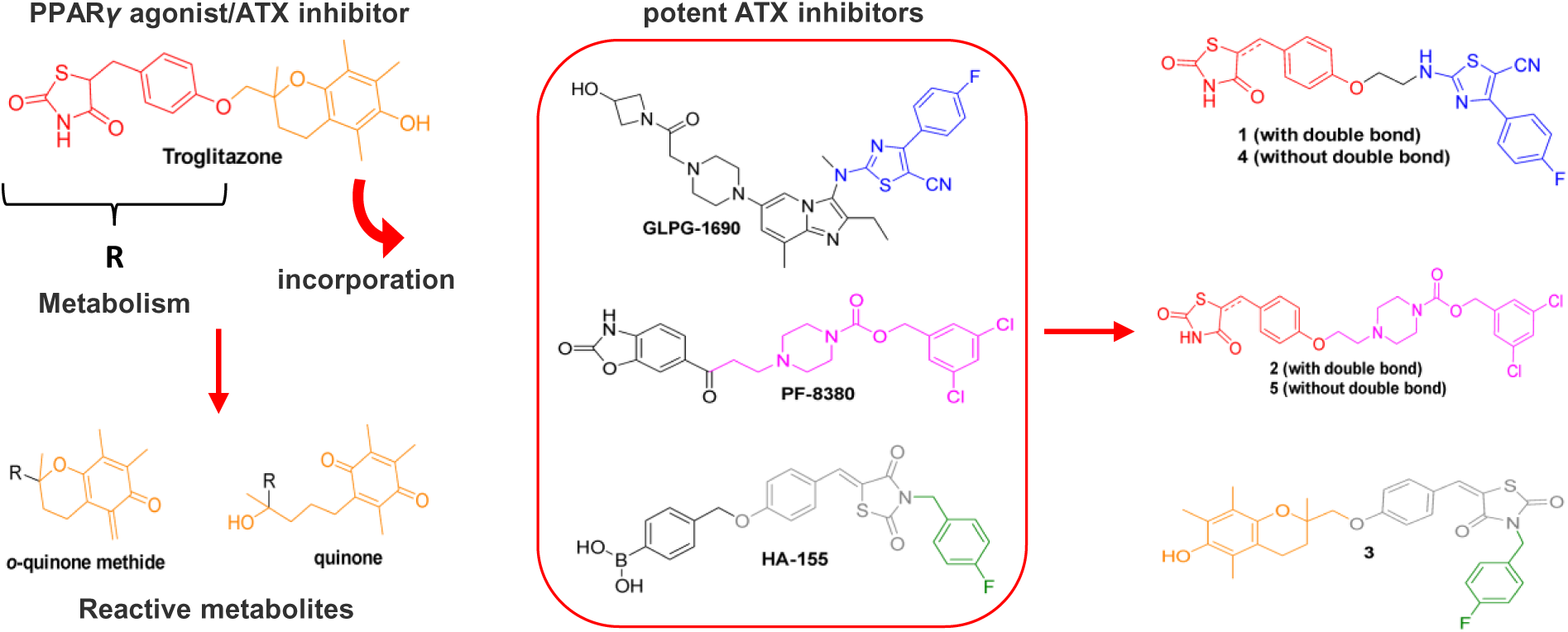
Design of TGZ derivatives 1-5. Incorporation of structural moieties of TGZ and three chemically diverse ATX inhibitors (GLPG1690, PF-8380 and HA-15) in one structure. In 1, 2, 4 and 5, the metabolically labile chromane moiety of TGZ (in orange), which is considered responsible for TGZ-induced hepatotoxicity, was replaced by the lipophilic substituents of GLPG1690 (in blue) and PF-8380 (in purple). The lipophilic moieties of the reference compounds highlighted in blue (GLPG1690), purple (PF-8380) and green (HA-155), bind to the hydrophobic pocket of ATX according to X-ray crystallography studies.

**Table 1.**
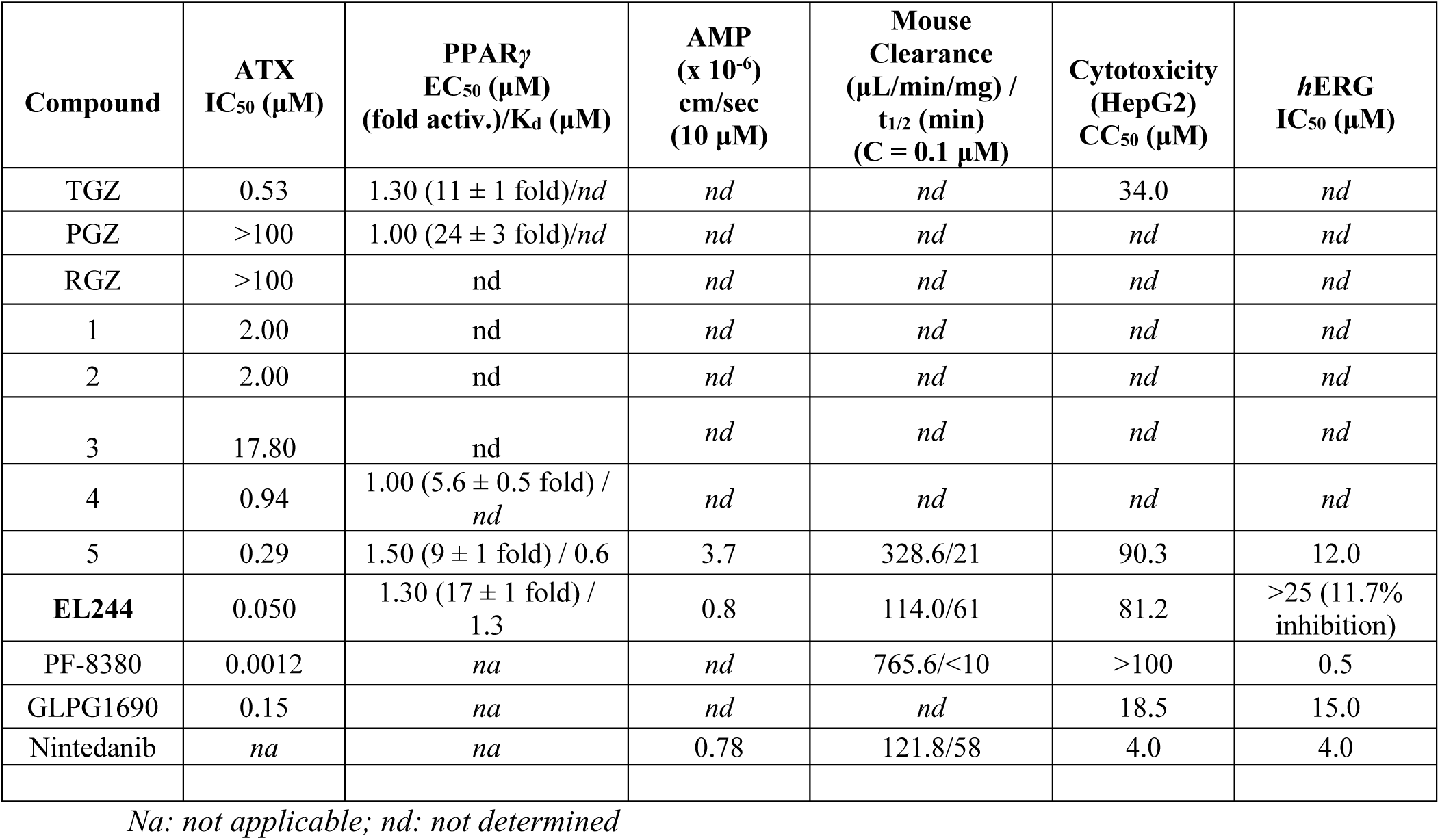
*In vitro* and “drug-like” profile of compounds 1-5, EL244 and references.

To improve the ATX inhibitory activity of TGZ and its drug-like features, while maintaining its PPARγ agonism which offers additional therapeutic benefits, three new molecules were initially designed (1-3, Fig. 2) and synthesised by incorporating structural features of TGZ and known potent ATX inhibitors (GLPG1690, PF-8380 and HA-155) in one structure (Fig. 2). Specifically, in compounds 1 and 2 the metabolically labile 5,7,8-trimethyl-chroman-6-ol group of TGZ (in orange, Fig. 2), which according to MD simulations appears to bind to the hydrophobic pocket of ATX (Fig. 1E,F), was replaced by the respective lipophilic moieties of GLPG1690 (in blue, Fig. 2) and PF-8380 (in purple, Fig. 2), respectively, which follow a similar binding pattern ^11,37^. In contrast, compound 3, derived from the substitution of the (4-(hydroxymethyl)phenyl) boronic acid group in HA-155 ^42,43^, which interacts with the catalytic site of ATX, with 2-(hydroxymethyl)-2,5,7,8-tetramethylchroman-6-ol of TGZ (Fig. 2). Furthermore, the reduced analogues of 1 and 2, compounds 4 and 5, respectively, were synthesised, aiming at exploring the impact of rigidity/flexibility on activity (Fig. 2). The synthetic procedures followed for the synthesis of 1, 2, 4 and 5 are displayed in Figure 3, while compound 3 was synthesised according to the route depicted in Figure S3 (the synthetic methodology is described in detail in the Supplementary Materials and Methods).

**Fig. 3.**
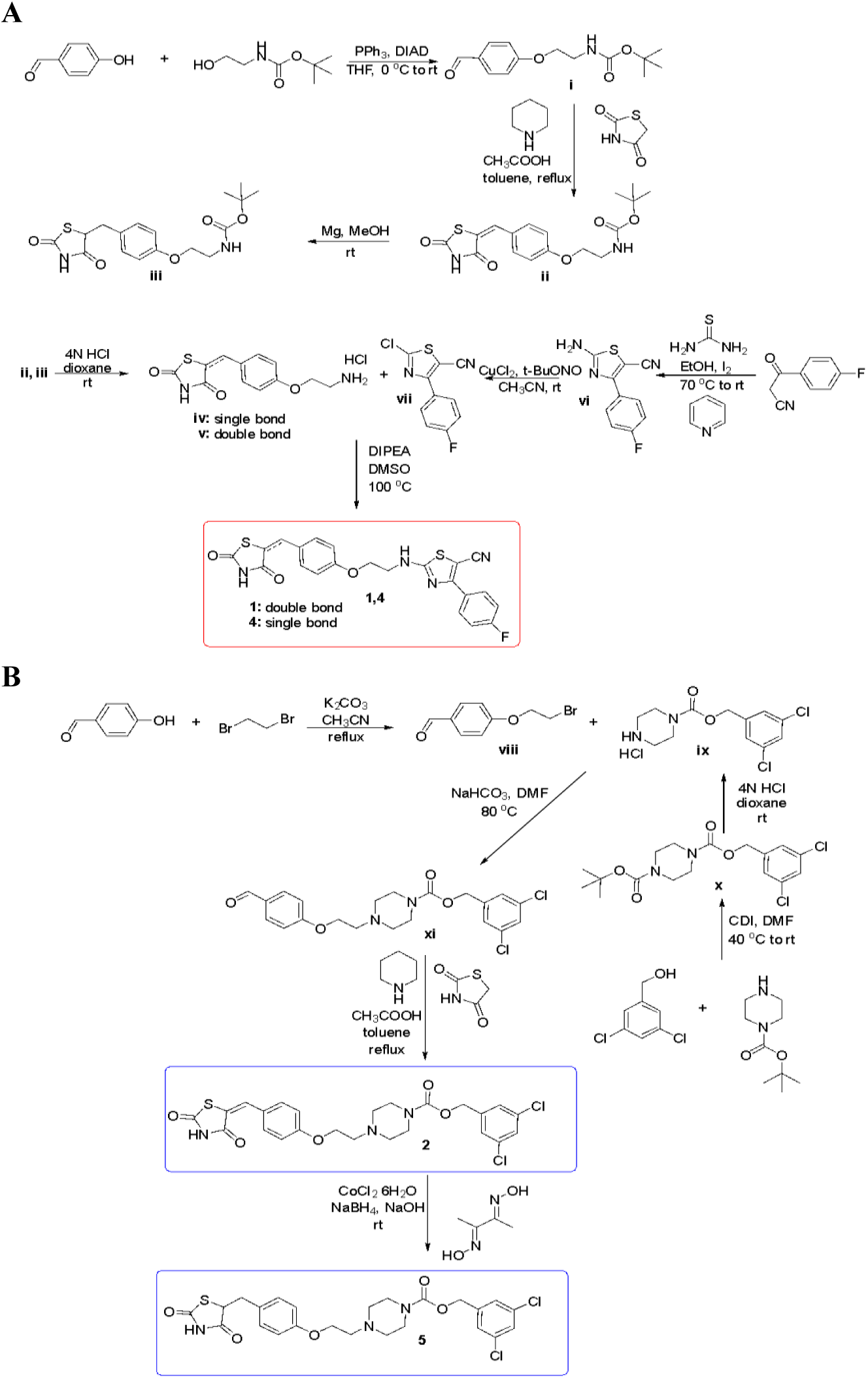
**Synthetic routes for compounds 1, 2, 4 and 5.**

Following synthesis, the ATX inhibitory activity of compounds 1-3 was tested *in vitro* (Table 1, Fig. 4A). Both 1 and 2 exhibited the same activity against ATX (IC_50_ values of 2.00 μΜ), although 4-fold lower than that of TGZ (IC_50_ = 0.53 μΜ), while the inhibition offered by compound 3 was found to be very weak (IC_50_ = 17.80 μΜ). Of note, the reduction of the double bond of 1 and 2 afforded derivatives exhibiting an ATX inhibitory activity similar to TGZ (compound 4, IC_50_ = 0.94 μΜ, Table 1) or equipotent to GLPG1690 (compound 5, IC_50_ = 0.29 μΜ, Table 1, Fig. 4B, Fig. S4A). Minimal interference of compounds 4 and 5 on the 2nd and 3rd steps of the Amplex Red assay was observed (Fig. S4B), further supporting their ATX inhibitory properties. Mode of inhibition analysis with Lineweaver-Burk plots revealed that compound 5 can bind to both the free enzyme and the enzyme-substrate complex (Fig. 4C), acting as a non-competitive ATX inhibitor. Moreover, compound 5 was found, with the TOOS assay ^44^, to also inhibit ATX activity in serum, exhibiting a dose-dependent effect upon incubation with a high concentration of exogenous LPC (2 mM)(Fig. 4D).

**Fig. 4.**
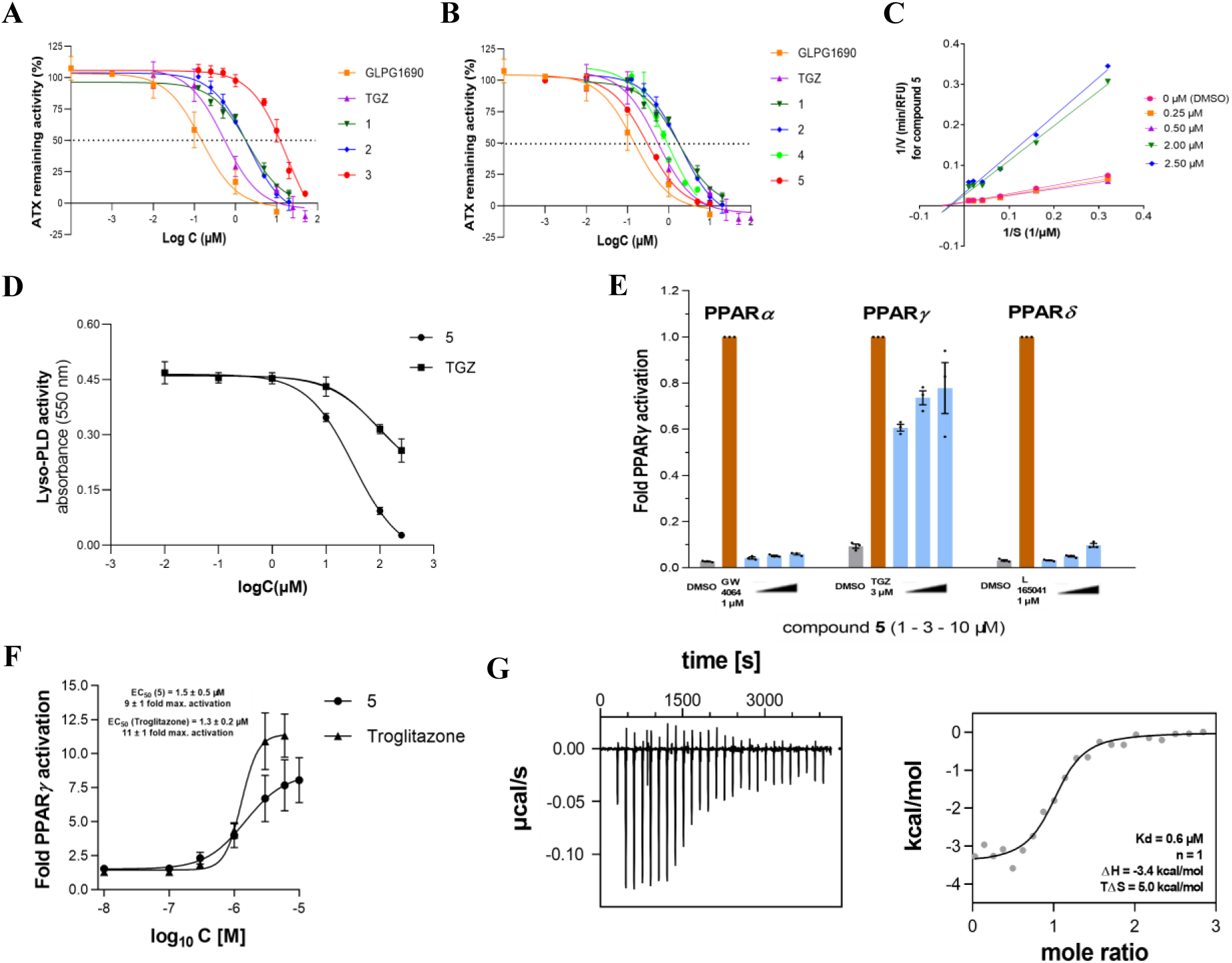
Compound 5 is a dual ATX inhibitor/PPARγ agonist. (**A**) Dose-response curves for GLPG1690, TGZ, and 1-3. (**B**) Dose response curves for 1, 2, 4 and 5. GLPG1690 and TGZ were used as references. (**C**) Mode of inhibition of compound 5. (**D**) Dose-response curves of lyso-PLD inhibitory activity by compound 5 and TGZ in serum and in the presence of high (exogenous) LPC concentration (2 mM; TOOS assay). (**E**) Activity of 5 against PPAR*_a_*, PPAR*_γ_* and PPAR*_δ_*. (**F**) Dose-response curves for the activation of PPAR*_γ_* by TGZ and compound 5. (**G**) Binding of 5 to the PPAR*_γ_* LBD was confirmed by ITC with a K_d_ value of 0.6 μM. The left panel displays the isotherm of the **5**-protein titration; while the right panel shows the fitting of the heat of binding.

Compound 5 was also tested against PPARγ, as well as its variants PPARα and PPARδ, using Gal4-hybrid receptor plasmids (pFA-CMV-hPPARα/γ/δ-LBD) transfected in HEK293 cells. The reporter system tests compound binding to the ligand binding domain (LBD) of the canonical isoform of the respective human PPARs ^45^. Compound 5 was found to be a selective PPARγ agonist (Table 1; Fig. 4E,F; EC_50_ 1.5 μΜ), being comparable to TGZ (EC_50_ = 1.3 μΜ), while it also exhibited a high binding affinity to the PPARγ ligand binding domain (LBD) in Isothermal Titration Calorimetry (ITC)(K_d_ = 0.6 μΜ; Table 1, Fig. 4G), indicating that compound 5 retains its PPARγ agonist properties.

### Compound 5 exhibits a promising ADMET and PK profile

Compound 5, exhibiting the most balanced ATX inhibitor/PPARγ agonist profile, was then evaluated *in vitro* in terms of representative ADMET properties potentially affecting the *in vivo* pharmacokinetic (PK) profile of a bioactive molecule (Table 1). It exhibited increased cell membrane permeability and a moderate metabolic clearance, without displaying significant cytotoxicity (EC_50_ = 90.3 μΜ, Table 1). In addition, a weak inhibitory activity against the cardiac potassium ion channel *h*ERG (*human* Ether-à-go-go-Related Gene) was exhibited (IC_50_ = 12 μΜ), which was much lower than that of PF-8380 (IC_50_ = 0.5 μΜ), was similar to that of GLPG1690 (IC_50_ = 15 μΜ), and was lower than that of the SOC nintedanib (IC_50_ = 4 μΜ)(Table 1).

Fast track PK analysis of compound 5, and specifically its plasma exposure, was measured after different routes of administration (*iv*, *ip*, *inhaled* and *per os*). The plasma concentrations of compound 5 after *ip* and *per os* administration, and both at 1 and 3h post-dose, were comparable to the respective *iv* concentrations used as reference (100% absorption)(Fig. S5), indicating favourable compound absorption into the circulation. Notably, the direct administration of compound 5 in the lung through inhalation demonstrated its prolonged exposure and retention in the tissue, with no detectable amounts of the compound in the circulation even three hours post-dose (Fig. S5), suggesting targeted delivery of compound 5 to the respiratory tract.

### Compound 5 attenuates BLM-induced pulmonary fibrosis

Given the established role of ATX in the pathogenesis of pulmonary fibrosis ^9,10^ and the suggested involvement of PPARγ ^26,46^, the efficacy of compound 5 to inhibit BLM-induced pulmonary fibrosis was then tested.

Compound 5 was first tested on mouse precision-cut lung slices (PCLS), living lung slices isolated from BLM-induced fibrotic mouse lung tissue (Fig. S6A), as recently published in detail ^47^. PCLS, a bridge between traditional *in vitro* cell cultures and *in vivo* animal models, have emerged as a valuable pre-clinical platform to test new pharmacological compounds ^48^. Compound 5, at 30 μΜ and after 72h of incubation, improved the lung architecture compared to the BLM group, with much fewer fibrotic lesions being observed (Fig. S6B,C), decreasing at the same time the *m*RNA expression levels of the profibrotic gene expression markers collagen1α1 (*Col1a1*) and fibronectin 1 (*Fn1*)(Fig. S6D,E).

Compound 5 was then evaluated in the BLM-induced pulmonary fibrosis model, the most widely used animal model for pulmonary fibrosis ^49,50^. BLM (0.8 U/Kg) was administered via oropharyngeal administration (OA) to littermate C57Bl/6 mice, as previously described ^50,51^, as analysed in detail (<u>protocols.io</u>), and as summarised graphically (Fig. S7A). Given the favourable PK profile upon inhalation (Fig. S5), the therapeutic potential of compound 5 was evaluated following its inhaled administration (15 mg/Kg) twice daily (b.i.d.) in a therapeutic mode, starting the administration 7 days post BLM (Fig. S7A). The aerosolised delivery of compound 5 efficiently attenuated the BLM-induced impairment of respiratory functions, as indicated in mean static lung compliance (Cst), mean respiratory system compliance (Crs) and total mean lung capacity (A) values (Fig. S7B). Accordingly, compound 5 reduced vascular leak and pulmonary oedema as indicated by the decreased total protein levels in the bronchoalveolar lavage fluid (BALF)(Fig. S7C) and decreased inflammatory cells in the BALF (Fig. S7D). Although the decrease of BALF soluble collagen (Fig. S7E) and lung tissue *Col1a1* mRNA levels (Fig. S7F) did not reach statistical significance following compound 5 administration, histological analysis revealed fewer fibrotic regions and decreased collagen deposition as evaluated with H&E and Fast green Sirius red staining (Fig. S7G,H). Therefore, the favourable PK profile and the promising efficacy of compound 5 suggest its further pre-clinical development and optimisation.

### Development of EL244, an optimised dual ATX inhibitor/PPARγ agonist

Based on the ADMET profile of the molecule (Table 1), we focused on its *h*ERG inhibition, which, although moderate to weak, could lead to potential side effects related to cardiovascular toxicity. Importantly, the *h*ERG toxicity evaluation early in the preclinical setting is strongly recommended by FDA and EMA, while cardiovascular assessment constitutes a pivotal point for obtaining investigational new drug (IND) status ^52^. Accordingly, to avoid *h*ERG interactions, the piperazine group of compound 5 was replaced by the piperidine one (Fig. 5A), on the grounds that highly basic (ionizable) amine motifs are well-recognized and accommodated in the negative electrostatic potential located in *h*ERG’s central cavity ^53,54^. It should be noted that the most potent ATX inhibitor currently available, PF-8380 (Fig. 2), has been found to significantly inhibit *h*ERG (Table 1), an effect attributed to the basic piperazine group ^55^. Furthermore, the former clinical candidate GLPG1690 (Fig. 2), exhibiting a moderate to weak inhibitory activity against *h*ERG (Table 1)^11^ originated from reducing the basicity of a precursor molecule of the same series, which exerts a much more potent *h*ERG inhibition ^11,37^.

**Fig. 5.**
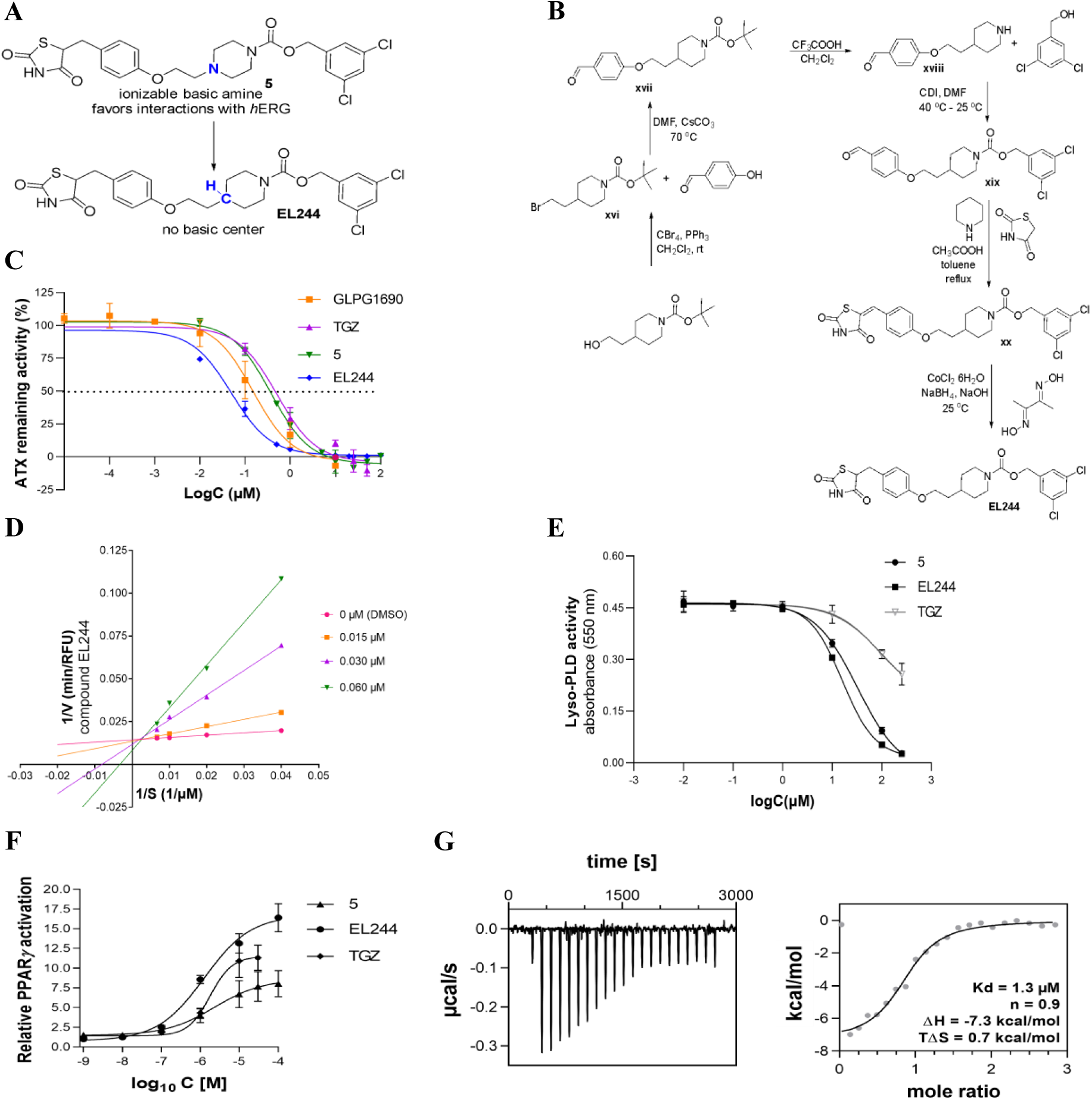
Synthesis and i*n vitro* and *ex vivo* activity of EL244. (**A**) Targeted design towards decreasing hERG interactions. (**B**) Synthetic procedure followed for the synthesis of EL244. (**C**) Dose-response curves for GLPG1690, TGZ, 5 and EL244 against ATX *in vitro* (Amplex-Red assay). (**D**) Mode of inhibition of compound EL244. (**E**) *Ex vivo* (lyso-PLD activity assay), in serum and in the presence of 2 mM exogenous LPC for TGZ, 5, and EL244. (**F**) Dose-response curves for the activation of PPARγ by TGZ, 5, and EL244. (**G**) Binding of EL244 to the PPARγ LBD as confirmed by ITC. The left panel shows the isotherm of the ΕL244-protein titration; the right panel shows the fitting of the heat of binding.

The synthetic route followed to produce compound EL244 is illustrated in Figure 5B, while the synthetic methodology is described in detail in the Supplementary Materials and Methods. Remarkably, this minor structural modification implemented on compound 5 proved advantageous in reducing its overall *h*ERG binding affinity, since EL244 inhibited the cardiac potassium ion channel in a very weak fashion (approximately 12% at 25 μΜ, Table 1). Furthermore, the cytotoxicity caused by EL244 in human HepG2 cells was found to be very weak (EC_50_ = 81.2 μΜ, Table 1), similar to that of compound 5, and much better compared to references TGZ, GLPG1690, and nintedanib (Table 1), indicating low liver toxicity. Notably, EL244 exhibited a much lower metabolic clearance rate compared to that of compound 5 (Table 1), thus identifying the tertiary amine of the piperazinyl group of compound 5 as a major metabolic “soft spot”. Importantly, this modification conferred a 5-, 10- and 3-fold better ATX inhibitory effect *in vitro* compared to that of compound 5, TGZ and GLPG1690, respectively (Table 1, Fig. 5C). Mode of inhibition analysis (Fig. 5D) indicated a competitive inhibition for EL244, providing evidence that the compound binds only to the free enzyme, and in such a way as to compete with LPC binding to ATX. Improved lyso-PLD inhibition *ex vivo* for EL244 was also observed (Fig. 5E).

EL244 activated PPARγ more efficiently than its precursors compound 5 and TGZ, and similar to PGZ, at the same EC_50_ concentration (Table 1, Fig. 5F). Of note, it can be inferred by the ITC studies performed that although the binding of both compound 5 and EL244 to the LBD of PPARγ is exothermic, with their ΔG and Kd values being close, each compound exhibits different thermodynamics (Figs. 4G and 5G). In particular, the protonated piperazinyl-carbamate side chain in compound 5 seems to favour an entropy-driven binding (Fig. 4G), while EL244, having the neutral piperidinyl-carbamate side chain, binds in an enthalpy-driven manner to the PPARγ-LBD (Fig. 5G).

Fast track PK analysis of EL244 upon *ip* administration (30 mg/Kg) indicated that the compound reaches a 40 μM concentration in the plasma in one hour, its plasma levels are retained up to 3h (∼35 μΜ), while it can still be detected at lower concentrations (15 μM) even 9 hours post-dose (Fig. S8A). Pharmacodynamic (PD) analysis of EL244, as measured by the reduction of plasma LPA levels using MS/MS, revealed a maximal effect at 3 hours post-administration (Fig. S8A). Interestingly, PK analysis of EL244 after inhaled administration (15 mg/Kg) indicated that the compound is highly retained in the lung, while its clearance rate from the lung and its absorption in the systemic circulation were found to be low (Fig. S8B), suggesting targeted delivery of EL244 directly to the lung.

### HDX-MS and MD simulations identify EL244 as a type IV ATX inhibitor

To gain structural insights into the binding mode of EL244 to ATX, we employed hydrogen-deuterium exchange mass spectrometry (HDX-MS) in conjunction with MD simulations. Optimised digestion conditions resulted in 310 peptides covering 84.5% of the ATX sequence with a redundancy of 5.62 (Table S4; Fig. 6A). Statistical analysis (α = 0.01) using a hybrid significant test of the differential HDX-MS data ^56^(Fig. S9A) identified several overlapping peptides (242-259, 276-289 and 214-231) as protected at different time points (0.5, 5 and 50 min) when bound to EL244 (Fig. S9B), covering the region 214-289. Mapping protected ATX fragments by EL244 into the crystal structure of ATX (2XR9)^57^(Fig. 6B) revealed that EL244 binds at the hydrophobic pocket and the allosteric tunnel of ATX ^15^, thus classifying EL244 as a type IV ATX inhibitor.

**Fig. 6.**
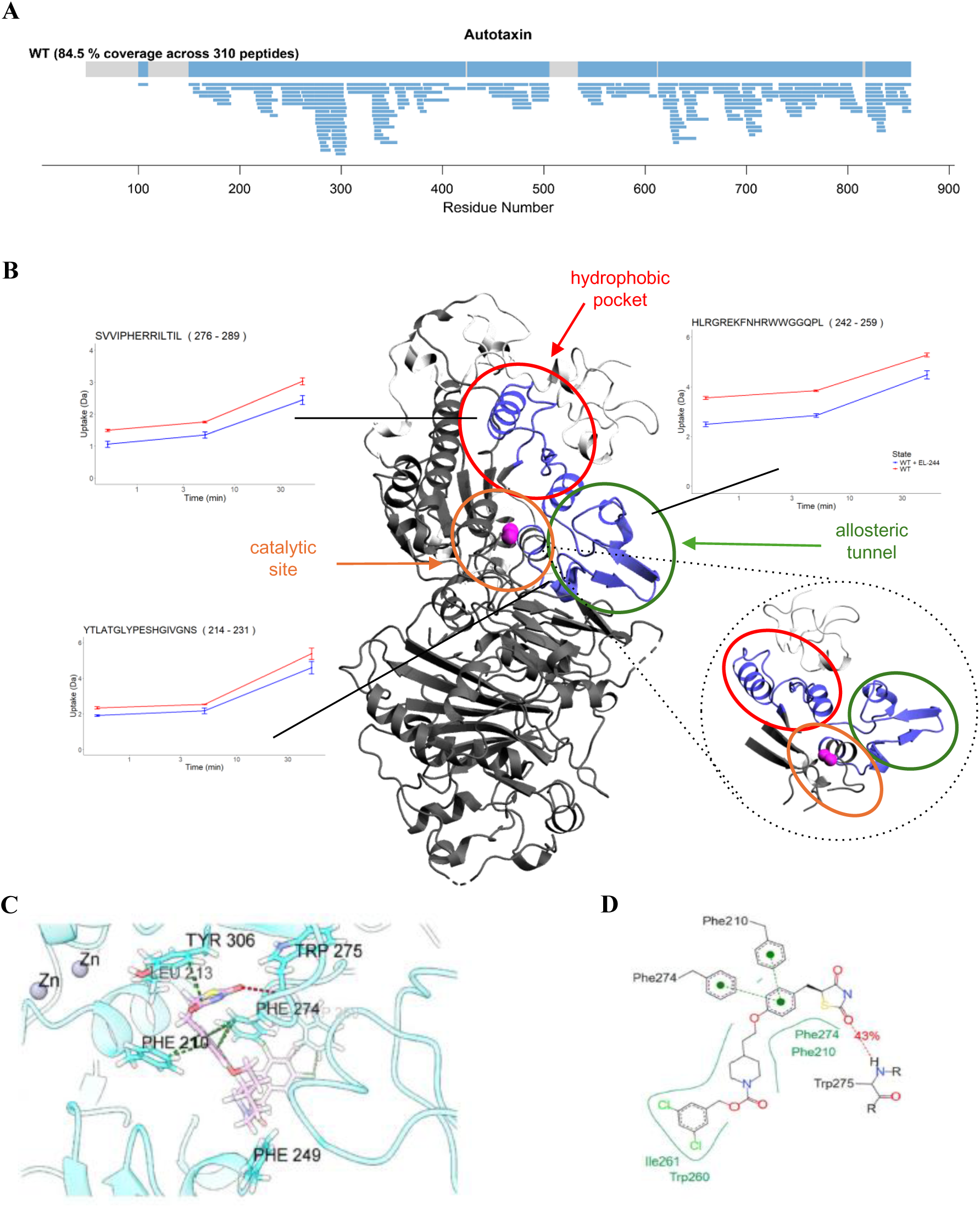
HDX-MS and MD simulations identify EL244 as a type IV ATX inhibitor. (**A**) ATX peptic peptide coverage map. (**B**) HDX-MS significant differences after binding with EL244 mapped onto the ATX crystal structure (2XR9) and representative uptake plots of the differences. Blue represents regions with increased protection upon binding, grey regions indicate no differences, and white, regions with no coverage. Purple spheres correspond to zinc ions in the crystal structure (catalytic cofactors). Close-up region shows the proposed binding region identified by HDX-MS. The three binding sites of ATX are displayed with circles, the region protected by both analogues is displayed with blue. (**C**) Three- and (**D**) two-dimensional MD simulation representations of isomer TGZ_1 in complex with ATX.

In alignment with the HDX-MS results, MD simulations with R-(EL244_1) and *S*-enantiomers (EL244_2) of EL244 indicated that only the binding of *S*-enantiomers is compatible with a type IV inhibition (Table S3; Fig. 6C,D). Accordingly, EL244_2 maintains a Type IV inhibitor binding mode, with its thiazolidine-2,4-dione moiety shifted towards the allosteric tunnel, forming a hydrogen bond with the backbone amino group of Trp275 (Fig. 6C,D), mediated by water for approximately 53% of the simulation (Table S5). The anisole fragment is oriented towards Phe274, calculated as the most favourable residue (Table S2), and Phe210, while the dichlorobenzene group extends further inside the allosteric hydrophobic tunnel, towards Trp260 and Ile261 (Fig. 6C,D).

MD analysis of EL244 isomers with PPARγ indicated that all complexes are structurally stable (Fig. S10A, Table S6), while MD analysis indicated that Cys285 is central in the binding of EL244 isomers to the LBD of PPARγ (Fig. S10B-E). The *S*-enantiomer of EL244 (EL244_2; Table S3) is characterised by favourable interactions and a hydrogen bond network formed between the 2,4-thiazolidinedione pharmacophore and Ser289, His323, His449 and Tyr473 residues (Fig. S10B,C; Table S6), contrary to its *R*-counterparts EL244_1 (Figure S10D,E; Tables S3 and S6), thus highlighting the importance of chirality in ligand recognition and protein function ^58^. It is worth mentioning that the latter hydrogen bond network is totally preserved in the active conformations (*S*-enantiomers) of the reference compounds (PGZ_2, RGZ_2, and TGZ_3, Table S3) within PPARγ-LBD (Figure S11, Table S6), thus representing the archetypal engagement of effective PPARγ agonists, which is preserved in EL244, further supporting efficient PPAR agonism by EL244.

### EL244 attenuates BLM-induced pulmonary fibrosis

Given the PK profile of EL244 (Fig. S8) and the therapeutic benefits of inhaled administration, the efficacy of EL244 was evaluated upon its inhaled administration in the BLM-induced model of pulmonary fibrosis ^49,50,59^, in prophylactic and therapeutic modes (15 mg/Kg; once daily/o.d.).

In the prophylactic mode, EL244 was administered for 15 consecutive days, starting one day before BLM administration (Fig. S12A). EL244 significantly improved respiratory functions (Fig. S12B) and markedly reduced BLM-induced pulmonary oedema and inflammation (Fig. S12C,D). Soluble collagen levels in BALF and *Col1a1* mRNA expression in lung tissue were also significantly reduced (Fig. S12E,F). Histological analysis confirmed the attenuated collagen deposition and the reduction of fibrotic regions (Fig. S12G,H). The two weeks of inhaled administration of EL244 did not cause any appreciable toxicity in the liver, as indicated by the ALT/AST levels (Fig. S12I), and as expected by the limited absorption of EL244 in the systemic circulation (Fig. S8B).

In the therapeutic mode, EL244 was administered for seven consecutive days starting on day 7 post-BLM administration, where inflammation starts to diminish, and fibrotic regions began to appear (Fig. 7A)^50,60^. EL244 significantly restored all assessed respiratory functions (Fig. 7B) and reduced vascular permeability (Fig. 7C). The EL244 effects on inflammation were minor, in line with the suggested limited role of ATX in acute inflammation in the lung ^61^. Importantly, EL244 demonstrated strong anti-fibrotic effects, significantly reducing soluble collagen in BALF and *Col1a1* expression in lung tissue (Fig. 7E,F). Histopathological analysis revealed reduced collagen deposition and fewer and smaller fibrotic regions (Fig. 7G,H).

**Fig. 7.**
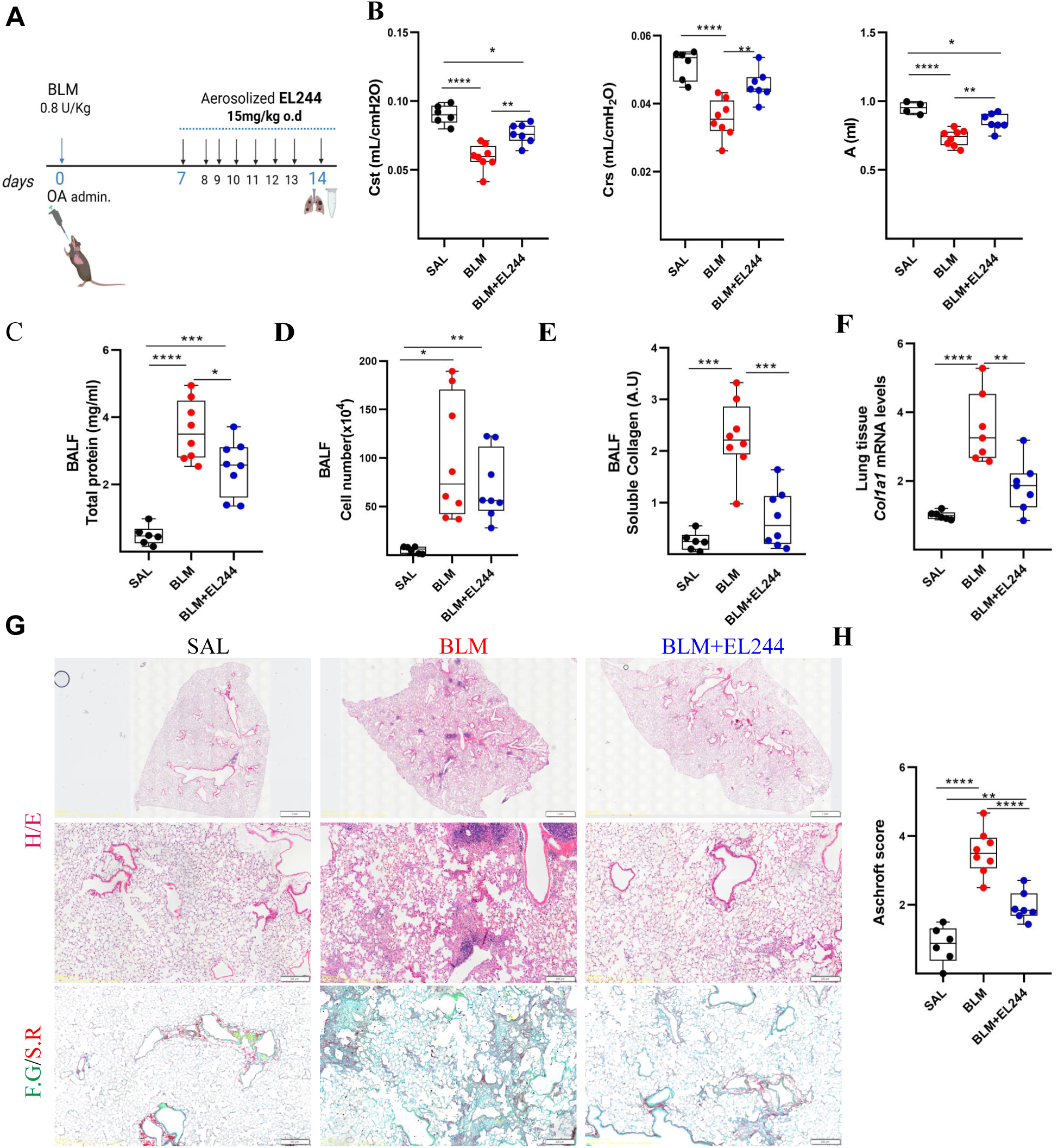
Inhaled therapeutic EL244 administration attenuates BLM-induced pulmonary fibrosis. (**A**) Sche-matic representation of drug administration. (**B**) Respiratory mechanics assessed using the FlexiVent system. (**C**) Total protein concentration in BALFs, as determined with the Bradford assay. (**D**) Inflammatory cell num-bers in BALFs, as counted with a haematocytometer. (**E**) Soluble collagen levels in the BALFs were detected with the direct red assay. (**F**) *Collal* mRNA expression was interrogated with Q-RT-PCR; values were normal-ized to the expression of *B2m* and presented as fold change over control. (**G**) Representative images from lung sections of murine lungs of the indicated genotypes, stained with H&E and Fast Green/Sirius Red (F.G/S.R; green/red). (**H**) Quantification of fibrosis severity in H&E-stained lung sections via the Ashcroft score. Follow-ing normality testing, statistical significance was assessed with one-way ANOVA and post-hoc Tukey’s test (B,C,E,F) or Welch ANOVA and post-hoc Games-Howell test (D,H).

### EL244 mode of action

To verify target engagement by EL244 and the diminished production of LPA, the enzymatic product of ATX mediating its pathological effects, LPA levels were measured using MS/MS in the plasma and BALF of mice after BLM administration and inhaled administration of EL244. All LPA species, except for 18:0, were found to be reduced in the BALF, including the most abundant species 16:0 and 18:2 (Fig. 8A). As a result, total LPA content in the BALFs was found reduced (Fig. 8B). Consistent with the high absorption of EL244 in the lung (Fig. S8B), no modulation of plasma LPA levels was detected (Fig. 8C,D), indicating that the effects of EL244 are localised in the lung.

**Fig. 8.**
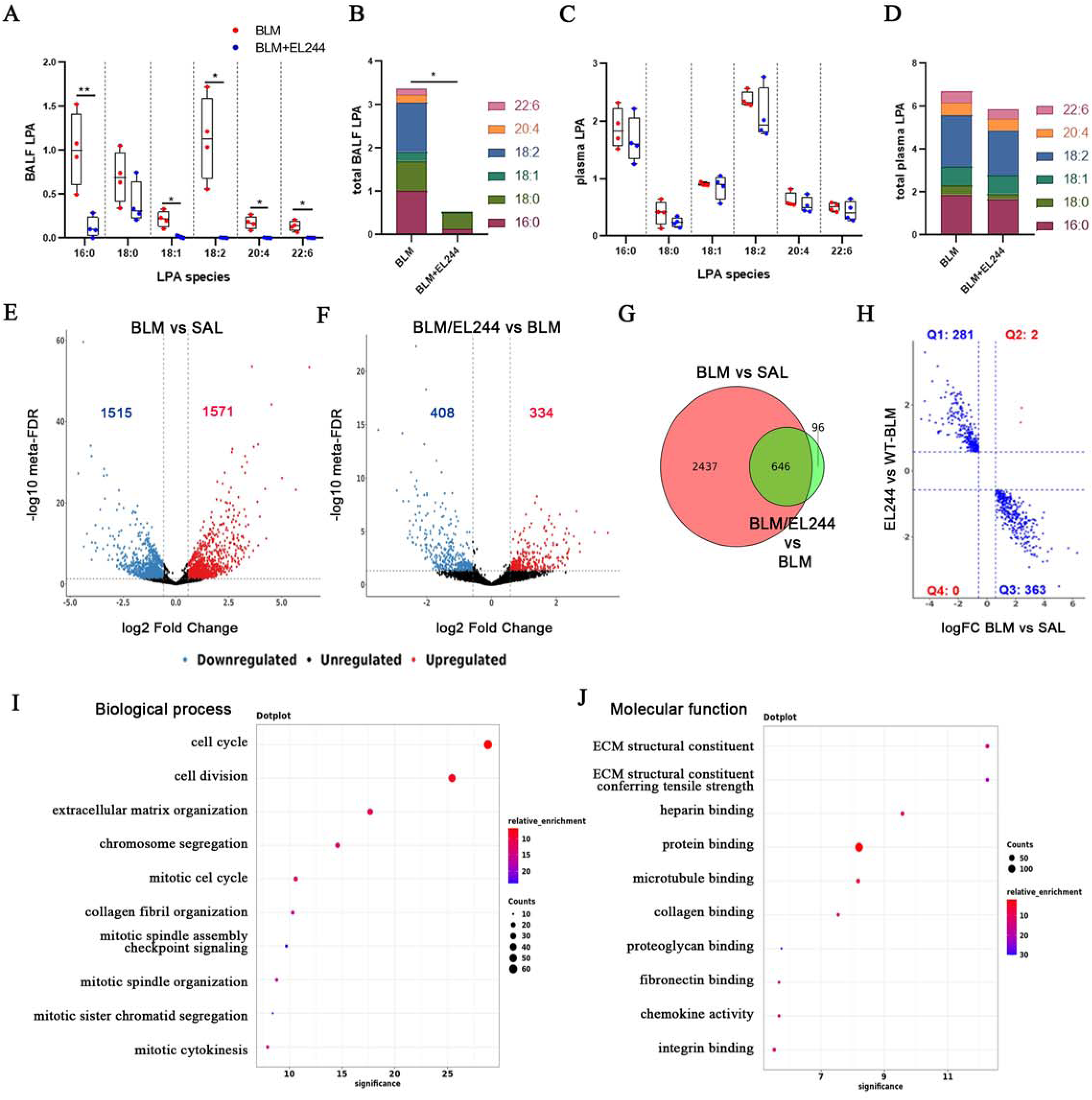
Mode of action of inhaled EL244 in BLM-induced pulmonary fibrosis. (**A**) LPA levels were measured with MS/MS. (**B**) Total LPA levels in BALFs. (**C**) LPA levels in plasma were measured with MS/MS. (**D**) Total LPA levels in plasma. (**E**) Volcano plot depicting the differentially expressed genes (DEGs) upon BLM treatment (compared to SAL). (**F**) Volcano plot depicting the DEGs upon EL244 treatment of BLM-treated mice (compared to BLM).(**G**) Venn diagram comparing the lists of DEGs following BLM (in red) and EL244 (in green) treatment. **(H)** Scatter plot showing the regulation patterns of the 646 common DEGs across the four indicated quadrants. **(1-J).** Dot plots showing the top 10 statistically enriched GO tem1s associated with the DEGs in Q3 (downregulated by EL244 and upregulated by BLM).Following normality testing, statistical significance was assessed with an unpaired t-test separately for each LPA species **(A-D).**

ATX pathologic effects in the lung are primarily attributed to the production of LPA ^8^, localised at the cell surface via ATX-integrin binding ^57^. LPA engages the adjacent LPA receptors (LPAR1-2), which activate the respective G-proteins and stimulate numerous universal and cell-specific cellular transduction pathways in pulmonary cells ^10,21^. To dissect the therapeutic effects of EL244 and the resulting diminished LPA production, the gene expression profiles of lung tissues post-BLM were analysed using RNA sequencing. BLM-induced pulmonary fibrosis was found to deregulate the expression of 3086 genes in the lungs (Differentially Expressed Genes/DEGs) in comparison with saline-treated mice (SAL)(Fig. 8E; Table S7), which include many well-known profibrotic DEGs (such as, most notably: *Col1a1, Cthrc1, Eln, Fn1, Spp1, Trem2, Tnc, Saa3;* Fig. 8E). Inhaled administration of EL244 in BLM-treated mice resulted in the deregulation of only 741 genes in comparison with the BLM-treated mice, including the downregulation of many of the identified pro-fibrotic BLM-DEGs (Fig. 8F; Table S8).

Remarkably, most EL244-deregulated DEGs (646) are a subset of BLM-induced DEGs (Fig. 8G). Among these, 363 BLM-induced DEGs were suppressed by EL244 treatment (Q3), Table S9; Fig. 8H). Gene ontology (GO) analysis of the biological processes (BP) of Q3 DEGs indicated that EL244 administration resulted in the down regulation of genes involved in cell cycle and mitosis (Fig. 8I), consistent with the well-known mitogenic properties of LPA, whose production was suppressed by EL244. Respectively, the most statistically enriched molecular function (MF) of these DEGs concerned the ECM (Fig. 8J), in agreement with the observed attenuation of BLM-induced pulmonary fibrosis by EL244.

## DISCUSSION

The establishment of ATX as a therapeutic target in IPF has led to the ISABELA clinical trials with GLPG1690, among the largest IPF clinical studies of the last decade. However, the trials were discontinued due to a low risk-to-benefit ratio ^16,17^. Moreover, the COVID-19 pandemic affected patient participation and compliance, and unexpectedly stimulated mortality in the GLPG1690 group ^16^. However, the complex effects of ATX/LPA on immune regulation ^62^, especially amid the cytokine storm in COVID-19 ^63^, remain unresolved. More importantly, the inhaled administration of ATX inhibitors, as suggested here for EL244, would alleviate systemic effects, including potential impacts on T-cell homeostasis.

Inhaled administration of EL244, restricting its effects in the lung, is predicted to avoid drug-drug interactions with the SOC compounds, as previously shown with GLPG1690 and nintedanib ^64^. Inhaled therapies have long been a cornerstone in the management of obstructive lung diseases due to their ability to deliver medication directly to the active disease site, thereby increasing drug tissue concentration and efficacy while reducing the dosage ^65^. Moreover, inhalation bypasses the intracellular and extracellular drug-metabolising enzymes in the liver and the gastrointestinal tract, while minimising potential systemic side effects ^65^. Despite these clear advantages, their application in idiopathic pulmonary fibrosis (IPF) has, paradoxically, been relatively underexplored. However, the inhaled delivery of the SOC compounds is currently explored in ongoing clinical trials (NCT06329401, NCT06951217)^66,67^, while several other agents targeting various factors (TGF, WNT, NAC, GAL-3, CNN2, MMP7) are also delivered by inhalation in ongoing clinical trials ^68^. In some cases, delivery by inhalation has been reported to have superior efficacy when compared to oral administration ^69,70^. Notably, inhaled Treprostinil administration in patients with ILD and associated pulmonary hypertension was associated with improvements in FVC, particularly evident in patients with IPF ^71,72^, further suggesting drug delivery by inhalation as the preferred method for treating IPF.

EL244, apart from exhibiting potent ATX inhibitory activity and PPARγ activation, also showed a favourable ADMET profile (intrinsic clearance, hepatotoxicity, and cardiotoxicity), outperforming reference compounds (Table 1). Inhaled administration of EL244 attenuated BLM-induced pulmonary fibrosis, both in preventive and therapeutic treatment modes. Importantly, EL244 was able to restore respiratory functions, a functional pathologic readout better aligned with the human disease that was previously suggested to be necessary for improving drug testing in animal models ^50,73^. Using translatable pulmonary function tests, such as measurement of murine FVC, a latest development in instrumentation, and activity monitoring, could identify better candidates to be tested in IPF clinical trials, where the main readouts are FVC and 6-min walking distance (6MWD) ^73^.

Inhaled EL244 decreased LPA levels in the BALF but not in the plasma, indicating that the EL244-mediated effects were confined to the lung. Moreover, EL244 decreased all LPA species and total LPA levels, much like the genetic deletion of ATX ^74^, while increased levels of most LPA species (16:0, 16:1, 18:1, 18:2, 20:4) have been reported in IPF patients ^75^. Therefore, EL244 exhibits efficient and physiologically relevant target engagement. This broader approach, compared to focusing on a single LPA species (18:2), offers a more effective PD biomarker for evaluating the target engagement of ATX inhibitors. ATX is necessary for embryonic development ^74^, but is largely dispensable for adult healthy homeostasis, including the lung ^44^. Therefore, LPA long-term reductions in the lung with EL244 are not anticipated to be toxic, pending the results of inhaled toxicity studies.

Beyond ATX inhibition, EL244 is also a potent PPARγ agonist, as it was developed from the repositioned TGZ. PPARγ plays a central role in regulating metabolic reprogramming by integrating signals from key metabolic sensing systems ^23^. In the context of pulmonary fibrosis, TGFβ, the main pro-fibrotic factor, has been shown to suppress PPARγ, while, conversely, PPARγ activation has been demonstrated to suppress TGFβ-induced mitochondrial activation ^76^. Interestingly, it has been more recently suggested that the pathogenesis of pulmonary fibrosis involves TGFβ-induced differentiation of lipofibroblasts, a novel subset of pulmonary fibroblasts, into myofibroblasts, and that activation of PPARγ inhibits this differentiation, suppressing the development of pulmonary fibrosis ^77^. Moreover, PPARγ activation in macrophages, cells central to pulmonary immunometabolism, has been suggested to directly regulate several macrophage functions, including differentiation from monocytes and polarisation ^78^. In accordance with the multifactorial role of PPARγ in pulmonary fibrosis, PPARγ agonists have been reported to attenuate BLM-induced pulmonary fibrosis ^26^, suggesting that a part of the therapeutic efficacy of EL244 can be attributed to PPARγ agonism.

Intriguingly, LPA, the enzymatic product of ATX, has been suggested to suppress PPARγ ^22^, although the underlying mechanisms, possibly including the Wnt pathway ^79^, remain understudied. ATX inhibition by PF-8380 has been reported to moderately upregulate PPARγ and its target genes in adipocytes *in vitro* ^80,81^. PPARγ and target gene expression were also increased in the adipose tissue upon the genetic deletion of ATX ^82^, suggesting a close interplay of the ATX/LPA axis and PPARγ.

EL244 reduces LPA production and its associated profibrotic effects, while also preventing LPA-mediated suppression of PPARγ. Simultaneously, it directly activates PPARγ, promoting metabolic reprogramming, and inhibits ATX expression, creating a positive feedback loop with therapeutic benefits. This dual mechanism of action enables EL244 to target pulmonary fibrosis through two interrelated cellular pathways, providing a distinct advantage over single-agent therapies and suggesting that EL244 is a promising therapeutic candidate for the treatment of IPF and ILDs.

## MATERIALS AND METHODS

### Virtual Screening (VS)

VS of the Prestwick Chemical Library was performed using the crystal structure of the ATX protein (PDB ID: 2XRG)^57^ via the Enalos Asclepios KNIME pipeline (Fig. S1)^83^. Computational details for VS, molecular docking and molecular dynamics (MD) simulations are provided in the Supplementary Materials and Methods.

**ATX inhibitory activity *in vitro*** of examined compounds was assessed with the Amplex Red assay, as previously reported ^28,31,32^, and as recently published in detail ^33^.

**ATX inhibitory activity *ex vivo*** of examined compounds was assessed via the colorimetric TOOS assay, as previously reported ^28,31,32^.

**PPARα/γ/δ hybrid reporter gene assays and Isothermal Titration Calorimetry** (ITC) were performed as previously described ^45^

**Mice** were housed and bred under specific pathogen-free (SPF) conditions at 20–22°C with 55 ± 5% humidity, a 12-hour light/dark cycle, and unrestricted access to food and water. Experimental cohorts were randomly assigned and consisted of sex- and age-matched littermates. Daily health checks ensured proper animal welfare, and no unanticipated mortality occurred. At designated time points, euthanasia was carried out in a CO₂ chamber using a gradual fill method to ensure humane treatment. All procedures received approval from the Protocol Evaluation Committee (PEC) of the Biomedical Sciences Research Center “Alexander Fleming” and were licensed by the Veterinary Authority of the Attica region, Greece (#927781, 2023). The institution’s Animal Welfare Body (AWB) provided oversight of animal welfare compliance.

**BLM-induced pulmonary fibrosis** was induced via the oropharyngeal aspiration of BLM (0.8 U/Kg), as previously described ^50,51,84^, and as detailed at protocols.io.

**Drug inhalation** of compound 5 and EL244 to conscious, softly restrained mice was performed using the inExpose system (Scireq), as previously described ^51^. Compound 5 and EL244 were diluted in Kolliphor (15% and 10%, respectively) to final concentrations of 6.5 and 9.02 mg/ml, respectively. Volumes of 1 mL (for 5) and 0.6 mL (for EL244) were aerosolised over 10 and 15 min, respectively in groups of 6 mice, corresponding to a final estimated dose of 15mg/Kg per mouse. Control groups received an aerosolised solution of Kolliphor (10-15%) in saline.

**Precision cult lung slices (PCLS)** were utilised as previously reported ^51^, and as described in detail in the recently published dedicated protocol ^47^.

**MS/MS** was performed as previously reported ^85^ and as described in detail in the supplementary Materials and methods.

#### RNA sequencing

Briefly, the QuantSeq 3’mRNA-Seq Library Prep Kit (Lexogen) was used according to the manufacturer’s instructions. The libraries were then used for DNB preparation and sequencing on a DNBSEQ-G400 platform, utilising a G400 App-A FCS SE100 High-throughput Sequencing Set (MGI Tech Co., Ltd.), also according to the manufacturer’s instructions. Data analysis was performed as previously reported ^86^, and as described in detail in the Supplementary Materials and Methods.

**HDX-MS** was performed using a dual-head parallel Trajan LEAP automation system (Carrboro, NC, USA), as detailed in the supplementary Materials and methods. Following the HDX-MS community guidelines, Tables S4 and S4B include the HDX summary and data table, respectively.

### Statistical Analysis

Statistical significance was assessed with the GraphPad software and its built-in recommendations. One-way ANOVA and post hoc Tukey’s test were used in the case of normally distributed data. In the case of unequal SDs, Brown-Forsythe and Welch ANOVA and post-hoc Games-Howell’s multiple comparison test were used. In the case of non-normally distributed data, Kruskal-Wallis test was performed, followed by post hoc Dunn’s test. Data were presented as box and whiskers graphs depicting all experimental values as well as the medians. ^*/**/***/****^ denote p<0.05/0.01/0.001/0.0001, respectively. Statistical analysis of RNA sequencing results is described in detail in the Supplementary Materials and Methods.

## Supplementary Materials

Figs. S1 to S12

Tables S1 to S9

Supplementary Materials and Methods

## ACKNOWLEDGEMENTS

### Authors Contribution

KDP and AF performed all HVTS computations and MD analyses. EL and AM synthesised all compounds. EL, CM, SGD, EK, IT and AM performed PK/PD analyses. E-AS, EK and CM tested ATX inhibition of compounds, while SW and DM tested PPARγ agonism. PK, DN, SM and CM performed animal studies and all related readout assays. JPRP and AP performed and analysed HDX-MS. KMA and AUW supervised the clinical applicability and translatability of findings in the context of the current state of the art. AG and PH performed and analysed RNA sequencing. The manuscript was written by AM with the assistance of KDP, JPRP, PK, and CM. It was edited by VA and critically reviewed by all authors.

### Funding

This research was co-financed by the European Union and Greek national funds through the Operational Program Competitiveness, Entrepreneurship and Innovation and the European Regional Development Fund, via the General Secretariat for Research and Innovation (GSRI) grant (T1EDK-0049 to VA), and the Hellenic Foundation for Research and Innovation (HFRI) grants (#3565 to VA, #01144 to CM, #7337 to ANM, #3780 to PH, #5691 to AG). It was also partly supported by the European Commission (EC)(#101037509 to AF) and by computing time awarded on the Cyclone supercomputer of The Cyprus Institute (#pro24a01). The funders had no role in the study’s design, data collection, analysis, or interpretation, nor in the writing of the manuscript or the decision to publish the results.

### Data availability

RNA sequencing data have been deposited to Gene Expression Omnibus (GEO) (#GSE297484). Following the HDX-MS community guidelines, Tables S4 and S4B include the HDX summary and data table, respectively.

