## Supplementary Materials and Methods for "An aerosolised dual-action Autotaxin inhibitor-PPARγ agonist for the treatment of pulmonary fibrosis"

*[1]Institute for Bioinnovation, Biomedical Sciences Research Center "Alexander Fleming", 16672 Athens, Greece. [2]Institute for Fundamental Biomedical Research, Biomedical Sciences Research Center "Alexander Fleming", 16672 Athens, Greece. [3]Department of Biology, National & Kapodistrian University of Athens, 15772 Athens, Greece. [4]Department of ChemoInformatics, Novamechanics Ltd., 1070 Nicosia, Cyprus. [5]Department of Pharmacy, Ludwig-Maximilians-Universität München, 81377 Munich, Germany. [6]Faculty of Biology, Medicine and Health, University of Manchester, Manchester M13 9PT, U.K. [7]Manchester Institute of Biotechnology, University of Manchester, Manchester M1 7DN, U.K. [8]Department of Biochemistry and Biotechnology, University of Thessaly, 41334 Larisa, Greece. [9]Pereleman School of Medicine, University of Pennsylvania, 19104 Philadelphia, USA. [10] Uni-Pharma S.A., 14564 Athens, Greece. [11]Department of Respiratory Medicine, School of Medicine, University of Crete; 70013 Heraklion, Greece. [12]Interstitial Lung Disease Unit, Royal Brompton Hospital, London, SW3 6NP, UK. [13]National Heart and Lung Institute, Imperial College London, London SW3 6LY, UK. [14]Margaret Turner Warwick Centre for Fibrosing Lung Disease, Imperial College London, London, UK.*

### Supplementary Materials and Methods

### Computational

#### Virtual screening

Virtual screening (VS) of the Prestwick Chemical Library was performed by means of the rDock molecular docking software (*1*). Initially, the crystal structure of the ATX protein (PDB ID: 2XRG)(*2*) was retrieved from the Protein Data Bank and pre-processed using PDBFixer (*3*) and pdb4amber from AmberTools21 (*4*) via the "Protein Structure Preparation" branch of the Enalos Asclepios KNIME pipeline (Fig. S1)(*5*). Preparation steps included addition of missing heavy and hydrogen atoms, reconstruction of non-terminal loops, conversion of non-standard residues to their standard counterparts, resolution of alternate atomic positions, and removal of heteroatoms. Simultaneously, the Prestwick compounds were prepared using their SMILES strings via the "Ligand Structure Preparation" workflow branch in the Asclepios KNIME pipeline. Hydrogens were added using OpenBabel (*6*) through the AsclepiosAddHydrogens node, adjusting the protonation state to pH 7.4. The AsclepiosGenerate3DCoords node then converted the 2D structures to 3D conformers through energy minimization. VS calculations were carried out using the RunRxDock node with the RxDock engine (*1*). The binding site was defined based on the co-crystallized inhibitor using the reference ligand method, with a cavity radius of 6.0 Å and a small sphere radius of 1.5 Å. Cavity mapping employed a 0.5 Å resolution grid and the RbtCavityGridSF scoring function. Parameters were set to accept a maximum of one cavity with a minimum volume of 100 $Å^3$. A cavity restrain function (weight = 1) was applied to prevent ligand escape during docking. Each compound was subjected to 50 independent docking runs. The resulting poses were ranked based on their intermolecular score, which reflects protein-ligand binding energy. Although total docking scores typically include intramolecular energy contributions, this term was excluded due to its known reduction in predictive reliability (*7, 8*). Conformations were further evaluated based on root-mean-square deviation (RMSD) relative to the reference pose, prioritizing top-scoring ligands for *in vitro* testing on the basis of both docking score and key interactions with the $Zn^{2+}$ ions, catalytic residue Thr209, and the hydrophobic pocket.

#### Molecular docking

Molecular docking at ATX and PPARγ was performed using NovaMechanics Asclepios (*5, 9*). 3D structures and physiologically relevant protonation states (pH 7.4) were generated via Gypsum-DL (*10*), which converts SMILES or SDF files into 3D models with alternate ionization, tautomeric, chiral, and conformational states. Dimorphite-DL (*11*) was used for empirical protonation state prediction, based on substructure searches and a curated ionizable compound database. To refine results, pKa values of key functional groups were predicted with MolGpka (*12*), which uses a graph-convolutional neural network trained on chemical patterns. Only the most probable protonation variant per compound, aligned with physiological pH, was retained. Initial geometry optimization used UFF via Gypsum-DL (*13*), followed by refinement with xTB (v6.6.0) (*14*) using the GFN2-xTB method (*15*) and ALPB solvation (*16*). Docking cavities were defined via the reference ligand method; 100 RxDock runs per ligand were performed. The PPARγ ligand-binding domain (PDB ID: 5YCP) complexed with RGZ was used as the structural model (*17*). Only crystallographically known stereoisomers of RGZ, TGZ, and PGZ were considered, in line with prior studies (*18*). Docking was executed using RxDock, a fork of rDock (*19*), and poses were ranked by intermolecular scores reflecting protein-ligand binding free energy (*20*).

**Molecular Dynamics Simulations**

The top-scoring docking poses for each ligand were used as initial conformations in Molecular Dynamics (MD) simulations with OpenMM 7.5 (*3*), implemented via the Asclepios KNIME workflow. Protonation states of ATX and PPARγ residues were assigned using the AsclepiosPDBFixer node, which integrates PDBFixer and pdb4amber (*21*) to add missing residues/heavy atoms, remove heteroatoms, and standardize residue names. Disulfide bonds were incorporated using pdb4amber and prepareforleap (*21*), as applicable. The AMBER14SB force field (*22*) was used for both proteins. Given its functional relevance, only the N-glycan linked to ATX Asn524 (*23*) was retained, modelled using the glycan from PDB 5MHP (*24*) and parameterized with GLYCAM06 24 (*25*). Ligand geometry optimization was performed at the B3LYP/6-31G* level (*26-28*), followed by HF/6-31G* electrostatic potential (ESP) calculations using Gaussian 09 (version D.01), via PyRED on the R.E.D. Server (*29-32*). RESP charges were derived using the standard two-stage fitting (*31*), and GAFF2.1 (*33-35*) parameters were assigned. TIP3P water molecules (*36*) were added with a 10 Å buffer, using rectangular periodic boundary conditions. Neutralizing $Na^+$/$Cl^-$ ions were introduced. Energy minimization (20,000 steps) applied positional restraints on protein-ligand atoms, gradually reduced every 5,000 steps (100 → 0 kcal $mol^{-1}Å^{-2}$). Systems were then equilibrated (1 fs time step): heating (200 ps, NVT) from 0→300 K in 3 ps intervals using a Langevin thermostat (*37*), with 20 kcal $mol^{-1}Å^{-2}$ restraints on non-hydrogen atoms. For ATX, additional $Zn^{2+}$-centered restraints (100 kcal $mol^{-1}Å^{-2}$) were applied on coordinating atoms (*23*) during both equilibration and production. Pressure was gradually ramped from 0.1 to 1 atm (20 ps intervals, NPT) using a Monte Carlo barostat (*38, 39*), with 2 kcal $mol^{-1}Å^{-2}$ restraints, later lifted over 1 ns, followed by 1 ns unrestrained NPT. Long-range electrostatics were handled via Particle Mesh Ewald (*40*), with 10 Å cutoffs for electrostatics and van der Waals interactions. Production MD (200 ns) used 2-fs time steps, constraining bonds involving H atoms. A total of 10,000 frames were saved and analyzed using CPPTRAJ from AmberTools21 (*4*). RMSDs (mass-weighted) were computed for Cα atoms (proteins) and heavy atoms (ligands). Hydrogen bonds were evaluated over the final 5,000 frames, with a 3.5 Å donor–acceptor distance and 150° angle cutoff. Clustering was performed using DBSCAN (epsilon = 3 Å, minPts = 25, sieve = 10), and centroid structures were visualized using PoseEdit (*41*)and ChimeraX (*42-44*). MM-GBSA binding free energies were calculated using the AmberTools21 MMPBSA.py implementation in the Enalos Asclepios workflow (*4*), based on gas-phase interaction and solvation energies (GB model) (*45*). Enthalpic contributions were averaged over the final 5,000 frames, with per-residue decomposition. Standard error of the mean (SEM) was used to report statistical uncertainties: $\sigma/\sqrt{N}$ (N = 5000).

**HDX-MS**

All HDX experiments were performed using a dual head parallel Trajan LEAP automation system (Carrboro, NC, USA) similarly as described previously (*46*). Briefly, 5µL of reconstituted Autotaxin (1.25mg/mL) with and without EL244, was diluted into 100µl of labelling buffer (10mM PBS, 150mM sodium chloride, pD =7.4 in D2O). pD was determined using pH measurements with a glass electrode and corrected for the isotopic effect (*47*). Samples were labelled in quadruplicate for each labelling time (30s, 300s and 3000s) at 25°C. For experiments with bounded Autotaxin, both the protein stock and labelling buffer contained 50µM of the ligand to ensure >95% bound protein during labelling (*48*). Afterwards, 100 µL of labelled samples were quenched by mixing with equal amounts of precooled quench buffer (100mM glycine, 3M urea, 0.5M TCEP, pH 2.3 in water) at 1°C and held at that temperature for 120s to improve disulfide bond reduction. Non-deuterated controls and initial peptide map samples were prepared identically but using 10mM PBS, 150mM sodium chloride, pH =7.4 in water as labelling buffer. Immediately after, 198µL of the sample was injected into a temperature-controlled chromatography cabinet connected to two ACQUITY I Class binary pumps. Sample was passed at 200 µL min–1 with 0.1% formic acid in water through a pepsin column (2.1 × 30 mm) immobilized in house kept at room temperature for 210s. Resulting peptic peptides were trapped and desalted in a XBridge C8 (2.1 × 5 mm, 5 µm) VanGuard precolumn (Waters Corporation) and separated using a Waters XBridge peptide BEH C18 analytical column (1.0 × 50 mm, 3.5 µm) with an 10 min linear gradient of 0.1% formic acid in acetonitrile increasing from 13 to 40% at 100 µL min–1. To avoid peptide carryover, the pepsin column was washed two times after each run with 85 µL of pepsin wash (2 M guanidine HCl, 5% acetonitrile, 100 mM phosphate buffer, pH 2.5) and blanks ran every 3 runs. Peptide masses were measured using a Waters SELECT SERIES cyclic IMS QTOF system using positive mode ionization and single pass ion mobility. All HDX experiments were measured using HDMS mode while initial peptide maps used HDMSE with a collision energy ramp of 20-45v (*49*).

HDMSE files were processed using ProteinLynxGlobal Server (PLGS, v.3.0.3) (Waters Corporation) and then imported into DynamX v. 3.0.0 (Waters Corporation) as described previously (*49*). Peptides were manually curated in DynamX and HDX data exported into Deuteros v. 2.0 (*50*) for statistical analysis and identification of significant peptides (*51*). Significant peptides were mapped into the crystal structure 2XR9 (*52*). To allow access to the HDX data of this study, the HDX data summary table (Table S4) and the HDX data table (Table S4B) are included in the supporting information as per consensus guidelines (*53*).

### RNA sequencing

RNA quality was assessed on an Agilent Tapestation 4150 with the High Sensitivity RNA ScreenTape (Agilent), both according to the manufacturer's protocols. For library preparation, the QuantSeq 3‘mRNA-Seq Library Prep Kit (Lexogen) was used according to manufacturer’s instructions. Briefly, 500 ng of RNA from each sample was used for first-strand synthesis, followed by RNA template removal, and second-strand synthesis initiated by random primers. In-line barcodes were introduced at this step, and this was followed by magnetic bead-based purification. The resulting libraries were amplified for 15 cycles and re-purified, with quantity and quality assessed on a Qubit 4 Fluorometer with the Invitrogen Qubit dsDNA HS Assay and an Agilent Tapestation 4150 with the High Sensitivity D1000 assay kit, respectively. The quantified libraries were pooled equimolarly; 50ng of total library mix was used for Adapter Conversion PCR Amplification using the Universal Library Conversion Kit (App-A), Version: V1.0 (MGI Tech Co., Ltd.), according to the manufacturer's instructions.

The quantity and quality of the purified adapter conversion PCR (AC-PCR) product were evaluated on a Qubit 4 Fluorometer with the Invitrogen Qubit dsDNA HS Assay and on an Agilent Tapestation 4150 with the High Sensitivity D1000 assay kit, respectively. This was followed by DNA denaturation, Single-Strand Circularisation, Enzymatic Digestion, Enzymatic Digestion Product Cleanup and Quality Control steps on 1 pmol of AC-PCR product, according to the Universal Library Conversion Kit (App-A) User Manual (Version: A3).

Finally, 60 fmol of ssCirDNA were used for DNB preparation and sequencing on a DNBSEQ-G400 platform at the BSRC Alexander Fleming Genomics Facility, using a G400 App-A FCS SE100 High-throughput Sequencing Set (MGI Tech Co., Ltd.), according to the manufacturer's instructions.

Alignment of the FASTQ files was performed against the mouse genome build mm10 using a custom Bash script, as previously described (*54*). Read counting on the 3’ UTRs and differential expression analysis were conducted using the metaseqR2 bioconductor package (*55*). Genes corresponding to the following biotypes were excluded: polymorphic_pseudogene, processed_transcript, pseudogene, IG_V_pseudogene, misc_RNA, IG_C_gene, IG_J_gene, IG_D_gene, IG_LV_gene, TR_V_gene, TR_V_pseudogene, TEC, and processed_pseudogene. Gene counts were normalized with DESEQ (*56*) through metaseqR2. Genes with more than 10 counts in at least 50% of the samples were kept for further processing. For differential expression analysis, the PANDORA algorithm (*57*) was used to estimate a meta p-value integrating the results from the DESEQ, DESEQ2 (*58*), edgeR (*59*), limma (*60*), NBPSeq (*61*), and NOISeq (*62*) algorithms. All additional parameters were set to default values.

Pathway enrichment analysis was performed for the Gene Ontology Biological process (BP) and Molecular Function (MF) terms through GeneCodis 4 (*63*). The volcano plots, scatter plots, Venn diagrams, and dot plots were generated using the ggplot2 Bioconductor package.

All code of the RNA sequencing analysis is available at the github repository: https://github.com/alex-galaras/matralis_et.al.2025.git.

**HPLC-MS/MS**

Plasma (50µl), BALF (300 µl) or homogenized lung tissue (10-50 mg) were mixed with ice-cold PBS spiked with the internal standard mixture (LPA-d9, Cayman) in a glass tube (total final volume 1ml). 2 mL of ice-cold chloroform and 1 ml of methanol was added in each sample, following by vortexing for 1 min and then centrifugation at 4 $^{o}$C for 5 min at 4000 g. The lower transparent organic phase (chloroform phase-contains the neutral lipids) was collected in a glass tube. The remaining aqueous phase (upper phase) was left on ice for 10 minutes and acidified (pH 3-4) with Formic Acid 10% in water. Then 1.5 ml chloroform was added followed by thorough mixing for 1 min and centrifugation at 4 $^{o}$C for 5 min at 2000 g. The lower organic phase was collected and neutralized to pH 6-7. The neutralized organic phase from the acid extraction were evaporated to dryness. After evaporation we redissolve in water/methanol (1:9) with 0.1% formic acid and run in the mass spec. All MS quantitation lipid standards were purchased from Cayman unless otherwise mentioned. All LPA species analyzed in this study were quantified using the multiple reaction monitoring (MRM) scanning method on an Waters-QQQ mass spectrometer. All data were acquired and analyzed using the Waters data analysis software. The LC separation was achieved using a Gemini 5U C18 column (Phenomenex, 5 µm, 50 × 4.6 mm) coupled to a Gemini guard column (Phenomenex, 4 × 3 mm). The LC solvents were as follows: buffer A, 85% CAN, 5% MeOH + 0.1% formic acid + 10% 10 mM ammonium formate buffer pH5; buffer B, 95:5 (v/v) ACN/MeOH + 0.1% formic acid. A typical LC-run was 35 min, with the following solvent run sequence post injection: 0.35 ml/min 5% B for 5 min, linear gradient of B from 0–95% over 25 min,and re-equilibration with 0.5 ml/min of 5% B for 5 min. All lipid estimations were performed using an electrospray ion (ESI) source, with the following MS parameters: turbo spray ion source, medium collision gas, curtain gas = 20 l/min, ion spray voltage =−5000 V (negative mode), at 350 °C. All the endogenous lipid species were quantified by measuring the area under the curve in comparison to the respective internal standard, and then normalized to the total protein content of the liver tissue. In the following table is the MRM used for LPA quantification.

| Species targeted | Precursor Ion Mass | Product Ion Mass | Collision Energy (V) | Ionization Mode |
|---|---|---|---|---|
| LPA 14 | 381 | 153 | 20 | Negative |
| LPA 16 | 409 | 153 | 20 | Negative |
| LPA 18 | 437 | 153 | 20 | Negative |
| LPA 18:1 | 435 | 153 | 20 | Negative |
| LPA 18:2 | 433 | 153 | 20 | Negative |
| LPA 20:4 | 457 | 153 | 20 | Negative |
| LPA 22:6 | 481 | 153 | 20 | Negative |

### Synthesis

#### Synthesis of compounds 1 and 4.

***tert*-butyl (2-(4-formylphenoxy)ethyl)carbamate (i) (10.1021/jm0510880).** To a solution of 2-(boc-amino)ethanol (1 g, 6.20 mmol), 4-hydroxybenzaldehyde (0.91 g, 7.44 mmol) and triphenyl phosphine (2.44 g, 9.30 mmol) in dry THF (25 mL) was added *di*-isopropyl azodicarboxylate (1.83 mL, 9.30 mmol) dropwise at 0 $^{o}$C. The mixture was then allowed to stir at rt for 2.5 h. The solvent was removed under vacuum, the residue was dissolved in ethyl acetate (50 mL), washed with 1N NaOH (10 mL), water (10 mL) and brine (10 mL), dried ($Na_2SO_4$), filtered and concentrated in vacuum. The residue was purified by flash column chromatography eluted with hexane:ethyl acetate (4:1), furnishing the desired product as a white solid. Yield = 1.64 g (quant.). $^1$H-NMR ($CDCl_3$, 400 MHz) $\delta$ 1.48 (s, 9H), 3.55 (d, $J$ = 4.5 Hz, 2H), 4.11 (t, $J$ = 5.0 Hz, 2H), 5.13 (brs, 1H), 7.05 (d, $J$ = 8.5 Hz, 2H), 7.86 (d, $J$ = 9.2 Hz, 2H), 9.91 (s, 1H). MS [ESI+] m/z 266.1 $[M + H]^+$.

***tert*-butyl (*E*)-(2-(4-((2,4-dioxothiazolidin-5-ylidene)methyl)phenoxy)ethyl)carbamate (ii) (10.1021/jm0510880).** To a solution of compound **i** (1.50 g, 5.65 mmol) in dry toluene (15 mL), 2,4-thiazolidinone (0.80 g, 6.79 mmol) was added, followed by piperidine (0.28 mL, 2.83 mmol) and acetic acid (0.162 mL, 0.83 mmol). The mixture was stirred at reflux for 8 h and then it was allowed at rt overnight. The resulting solid precipitated was filtered, washed with toluene (3 mL) and hexane (5 mL), and dried at 45 $^{o}$C overnight to give the desired product as a beige/brownish amorphous solid. Yield =1.63 g (80%). $^1$H-NMR (DMSO-$d_6$, 400 MHz) $\delta$ 1.38 (s, 9H), 3.31 (m, 2H), 4.06 (m, 2H), 7.01 (s, 1H), 7.10 (d, $J$ = 7.4 Hz, 2H), 7.55 (d, $J$ = 8.7 Hz, 2H), 7.75 (s, 1H), 12.49 (brs, 1H). MS [ESI+] m/z 365.2 $[M + H]^+$.

***tert*-butyl (2-(4-((2,4-dioxothiazolidin-5-yl)methyl)phenoxy)ethyl)carbamate (iii) (10.1021/jm0510880).** The thiazolidinone derivative **ii** (0.80 g, 2.20 mmol) and dry magnesium turnings (1.07 g, 43.91 mmol) were added in a flask and air was removed in vacuum. Then Argon was placed in the flask and anhydrous methanol (27 mL) was added. The mixture was stirred at rt and under Ar for 4h. The reaction mixture was acidified with 6N HCl to pH 5-6 and extracted with dichloromethane (2 x 25 mL) and the combined organic phase was washed with water (15 mL) and brine (15 mL), dried ($Na_2SO_4$), filtered and concentrated in vacuum. The residue was purified by flash column chromatography eluted with hexane:ethyl acetate (3:2), affording the desired compound as a yellow oil. Yield = 0.37 g (46%). $^1$H-NMR ($CDCl_3$, 400 MHz) $\delta$ 1.50 (s, 9H), 3.13 (dd, $J_1$ = 3.5 Hz, $J_2$ = 14 Hz, 1H), 3.44 (dd, $J_1$ = 3.5 Hz, $J_2$ = 14.0 Hz, 1H), 3.58 (brd, $J$ = 4 Hz, 2H), 4.06 (t, $J$ = 5 Hz, 2H), 4.53 (dd, $J_1$ = 3.5 Hz, $J_2$ = 9.5 Hz, 1H), 5.03 (brs, 1H), 6.89 (d, $J$ = 8.0 Hz, 2H), 7.21 (d, $J$ = 8.0 Hz, 2H), 8.99 (brs, 1H). MS [ESI+] m/z 367.1 $[M + H]^+$.

**Synthesis of compounds iv and v (10.1021/jm0510880).** A solution of 4N HCl in dioxane (1.90 mL, 7.61 mmol) was added at once to compounds **ii** and **iii** (0.22 g, 0.61 mmol) and the mixture was stirred at rt for 4h. The solvent was distilled in vacuum, the residue was washed with diethyl ether (15 mL) and dried, providing the desired products as a white solids.

**(*E*)-5-(4-(2-aminoethoxy)benzylidene)thiazolidine-2,4-dione hydrochloride (iv).** Yield = 0.111 g (quant.). $^1$H-NMR ($CH_3OD$, 400 MHz) $\delta$. 3.45 (dd, $J_1$ = 4.2 Hz, $J_2$ = 14.1 Hz, 1H), 4.55 (brs, 2H), 6.98 (d, J = 8.0 Hz, 2H), 7.23 (d, J = 8.0 Hz, 2H), 7.91 (s, 1H). MS [ESI+] m/z 302.0 $[M + H]^+$.

**5-(4-(2-aminoethoxy)benzyl)thiazolidine-2,4-dione hydrochloride (v).** Yield = 0.115 g (quant.). $^1$H-NMR ($CH_3OD$, 400 MHz) $\delta$ 3.18 (dd, $J_1$ = 9.0 Hz, $J_2$ = 14.0 Hz, 1H), 3.39 (brs, 2H), 3.44 (dd, $J_1$ = 4.0 Hz, $J_2$ = 14.0 Hz, 1H), 4.22 (brs, 2H), 4.75 (dd, $J_1$ = 4.0 Hz, $J_2$ = 9.5 Hz, 1H), 6.95 (d, *J* = 8.0 Hz, 2H), 7.26 (d, *J* = 8.0 Hz, 2H). MS [ESI+] m/z 304.1 $[M + H]^+$.

**2-amino-4-(4-fluorophenyl)thiazole-5-carbonitrile (vi) (10.1021/acs.jmedchem.7b00032).** To a solution of 4-fluorobenzoylacetonitrile (0.47 g, 2.89 mmol) in dry ethanol (6 mL), dry pyridine (0.24 mL, 2.89 mmol) was added and the mixture was stirred at 70 °C for 20 min and then cooled to rt. A previously stirred suspension of thiourea (0.44 g, 5.79 mmol) and iodine (0.73 g, 2.89 mmol) in dry ethanol (4 mL) was slowly added and the mixture was stirred at rt for 2h. Cold water (40 mL) was added and the resulting precipitate was filtered, washed with water (10 mL) and hexane (15 mL) and dried in vacuum to afford the desired product as a yellow solid. Yield = 0.63 g (quant). $^1$H-NMR (dmso-$d_6$, 400 MHz) $\delta$ 7.37 (t, *J* = 8.9 Hz, 2H), 7.95-8.00 (m, 2H), 8.25 (s, 2H). MS [ESI+] m/z 220.0 $[M + H]^+$.

**2-chloro-4-(4-fluorophenyl)thiazole-5-carbonitrile (vii) (10.1021/acs.jmedchem.7b00032).** To a solution of anhydrous $CuCl_2$ (0.47 g, 3.47 mmol) in dry $CH_3CN$ (6.5 mL) was added dropwise *tert*-butoxy nitrite (0.45 g, 4.34 mmol) and the mixture was stirred at rt for 45 min. Then, compound **vi** (0.63 g, 2.89 mmol) was added in portions and stirring was continued for further 2h. The reaction mixture was carefully quenched with 1N HCl (10 mL) and stirred for 15 min. The organic phase was separated, the aqueous phase was extracted with ethyl acetate (20 mL) and the combined organic phase was washed with brine (10 mL), dried ($Na_2SO_4$), filtered and concentrated in vacuum. The crude product was filtered on a silica plug eluted with dichloromethane. Solvents were distilled in vacuum and the residue was triturated with hexane, filtered and dried. The product is isolated as a bright orange thick solid. Yield = 0.49 g (71%). $^1$H-NMR ($CDCl_3$, 400 MHz) $\delta$ 7.19-7.25 (m, 2H), 8.12-8.17 (m, 2H). MS [ESI+] m/z 240.0 $[M + H]^+$.

**Synthesis of compounds 1 and 4.** Compounds **iv** or **v** (0.25 g, 0.83 mmol) and **vii** (0.18 g, 0.76 mmol) were dissolved in dry DMSO (7 mL), DIPEA (0.33 mL, 1.89 mmol) was added and the mixture was stirred at 100 °C for 8 h and at rt overnight. Water (15 mL) was added and the mixture was extracted with dichloromethane (2 x 30 mL). The combined organic phase was washed with water (2 x 20 mL) and brine (20 mL), dried ($Na_2SO_4$), filtered and concentrated in vacuum. The residue was purified by flash column chromatography eluted with hexane:ethyl acetate (7:3 to 1:1) furnishing the final products.

**(*E*)-2-((2-(4-((2,4-dioxothiazolidin-5-ylidene)methyl)phenoxy)ethyl)amino)-4-(4-fluorophenyl)thiazole-5-carbonitrile (1).** Light brown solid. Yield = 0.34 g (95%). $^{1}$H-NMR (dmso-d$_6$, 400 MHz) $\delta$ 3.80-3.81 (m, 2H), 4.27-4.28 (m, 2H), 7.15 (d, J = 8.2 Hz, 2H), 7.36 (t, J = 8.6 Hz, 2H), 7.56 (d, J = 8.3 Hz, 2H), 7.76 (s, 1H), 7.98-8.02 (m, 2H), 8.99-9.02 (m, 1H), 12.50 (s, 1H). $^{13}$C-NMR ($CDCl_3$, 100 MHz) $\delta$ 44.1, 66.6, 84.1, 115.6, 115.9, 116.2, 116.4, 121.0, 126.3, 129.5, 129.6, 130.3, 130.4, 132.2, 132.6, 160.0, 160.4, 162.1, 164.6, 167.9, 168.4, 170.2. HRMS (ESI): *m/z* 467.0652 $[M+H]^+$, [Calc. 467.0648].

**2-((2-(4-((2,4-dioxothiazolidin-5-yl)methyl)phenoxy)ethyl)amino)-4-(4-fluorophenyl)thiazole-5-carbonitrile (4).** Off-yellow amorphous solid. Yield = 0.20 g (56%). $^{1}$H-NMR ($CDCl_3$, 400 MHz) $\delta$ 3.14-3.20 (m, 1H), 3.42-3.47 (m, 1H), 3.81-3.85 (m, 2H), 4.21-4.23 (m, 2H), 4.50-4.54 (m, 1H), 6.43 (brs, 1H), 6.89 (d, J = 8.3 Hz, 2H), 7.15-7.20 (m, 4H), 8.05-8.08 (m, 3H). $^{13}$C-NMR ($CDCl_3$, 100 MHz) $\delta$ 37.6, 45.3, 53.4, 65.8, 77.2, 114.5, 114.8 (2C), 115.9 (2C), 116.1, 128.5, 130.0, 130.1, 130.7 (2C), 157.6, 159.5, 162.1, 169.8, 170.0, 173.6. HRMS (ESI): *m/z* 469.0811 $[M+H]^+$, [Calc. 469.0804].

**Synthesis of compounds 2 and 5.**

**4-(2-bromoethoxy)benzaldehyde (viii) (10.1002/anie.202105103).** 4-hydroxybenzaldehyde (0.75 g, 6.14 mmol) was dissolved in dry $CH_3CN$ (45 mL) and 1,2-dibromoethane (5.29 mL, 61.4 mmol) and $K_2CO_3$ (1.55 g, 11.2 mmol) were subsequently added. The mixture was stirred under reflux for 20 h, cooled to rt, water (45 mL) was added and the mixture was extracted with $Et_2O$ (2 x 30 mL). The combined organic phase was washed with brine (25 mL), dried ($Na_2SO_4$), filtered and concentrated in vacuum. The residue was recrystallized from $Et_2O$:hexane to give the product as a white solid. Yield = 0.92 g (65%). $^{1}$H-NMR ($CDCl_3$, 400 MHz) $\delta$ 3.69 (td, $J_1$ = 1.7 Hz, $J_2$ = 6.2 Hz, 2H), 4.40 (td, $J_1$ = 1.7 Hz, $J_2$ = 6.2 Hz, 2H), 7.04 (dd, $J_1$ = 1.7 Hz, $J_2$ = 8.7 Hz, 2H), 7.87 (dd, $J_1$ = 1.9 Hz, $J_2$ = 8.7 Hz, 2H), 9.92 (s, 1H). MS [ESI+] m/z 229.9 $[M + H]^+$.

**1-(*tert*-butyl) 4-(3,5-dichlorobenzyl) piperazine-1,4-dicarboxylate (x) (10.1021/acsmedchemlett.7b00312).** To a solution of (3,5-dichlorophenyl)methanol (1.50 g, 8.47 mmol) in dry DMF (15 mL) was added CDI (1.92 g, 11.86 mmol) and the reaction was stirred at 45 $^{o}$C for 2h. Then, 1-boc-piperazine (1.97 g, 10.59 mmol) was added and the reaction mixture was stirred at rt overnight. Water (30 mL) was added to the mixture and the precipitate was filtered, washed with water (2 x 10 mL) and hexane (10 mL) and dried. The crude product (white solid) was used immediately in the next step without further purification. Yield = 3.30 g (74%). MS [ESI+] m/z 390.1 $[M + H]^+$.

**3,5-dichlorobenzyl piperazine-1-carboxylate hydrochloride (ix) (10.1021/acsmedchemlett.7b00312).** 4N HCl in dioxane (16 mL, 63 mmol) was added to 1-(*tert*-butyl) 4-(3,5-dichlorobenzyl) piperazine-1,4-dicarboxylate (**xiii**, 2.44 g, 6.27

mmol) at 0 °C and the reaction mixture was stirred at rt for 3h. The solvent was evaporated under reduced pressure and the white solid remaining was used in the next step without further purification. Yield = 2 g (quant.). $^1$H-NMR ($CDCl_3$, 400 MHz) $\delta$ 3.09 (m, 4H), 3.65 (m, 4H), 5.10 (s, 2H), 7.47 (s, 2H), 7.57 (s, 1H), 9.49 (brs, 2H). MS [ESI+] m/z 326.1 $[M + H]^+$.

**3,5-dichlorobenzyl 4-(2-(4-formylphenoxy)ethyl)piperazine-1-carboxylate (xi).** A mixture of compounds **viii** (0.92 g, 4.02 mmol), **ix** (0.44 g, 4.42 mmol) and $NaHCO_3$ (1.35 g, 16.08 mmol) in dry DMF (20 mL) was stirred at 80 °C for 24 h and at 55 °C for 12 h. Then water (50 mL) was added, and the mixture was extracted with ethyl acetate (3 x 25 mL). The combined organic phase was washed with water (25 mL) and brine (25 mL), dried ($Na_2SO_4$), filtered and concentrated in vacuum. The residue was purified by flash column chromatography eluted with hexane:EtOAc (7:3 to 100% ethyl acetate), affording a yellowish oil which solidifies upon standing in the fridge. Yield = 1.76 g (71%). $^1$H-NMR (dmso-$d_6$, 400 MHz) $\delta$ 2.50-2.52 (m, 3H), 2.78 (t, J = 5.6 Hz, 2H), 2.97 (t, J = 5.6 Hz, 1H), 3.46 (m, 4H), 4.27 (t, J = 5.6 Hz, 2H), 5.11 (s, 2H), 7.12 (d, J = 8.4 Hz, 2H), 7.42 (s, 2H), 7.66 (d, J = 8.1 Hz, 2H), 7.70 (s, 1H), 9.90 (s, 1H). MS [ESI+] m/z 437.4 $[M + H]^+$.

**3,5-dichlorobenzyl (*E*)-4-(2-(4-((2,4-dioxothiazolidin-5-ylidene)methyl)phenoxy)ethyl)piperazine-1-carboxylate (2).** In an oven-dried round-bottom flask, compound **xi** (1.68 g, 3.85 mmol) and 2,4-thiazolidinedione (0.54 g, 4.62 mmol) were dispersed in dry toluene (16 mL). Then, piperidine (0.20 mL, 1.92 mmol) was added followed by acetic acid (0.11 mL, 1.92 mmol) and the mixture was refluxed overnight. The reaction mixture was left to cool at rt where a brownish solid precipitated. The mixture was filtered, washed with toluene (20 mL) and hexane (20 mL) and dried at 45 °C overnight. Off-yellow powder. Yield = 2 g (quant.). $^1$H-NMR (dmso-$d_6$, 400 MHz) $\delta$ 2.50-2.52 (m, 3H), 2.76 (t, $J$ = 5.6 Hz, 2H), 2.99 (t, $J$ = 5.6 Hz, 1H), 3.42 (brm, 4H), 4.17 (t, $J$ = 5.6 Hz, 2H), 5.08 (s, 2H), 7.10 (d, $J$ = 8.4 Hz, 2H), 7.42 (s, 2H), 7.53 (s, 1H), 7.56 (d, $J$ = 8.1 Hz, 2H), 7.70 (s, 1H). $^{13}$C-NMR (dmso-$d_6$, 100 MHz) $\delta$ 44.0 (2C), 53.2 (2C), 56.9, 65.2, 66.0, 115.3 (2C), 122.5, 126.6 (2C), 127.9, 128.9, 131.0 (2C), 134.2, 134.5 (2C), 141.8, 154.5, 158.6, 176.2, 183.5. HRMS (ESI): *m*/*z* 536.0815 $[M+H]^+$, [Calc. 536.0814].

**3,5-dichlorobenzyl4-(2-(4-((2,4-dioxothiazolidin-5-yl)methyl)phenoxy)ethyl)piperazine-1-carboxylate (5).** The thiazolidinone derivative **2** (0.45 g, 0.84 mmol) was mixed with water (25 mL), to which 5 drops of 0.5M aqueous solution of NaOH was added until pH 11. Then, a mixture of THF:DMF 2:1 (20 mL) was added followed by $CoCl_2 \cdot 6H_2O$ (0.128 g, 0.537 mmol), dimethylglyoxime (0.129 g, 1.107 mmol) and sodium borohydride (0.374 g, 9.88 mmol). The reaction mixture was stirred at room temperature for 24h, where a partial conversion of the starting material to the desired product was observed by TLC and MS. Then, more $CoCl_2 \cdot 6H_2O$ (0.128 g, 0.537 mmol), dimethylglyoxime (0.129 g, 1.107 mmol) and sodium borohydride (0.374 g, 9.88 mmol) were added and the mixture was stirred overnight. The pH of the reaction was adjusted to 3 with 6N HCl and then to 7-

8 by adding 1N NaOH. It was extracted with ethyl acetate (2 x 50 mL), the combined organic phase was washed with water (30 mL) and brine (30 mL), dried ($Na_2SO_4$), filtered and concentrated in vacuum. The residue was purified by flash column chromatography eluted with ethyl acetate, providing the desired product as a slightly yellow semisolid. Yield = 0.226 g (50%). $^1$H-NMR ($CDCl_3$, 400 MHz) $\delta$ 2.64 (brm, 4H), 2.87-289 (m, 2H), 3.09-3.14 (m, 1H), 3.40-3.45 (dd, $J_1$ = 3.8 Hz, $J_2$ = 14.2 Hz, 1H), 3.57-3.60 (m, 4H), 4.10-4.15 (m, 2H), 4.45 (dd, $J_1$ = 3.9 Hz, $J_2$ = 9.2 Hz, 1H), 5.09 (s, 2H), 6.85 (d, $J$ = 8.6 Hz, 2H), 7.16 (d, $J$ = 8.6 Hz, 2H), 7.24 (s, 2H), 7.28 (s, 1H), 7.33 (s, 1H). $^{13}$C-NMR ($CDCl_3$, 100 MHz) $\delta$ 37.7, 53.0, 53.7 (2C), 57.1, 60.4, 65.4, 65.6 (2C), 114.8 (2C), 126.1 (2C), 128.1, 128.2, 130.5 (2C), 135.1 (2C), 140.0, 154.7, 157.8, 170.5, 174.3. HRMS (ESI): *m/z* 538.0973 $[M+H]^+$, [Calc. 538.0970].

**Synthesis of compound 3.**

**2-(hydroxymethyl)-2,5,7,8-tetramethylchroman-6-ol (xii) (10.1248/cpb.48.272).** Lithium aluminum hydride ($LiAlH_4$, 0.194 g, 5.12 mmol) was suspended in dry THF (4 mL), and a solution of trolox (0.40 g, 1.60 mmol) in dry THF (6 mL) was added dropwise at room temperature under Argon. The mixture was stirred for 2.5 h and carefully quenched with cold water (10 mL). The aqueous layer was extracted with ethyl acetate (2 x 40 mL), the organic phase was washed with brine (25 mL), dried, filtered and concentrated in vacuum to give the desired product as a white solid. Yield = 0.24 g (63%).$^1$H-NMR ($CDCl_3$, 400 MHz) $\delta$ 1.22 (s, 3H), 1.73 (ddd, $J_1$ = 4.7 Hz, $J_2$ = 9.6 Hz, $J_3$ = 16.5 Hz, 1H), 1.99 (ddd, $J_1$ = 7.3 Hz, $J_2$ = 9.4 Hz, $J_3$ = 17.1 Hz, 1H), 2.12 (s, 3H), 2.14 (s, 3H), 2.21 (s, 3H), 2.62-2.71 (m, 2H), 3.62 (d, $J$ = 11.1 Hz, 1H), 3.67 (d, $J$ = 11.3 Hz, 1H).

**(6-((*tert*-butyldimethylsilyl)oxy)-2,5,7,8-tetramethylchroman-2-yl)methanol (xiii).** A suspension of sodium hydride (0.085 g of 60% oil dispersion, 2.133 mmol) in dry tetrahydrofuran (1.50 mL) was cooled at 0 $^o$C, and then a solution of the diol **xii** (0.24 g, 1.016 mmol) in dry THF (1 mL) was added dropwise under Argon. The mixture was stirred at the same temperature for 45 min, and then a solution of *tert*-butyldimethylsilyl chloride (0.199 g, 1.321 mmol) in dry THF (1 mL) was added dropwise at the same temperature. The reaction mixture was slowly warmed to room temperature and stirred for 3h. Upon completion, the mixture was quenched with cold water (5 mL) and extracted with hexane (2 x 20 mL). The combined organic phase was washed with brine (10 mL), dried, filtered and concentrated in vacuum. The residue was purified by flash column chromatography eluted hexane:EtOAc (97:3) to furnish the desired intermediate as a pale-yellow oil. Yield = 0.179 g (50%). $^1$H-NMR ($CDCl_3$, 400 MHz) $\delta$ 0.15 (s, 6H), 1.07 (s, 9H), 1.25 (s, 3H), 1.70-1.79 (m, 1H), 1.88 (brs, 1H), 1.97-2.11 (m, 1H), 2.11 (s, 3H), 2.13 (s, 3H), 2.15 (s, 3H), 2.62-2.68 (m, 2H), 3.63 (d, $J$ = 11.5 Hz, 2H).

**4-((6-hydroxy-2,5,7,8-tetramethylchroman-2-yl)methoxy)benzaldehyde (xiv).** To a suspension of potassium *tert*-butoxide (*t*-BuOK, 0.063 g, 0.562 mmol) in dry DMF (0.5 mL), a solution of the alcohol **xiii** (0.179 g, 0.511 mmol) in dry DMF (0.5 mL) was added dropwise at room temperature and under Argon. After stirring for 1 h, 4-

fluorobenzaldehyde was added and the mixture was heated at 80 °C for 6 h and at room temperature overnight. The reaction mixture was quenched with water (15 mL) and extracted with ethyl acetate (2 x 20 mL). The combined organic extracts were washed with water (15 mL) and brine (15 mL), dried, filtered and concentrated in vacuum. The residue was purified by flash column chromatography eluted with hexane:EtOAc (3:1) to furnish the desired (unprotected) intermediate as a pale-yellow oil, which was used directly in the next step. Yield = 0.055 g (32%). MS [ESI+] m/z 340.43 $[M + H]^+$.

**3-(4-Fluorobenzyl)thiazolidine-2,4-dione (xv) (10.1021/acs.jmedchem.8b00935).** A solution of thiazolidine-2,4-dione (1.17 g, 9.99 mmol) in DMF (5 mL), under Ar, was cooled to 0 °C, and sodium hydride (60% mineral oil dispersion, 0.36 g, 8.99 mmol) and a solution of 4-fluorobenzyl chloride (0.793 mL, 0.963 g, 6.66 mmol) in dry DMF (3 mL) were added. The reaction mixture was allowed to warm to room temperature over 5 h. The reaction mixture was then poured over crushed ice (50 mL), hexane (25 mL) was then added, and the product was allowed to crystallize overnight at 4 °C. The desired intermediate was collected as colourless needles via vacuum filtration and dried. Yield = 0.952 g (63%). $^1$H-NMR ($CDCl_3$, 400 MHz) $\delta$ 3.92 (s, 2H), 4.75 (s, 2H), 7.00-7.05 (m, 2H), 7.40-7.41 (m, 2H).

**(*E*)-3-(4-fluorobenzyl)-5-(4-((6-hydroxy-2,5,7,8-tetramethylchroman-2-yl)methoxy)benzylidene)thiazolidine-2,4-dione (3).** In an oven-dried microwave vial employed with a magnetic stirrer, the aldehyde **xiv** (0.051 g, 0.150 mmol) and the 2,4-thiazolidinedione derivative **xv** (0.054 g, 0.240 mmol) were placed followed by dry toluene (1 mL), piperidine (14.8 µL, 0.013 g, 0.023 mmol) and acetic acid (1.33 µL, 1.4 mg, 0.023 mmol). The mixture was stirred under reflux (111 °C) for 72h. The mixture was cooled to room temperature overnight, where a yellow solid precipitated. The solid was filtered, washed with toluene and hexane and dried at 40 °C overnight. Yellow semisolid. Yield = 11 mg (13.3%).$^1$H-NMR ($CDCl_3$, 400 MHz) $\delta$ 1.48 (s, 3H), 1.99 (s, 3H), 2.02 (s, 3H), 2.14 (s, 3H), 2.68-2.72 (m, 2H), 3.64-3.73 (m, 2H), 4.30-4.37 (m, 1H), 4.88 (s, 2H), 5.32 (s, 1H), 6.86 (d, J = 8.8 Hz, 2H), 7.01-7.05 (m, 3H), 7.42-7.47 (m, 4H), 7.87 (s, 1H). HRMS (ESI): *m/z* 547.1838 $[M+H]^+$, [Calc. 547.1829].

**Synthesis of EL244**

***tert*-butyl 4-(2-bromoethyl)piperidine-1-carboxylate (xvi) 10.1039/C6RA03841G).** In an oven-dried microwave vial employed with magnetic stirrer, *N*-Boc-4-piperidinethanol (0.521 g, 2.274 mmol) is dissolved in dichloromethane (10 mL). Then triphenylphosphine (0.835 g, 3.184 mmol) and carbon tetrabromide (1.207 g, 3.640 mmol) were added in portions, and the reaction mixture was stirred at room temperature for 72h. Upon completion, the solvent was evaporated and the residue was purified by silica gel flash column chromatography eluted with hexane:ethyl acetate (100% hexane to 5% ethyl acetate in hexane). Colourless oil. Yield: = 0.530 g (79%). $^1$H-NMR ($CDCl_3$, 400 MHz) $\delta$ 1.05-1.17 (m, 3H), 1.47 (s, 9H), 1.65-1.71 (m, 3H), 1.75-1.80 (m, 2H), 2.62-2.71 (m, 2H), 3.47 (t, *J* = 7.0 Hz, 1H), 4.00-4.14 (m, 2H). MS [ESI+] m/z 292.22 $[M + H]^+$.

***tert*-butyl 4-(2-(4-formylphenoxy)ethyl)piperidine-1-carboxylate (xvii).** In a round-bottom flask with magnetic stirrer, *tert*-butyl 4-(2-bromoethyl)piperidine-1-carboxylate (**xvi**) (0.530 g, 1.814 mmol), 4-hydroxybenzaldehyde (0.277 g, 2.268 mmol), caesium carbonate (1.478 g, 4.535 mmol) and dry DMF (3 mL) were consecutively added and the reaction mixture was stirred at 70 $^{o}$C for 6h and at room temperature overnight. Water (10 mL) was added and the mixture was extracted with ethyl acetate (3 x 20 mL). The combined extracts were washed with water (2 x 15 mL), saturated aqueous solution of sodium carbonate (2 x 10 mL) and brine (15 mL), dried ($Na_2SO_4$), filtered and concentrated in vacuum. The product was purified by silica gel flash column chromatography eluted with hexane:ethyl acetate (EtOAc 0-20%). Off-white/yellowish solid. Yield = 0.535 g (88%).$^1$H-NMR ($CDCl_3$, 400 MHz) $\delta$ 1.17-1.28 (m, 2H), 1.48 (s, 9H), 1.73-1.82 (m, 5H), 2,74 (t, *J* = 11.8 Hz, 2H), 4.12 (t, *J* = 6.2 Hz, 4H), 7.01 (d, *J* = 8.7 Hz, 2H), 7.86 (d, *J* = 8.8 Hz, 2H), 9.91 (s, 1H). MS [ESI+] m/z 333.43 $[M + H]^+$.

**4-(2-(piperidin-4-yl)ethoxy)benzaldehyde (xviii).** *Tert*-butyl 4-(2-(4-formylphenoxy)ethyl)piperidine-1-carboxylate (**xvii**) (0.531 g, 1.593 mmol) was dissolved in dry dichloromethane (4 mL) in a round-bottom flask and trifluoroacetic acid (3.50 mL, 5.447 g, 47.776 mmol) was subsequently added. The reaction mixture was stirred at room temperature for 2h, the solvent was evaporated in vacuum, saturated sodium bicarbonate aqueous solution (5 mL) was added and the product was extracted with ethyl acetate (3 x 15 mL). The combined organic phase was washed with brine (15 mL), dried ($Na_2SO_4$), filtered and concentrated in vacuum. Yellowish semisolid. Yield = 0.300 g (81%).$^1$H-NMR (dmso-$d_6$, 400 MHz) $\delta$ 1.35-1.43 (m, 2H), 1.74-1.85 (m, 5H), 2.74 (t, *J* = 13.7 Hz, 2H), 3.23 (d, *J* = 12.3 Hz, 2H), 4.11 (t, *J* = 6.0 Hz, 2H), 5.20 (brs, 1H), 7.00 (d, *J* = 8.5 Hz, 2H), 7.85 (d, *J* = 8.7 Hz, 2H), 9.90 (s, 1H). MS [ESI+] m/z 233.32 $[M + H]^+$.

**3,5-dichlorobenzyl 4-(2-(4-formylphenoxy)ethyl)piperidine-1-carboxylate (xix).** To a solution of 3,5-dichlorobenzyl alcohol (0.284 g, 1.607 mmol) in dry DMF (2.3 mL), carbonyl-*di*-imidazole (CDI, 0.365 g, 2.251 mmol) was added and the mixture was stirred at 45 $^{o}$C for 3h. Then, 4-(2-(piperidin-4-yl)ethoxy)benzaldehyde (**xviii**) (0.300 g, 1.286 mmol) was dissolved in dry DMF (2 mL) and added dropwise to the reaction mixture which was then stirred at 45 $^{o}$C for 3h and at room temperature overnight. Water (15 mL) was added and the mixture was extracted with diethyl ether (3 x 10 mL). The combined organic phase was washed with water (12 mL) and brine (12 mL), dried ($Na_2SO_4$), filtered and concentrated in vacuum. The product was purified by silica gel flash column chromatography eluted with hexane-ethyl acetate (85:15 to 70:30). Off-white semisolid. Yield = 0.533 g (95%). $^1$H-NMR ($CDCl_3$, 400 MHz) $\delta$ 1.24-1.31 (m, 2H), 1.79-1.81 (m, 5H), 2.78-2.93 (brm, 2H), 4.11-4.20 (m, 4H), 5.09 (s, 2H), 7.01 (d, *J* = 8.4 Hz, 2H), 7.25 (s, 2H), 7.32 (s, 1H), 7.86 (d, *J* = 8.5 Hz, 2H), 9.91 (s, 1H). MS [ESI+] m/z 436.33 $[M + H]^+$.

**3,5-dichlorobenzyl (*E*)-4-(2-(4-((2,4-dioxothiazolidin-5-ylidene)methyl)phenoxy)ethyl)piperidine-1-carboxylate (xx).** In an oven-dried microwave vial employed with a magnetic stirrer, 3,5-dichlorobenzyl 4-(2-(4-

formylphenoxy)ethyl)piperidine-1-carboxylate (**xix**) (0.521 g, 1.194 mmol) and 2,4-thiazolidinedione (0.168 g, 1.433 mmol) were placed followed by dry toluene (5 mL), piperidine (59.2 μL, 0.051 g, 0.597 mmol) and acetic acid (34.2 μL, 0.036 g, 0.597 mmol). The mixture was stirred under reflux (111 $^oC$) overnight. The mixture was cooled to room temperature, where a yellow solid precipitated. The solid was filtered, washed with toluene and hexane and dried at 45 $^oC$ overnight. Yellow powder. Yield = 0.500 g (78%).$^1$H-NMR (dmso-$d_6$, 400 MHz) $\delta$ 1.65-1.75 (m, 6H), 2.71-2.86 (brm, 2H), 3.00 (t, *J* = 5.7 Hz, 1H), 4.00 (d, *J* = 13.6 Hz, 2H), 4.10 (t, *J* = 5.8 Hz, 2H), 5.07 (s, 2H), 7.10 (d, *J* = 8.8 Hz, 2H), 7.41 (s, 2H), 7.54-7.56 (m, 3H), 7.75 (s, 1H), 12.51 (brs, 1H). MS [ESI+] m/z 536.42 $[M + H]^+$.

**3,5-dichlorobenzyl 4-(2-(4-((2,4-dioxothiazolidin-5-yl)methyl)phenoxy)ethyl)piperidine-1-carboxylate (EL244).**

**Catalyst:** 9 mg (0.038 mmol) of $CoCl_2{\cdot}6H_2O$ and 49 mg (0.413 mmol) of dimethylglyoxime were dissolved in 0.55 mL DMF under stirring, yielding a clear blue-green solution. It was maintained under stirring at rt until fully consumed.

**Reducing agent:** 0.177 g (4.670 mmol) of $NaBH_4$ were dissolved in 1.50 mL $H_2O$+ 0.5 mL of 0.1 M solution of NaOH under cooling in ice (at 0$^o$C). It was maintained in ice until fully consumed.

**Reaction:** 23.4 mg of NaOH (0.585 mmol)and subsequently 0.250 g (0.467 mmol) of 3,5-dichlorobenzyl 4-(2-(4-((2,4-dioxothiazolidin-5-yl)methyl)phenoxy)ethyl)piperidine-1-carboxylate (**xx**) were added in 5 mL of $H_2O$ and the obtained suspension was stirred and heated at 55 $^oC$ until a solution was formed. Part of the catalyst was added (0.15 mL of $CoCl_2$-DMG in DMF) during 1 min into the solution, followed by part of the reducing agent (0.50 mL of $NaBH_4$ in $H_2O$) during 2 min, and the mixture was stirred at 55 $^oC$ for 1 h. The same procedure was repeated 3 more times (each time, addition of 1/4 of the catalyst followed by 1/4 of the reducing agent, followed by 1 h stirring at 55 $^oC$, i.e. 1 addition every 1 hour). On completion of the additions, the mixture was stirred at 45 $^oC$ overnight. The following day an aqueous solution of 6 N HCl (10 mL) was added. The aqueous phase was extracted with EtOAc (2 x 20 mL). The combined organic phase was washed with water (20 mL) and brine (20 mL), dried, filtered and concentrated in vacuum. The product was purified via silica gel flash column chromatography eluted with hexane:EtOAc 3:2. White crystalline solid. Yield = 0.110 g (44%). $^1$H-NMR (dmso-$d_6$, 400 MHz) $\delta$ 1.05-1.10 (m, 2H), 1.64-1.74 (m, 5H), 2.73-2.85 (brm, 2H), 3.03-3.09 (m, 1H), 3.28 (d, *J* = 4.4 Hz, 1H), 3.97-4.01 (m, 4H), 4.87 (dd, $J_1$ = 9.0 Hz, $J_2$ = 4.3 Hz, 1H), 5.07 (s, 2H), 6.87 (d, *J* = 8.2 Hz, 2H), 7.14 (d, *J* = 8.3 Hz, 2H), 7.41 (s, 2H), 7.56 (s, 1H), 12.01 (brs, 1H). $^{13}$C-NMR (dmso-$d_6$, 100 MHz) $\delta$ 32.0, 32.6 (2C), 35.6, 36.7, 44.2 (2C), 53.5, 65.0, 65.5, 114.8 (2C), 126.6 (2C), 127.9, 128.9, 130.8 (2C), 134.5 (2C), 141.9, 154.5, 158.1, 172.2, 176.2. HRMS (ESI): *m/z* 537.1019 $[M+H]^+$, [Calc. 537.1018].

### NMR spectra of the final compounds

Compound **1** – $^{1}$H-NMR (dmso-d$_6$)

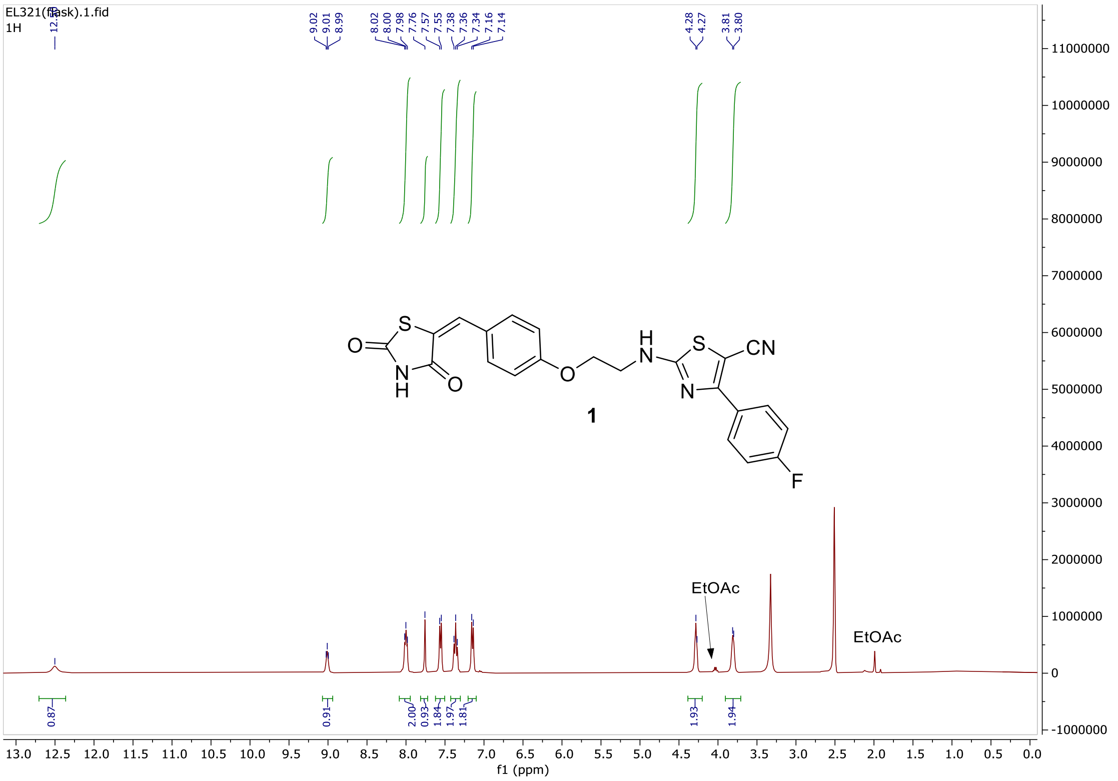

Compound **1** – $^{13}$C-NMR (dmso-d$_6$)

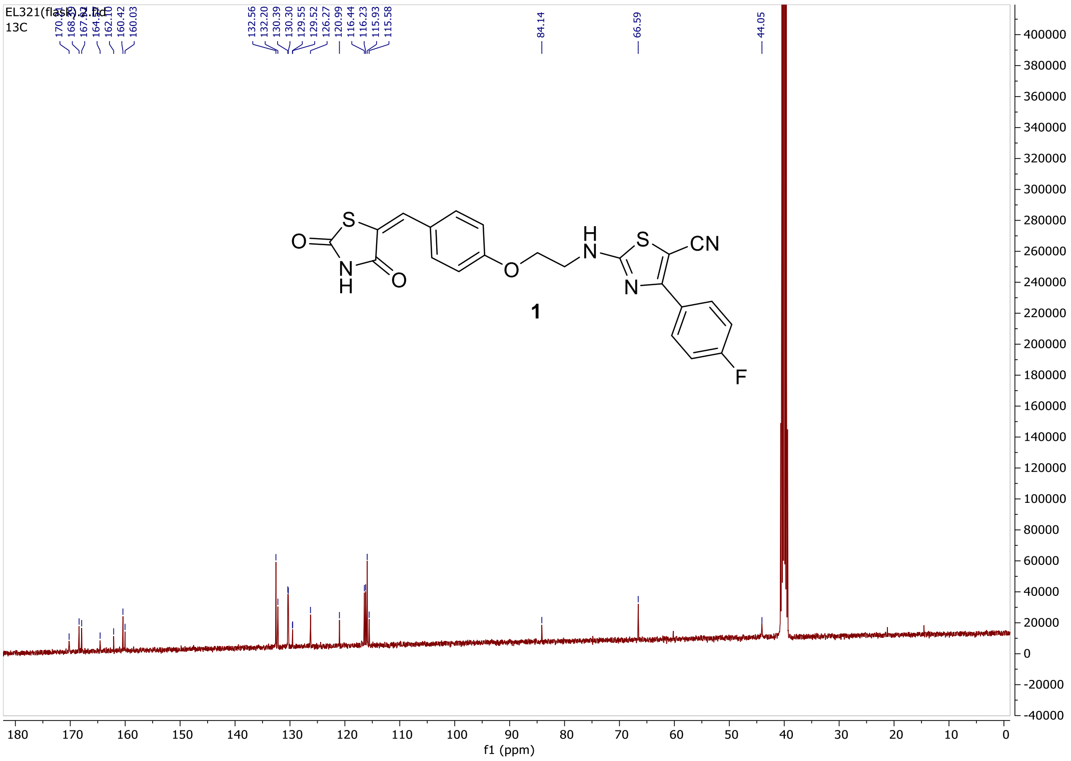

### Compound **2** – $^{1}$H-NMR (dmso-d$_6$)

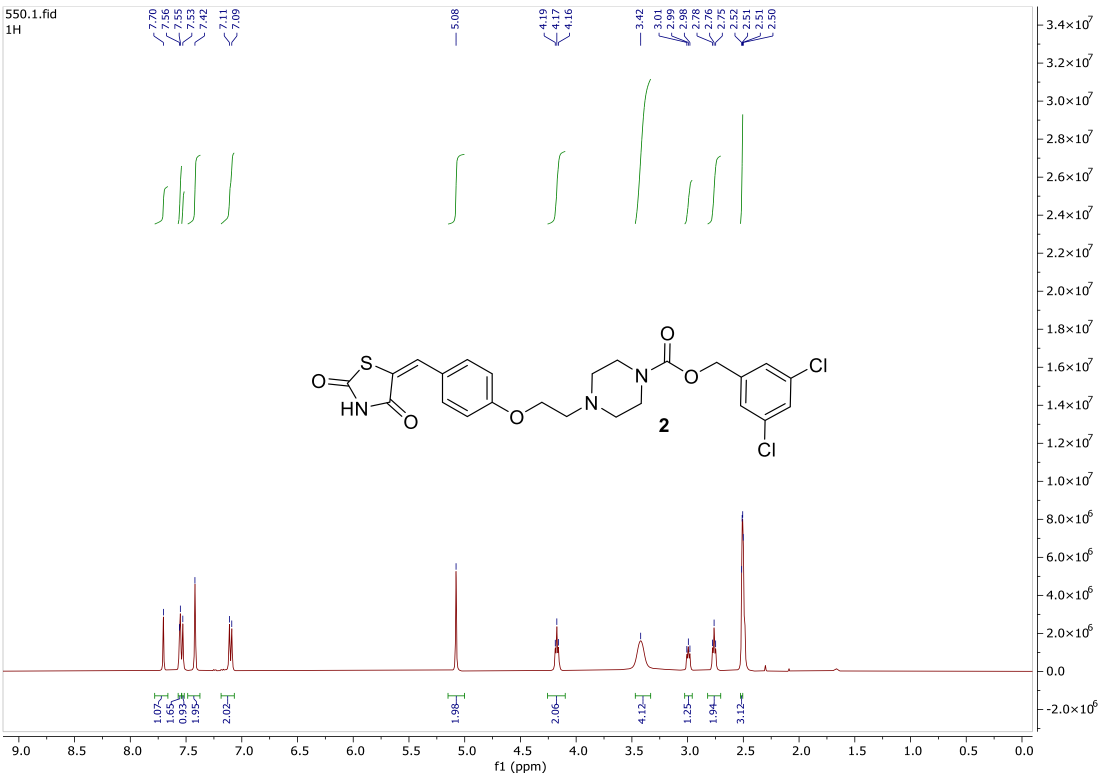


### Compound **2** – $^{13}$C-NMR (dmso-d$_6$)

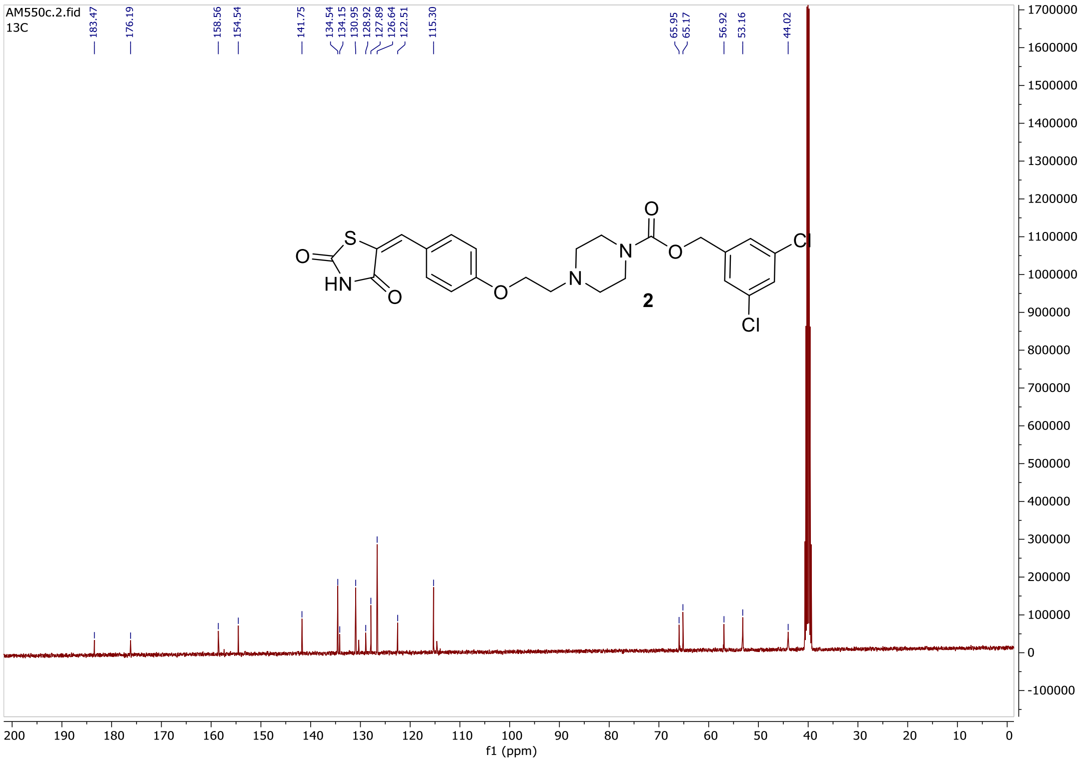

Compound **3** – [1]H-NMR ($CDCl_3$)

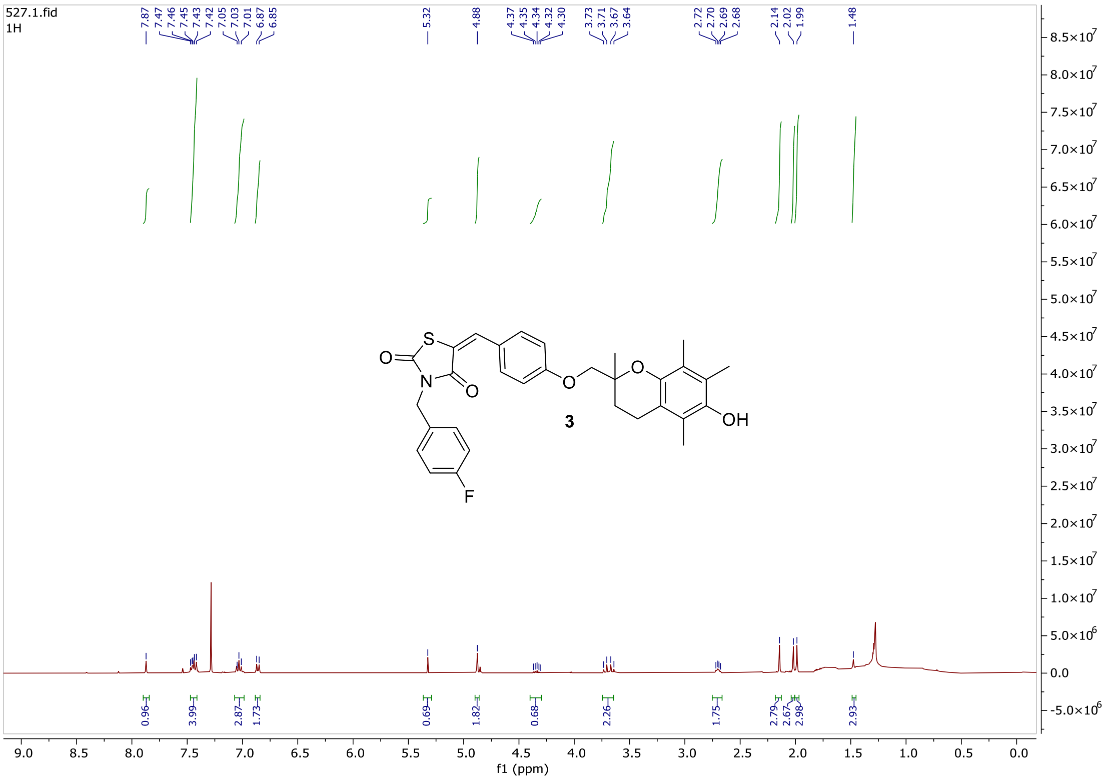

Compound **4** – [1]H-NMR ($CDCl_3$)

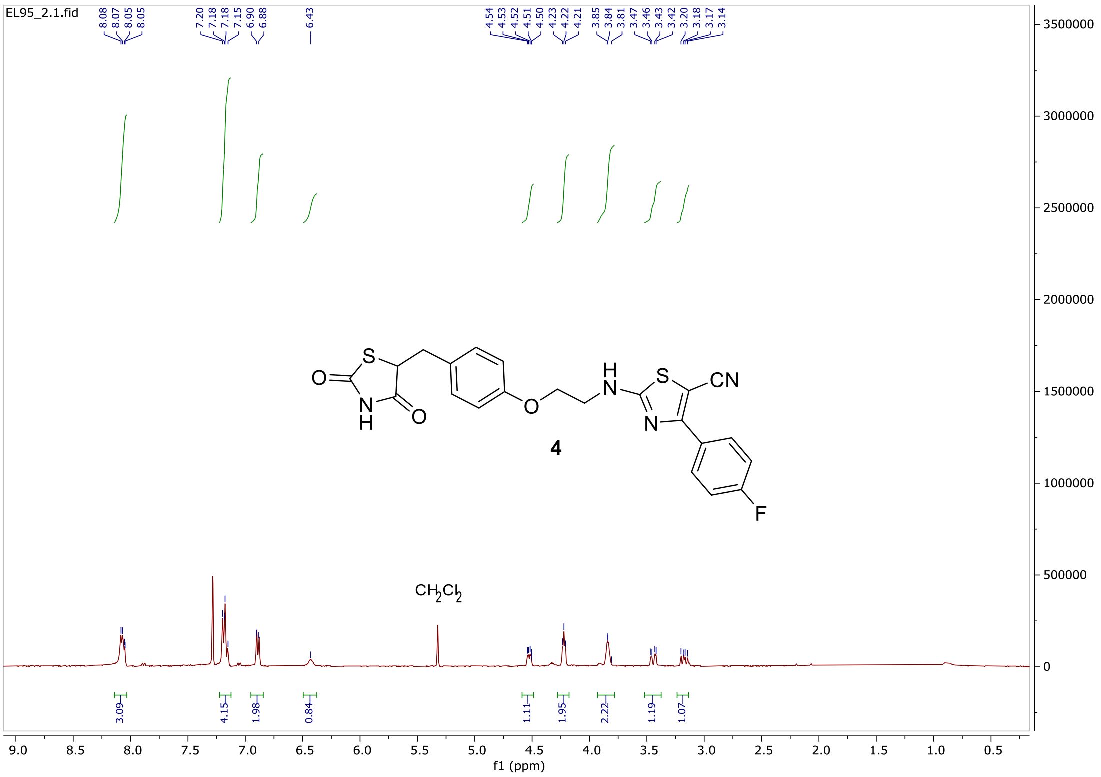

Compound **4** – $^{13}$C-NMR (CDCl$_3$)

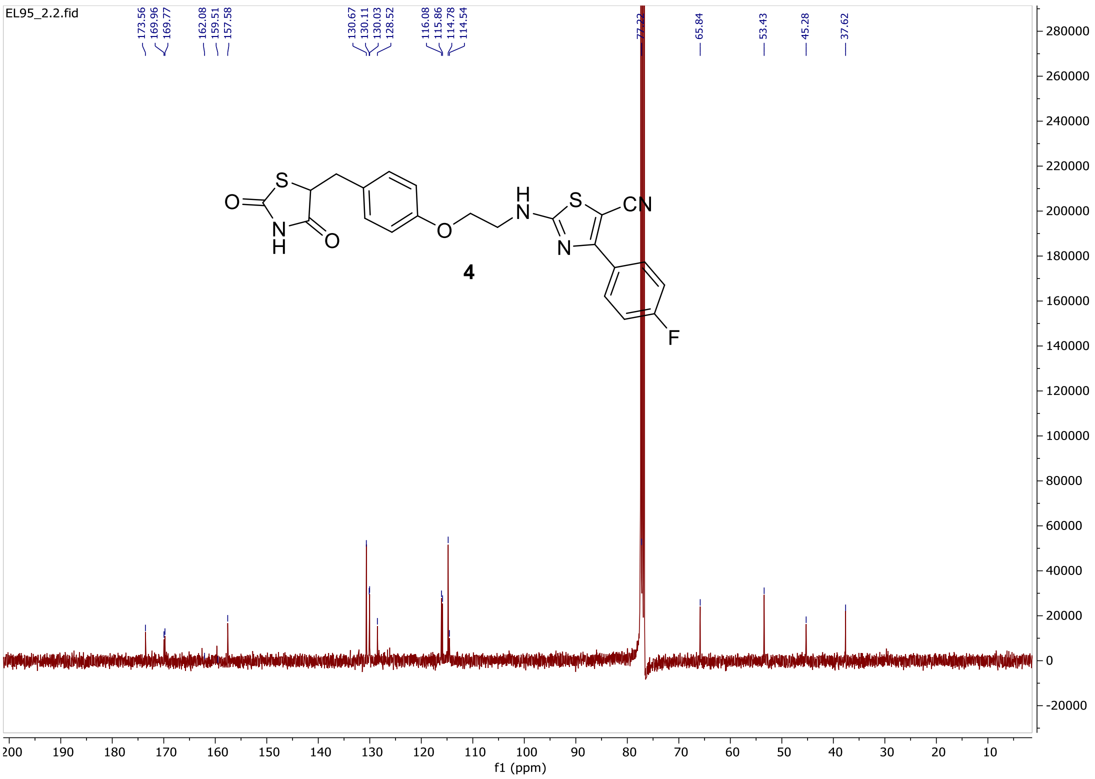


Compound **5** – $^{1}$H-NMR (CDCl$_3$)

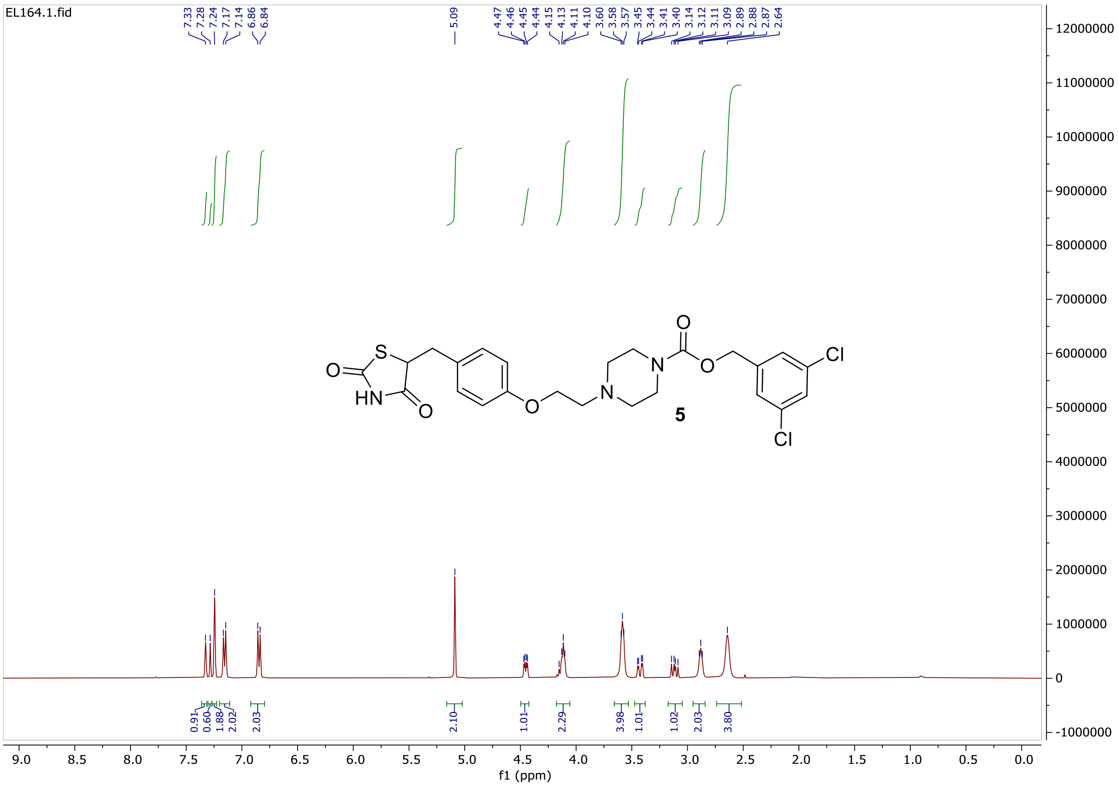

Compound **5** – $^{13}$C-NMR ($CDCl_3$)

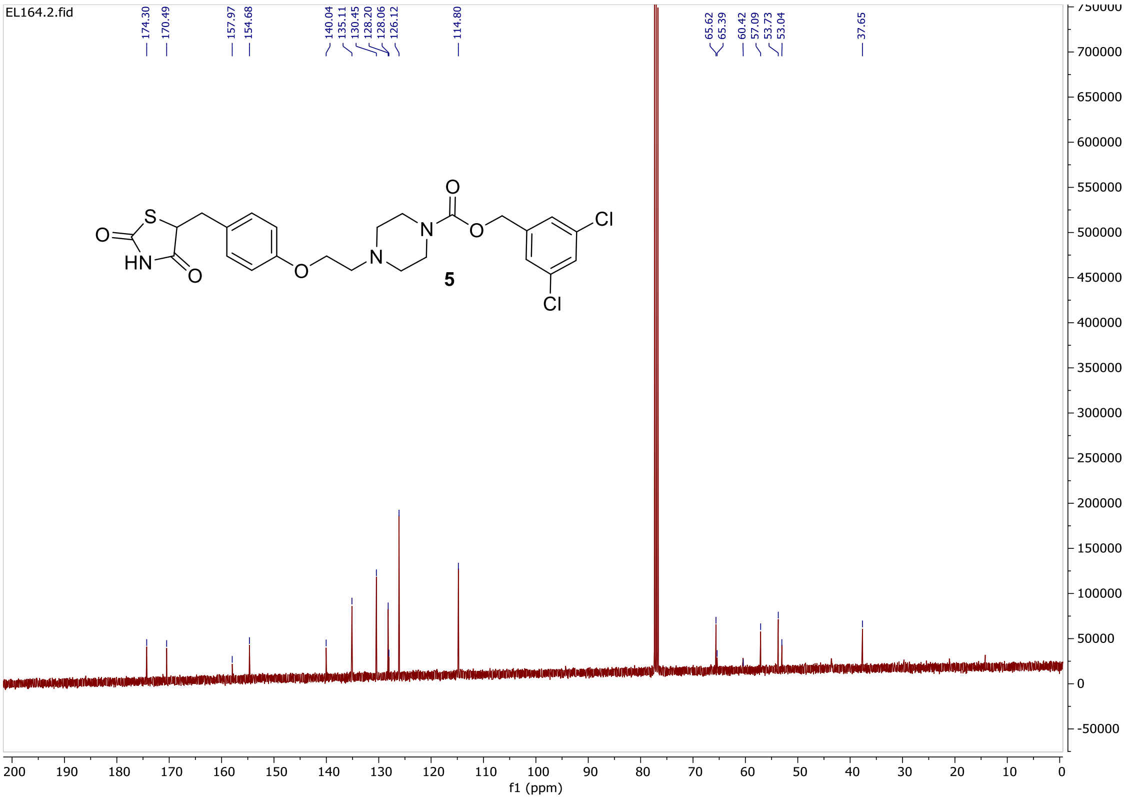


Compound **EL244** – $^{1}$H-NMR (dmso-$d_6$)

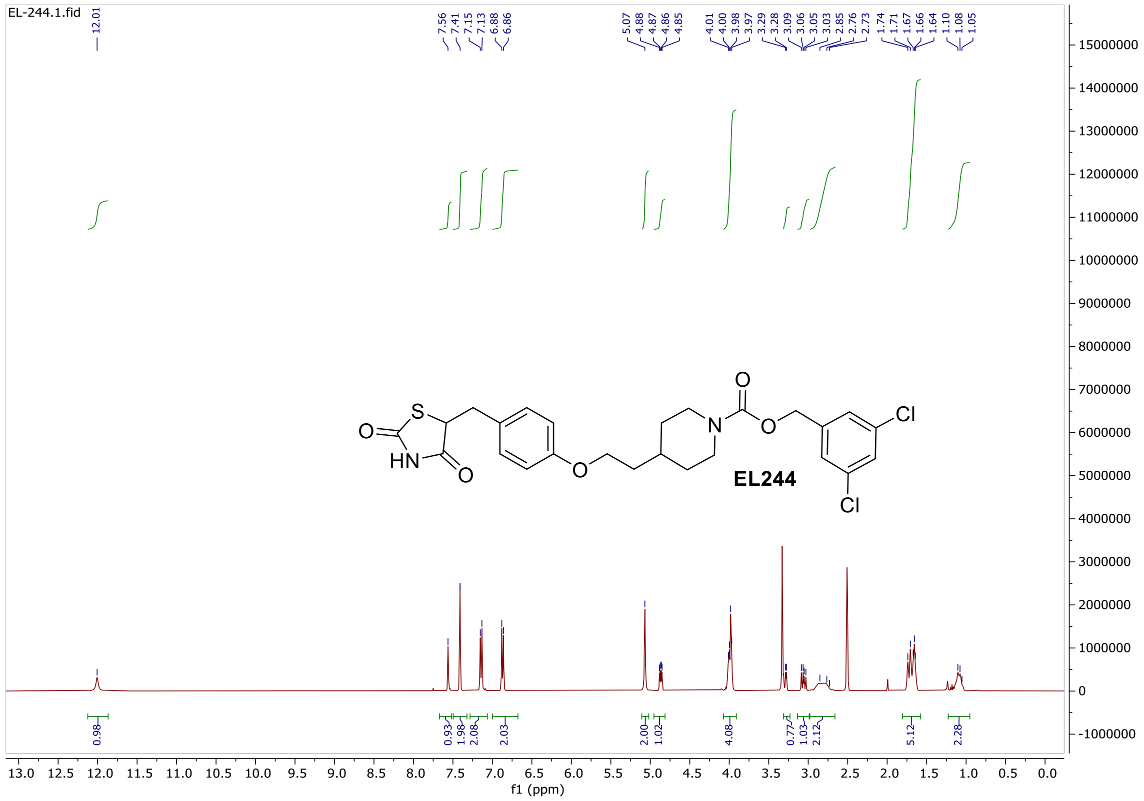

Compound **EL244** – $^{13}$C-NMR (DMSO-$d_6$)

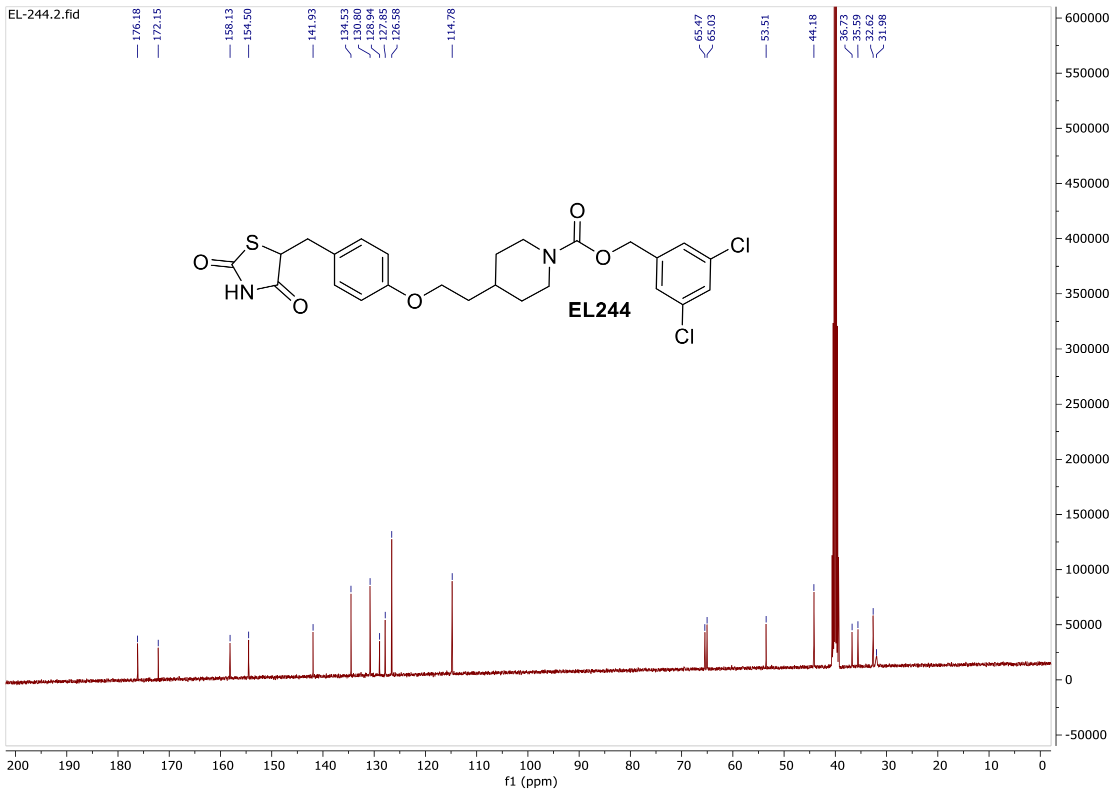
EL-244.2.fid
176.18
172.15
158.13
154.50
141.93
134.53
130.80
128.94
127.85
126.58
114.78
65.47
65.03
53.51
44.18
36.73
35.59
32.62
31.98
EL244
f1 (ppm)
