## Supplementary Figures for "An aerosolised dual-action Autotaxin inhibitor-PPARγ agonist for the treatment of pulmonary fibrosis"

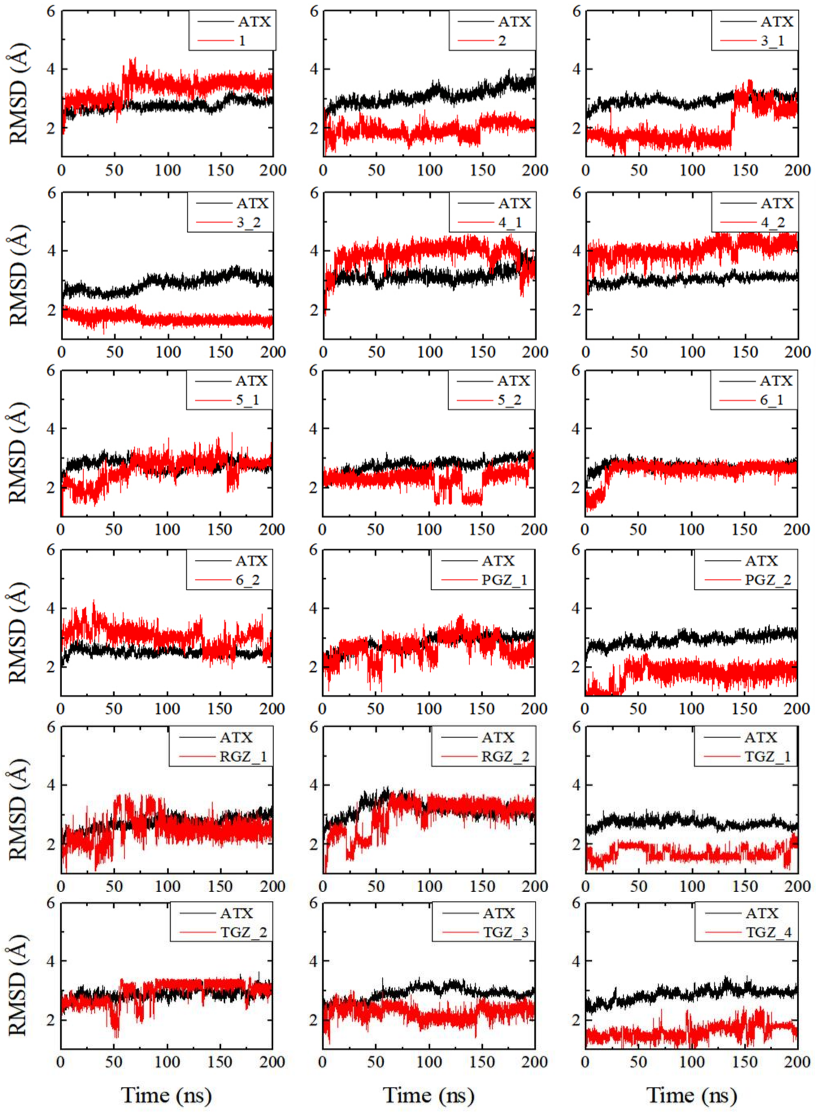


**Fig. S1. RMSD plots for the ATX protein systems analyzed in their complex bound states with TGZ diastereoisomers.**

A B

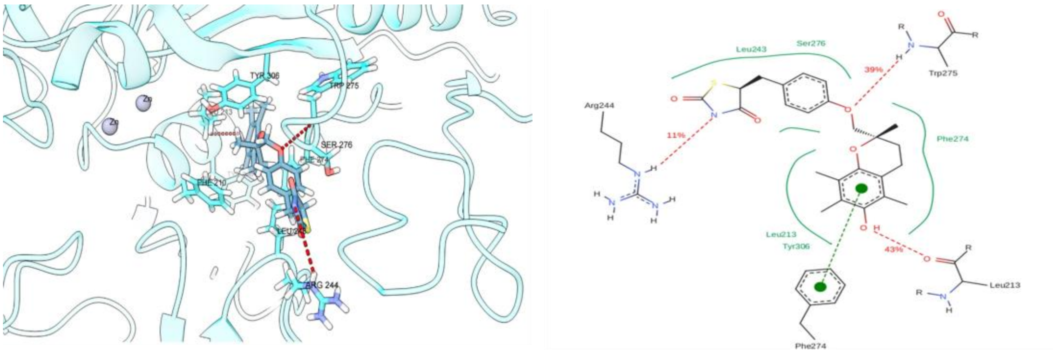


**Fig. S2. MD simulation of isomer TGZ_4 in complex with ATX.** (**A**) Three- and (**B**) two- dimensional representations.

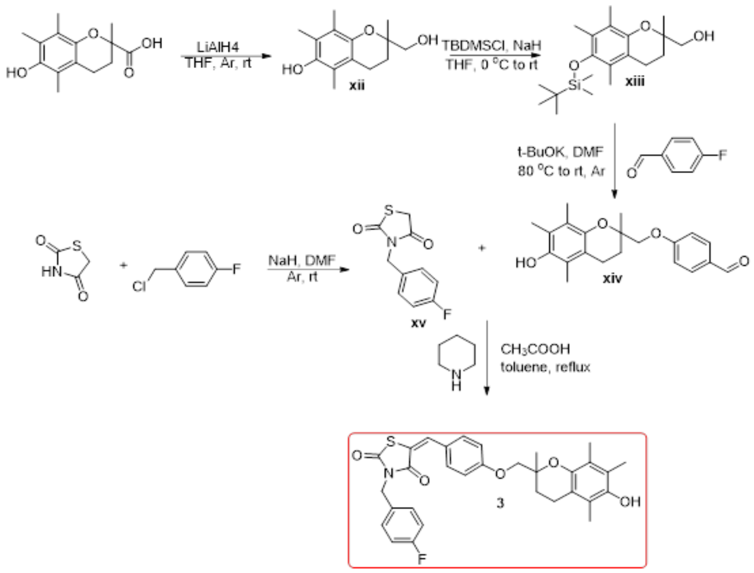


**Fig. S3. Synthetic route for compound 3.**

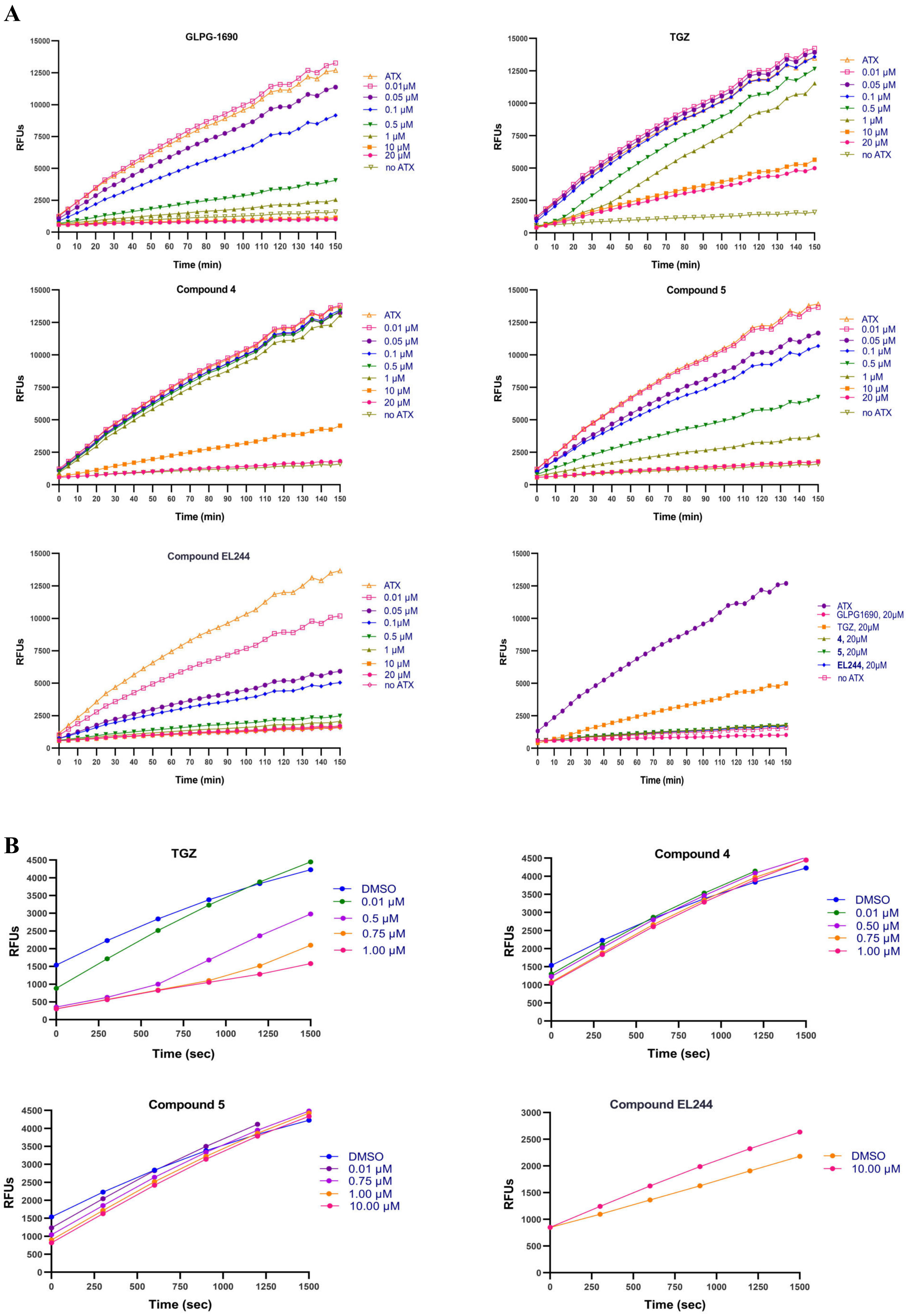


**Fig. S4. Kinetic analysis of ATX inhibition.** (**A**) Kinetic graphs of the inhibition of ATX by compounds 4, 5 and EL244, compared with the reference compounds GLPG-1690 and TGZ. (**B**) Kinetics for the second (oxidation of choline catalysed by choline oxidase) and third reaction (the conversion of the Amplex substrate to the fluorescent resorufin catalysed by horseradish peroxidase, HRP) of the human ATX Amplex Red Lyso-phospholipase D assay for representative compounds (TGZ, 4, 5 and EL244).

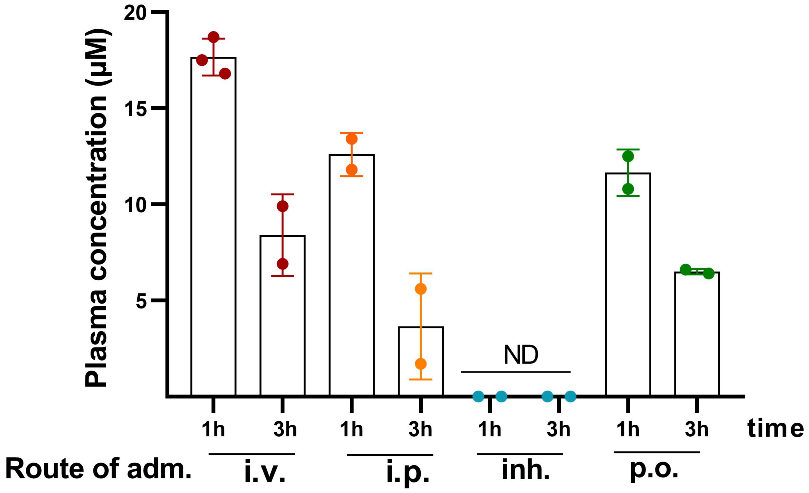


**Figure S5. Pharmacokinetic (PK) analysis of compound 5 reveals favourable systemic exposure.** Circulating plasma levels of compound 5 following different routes of administration (intravenous/i.v., intraperitoneal/i.p., inhaled/inh., and oral/p.o.) measured at 1- and 3-hours post-administration (30 mg/kg for i.v., i.p., p.o., 15 mg/kg for inhalation). ND: not detected, defined as plasma concentrations <50 nM.

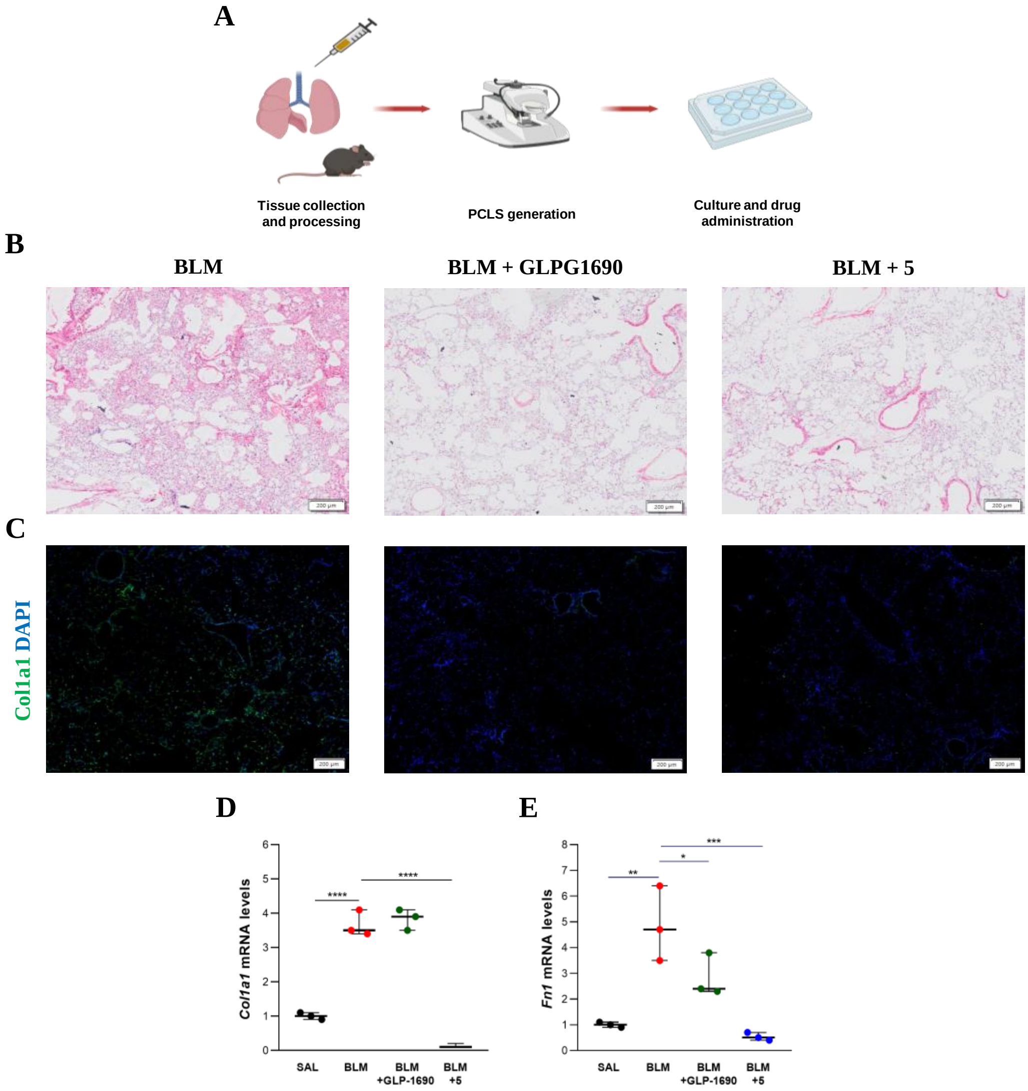


**Fig. S6. Compound 5 attenuates fibrosis in precision-cut lung slices (PCLS).** (**A**) Experimental set-up. Fibrotic PCLS were treated with compounds (30 µM) for 72h. (**B**) H&E staining in fibrotic slices treated with GLPG1690 and 5; scale bars 200µm. (**C**) Immunostaining for Col1a1 (green) and DAPI (blue); scale bars 200µm. (**D**) *Col1a1* and (**E**) *Fn1* mRNA levels were quantified with Q-RT-PCR; each sample is a pool of three slices. Values were normalized over the expression of *B2m* and presented as fold change over control. Statistical significance was assessed with one-way ANOVA.

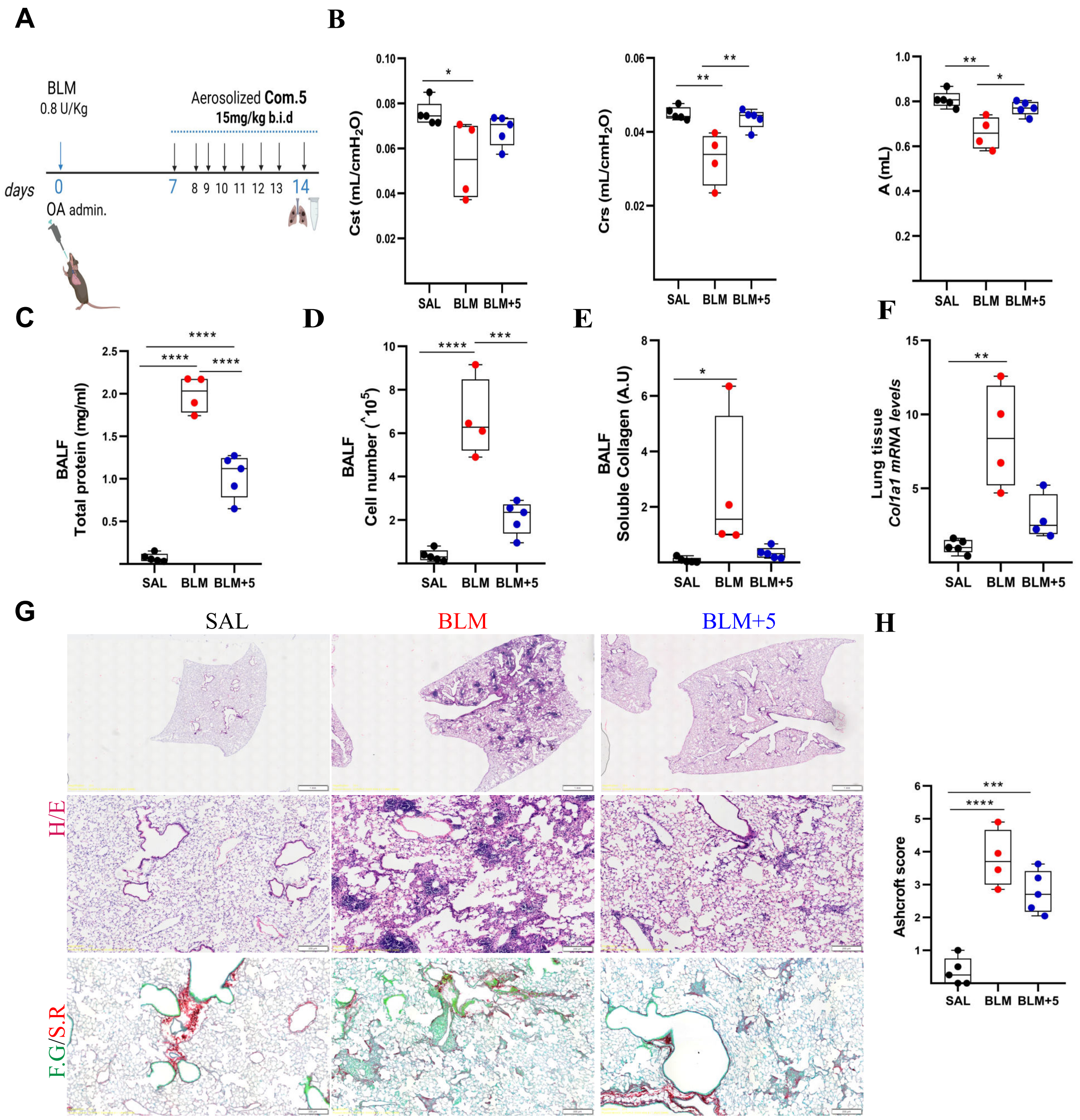


**Fig. S7. Compound 5 attenuates BLM-induced pulmonary fibrosis.** (**A**) Schematic representation of drug administration. (**B**) Respiratory mechanics assessed with FlexiVent. (**C**) Total protein concentration in BALFs, as determined with the Bradford assay. (**D**) Inflammatory cell numbers in BALFs, as counted with a haematocytometer. (E) Soluble collagen levels in the BALFs were detected with the direct red assay. (**F**) *Col1a1* mRNA expression was interrogated with Q-RT-PCR; values were normalised to the expression of *B2m* and presented as fold change over control. (**G**) Representative images of lung sections from mice of the indicated treatment groups, stained with H&E and Fast Green/Sirius Red (F.G./S.R.; green/red). (**H**) Quantification of fibrosis severity with the Ashcroft score. Following normality testing, statistical significance was assessed with Kruskal-Wallis and post-hoc Dunn's test (B/Cst) or one-way ANOVA and post-hoc Tukey's test (B/Crs, B/A, C, D, E, H) or Welch ANOVA and post-hoc Games-Howell test (F).

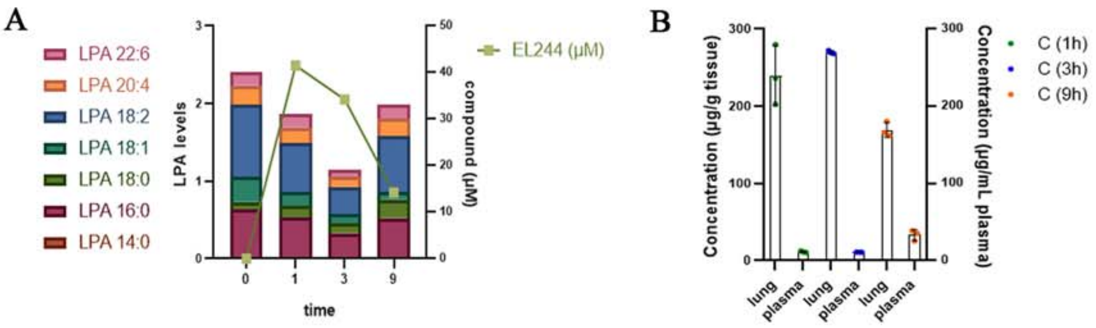


**Fig. S8. PK/PD analysis of EL244.** (**A**) EL244 concentrations and total LPA levels in the plasma, at 0, 1, 3 and 9h after its IP administration (30 mg/Kg). (**B**) EL244 concentrations in lung tissue and plasma, at 1, 3 and 9h after its inhaled administration (15 mg/Kg).

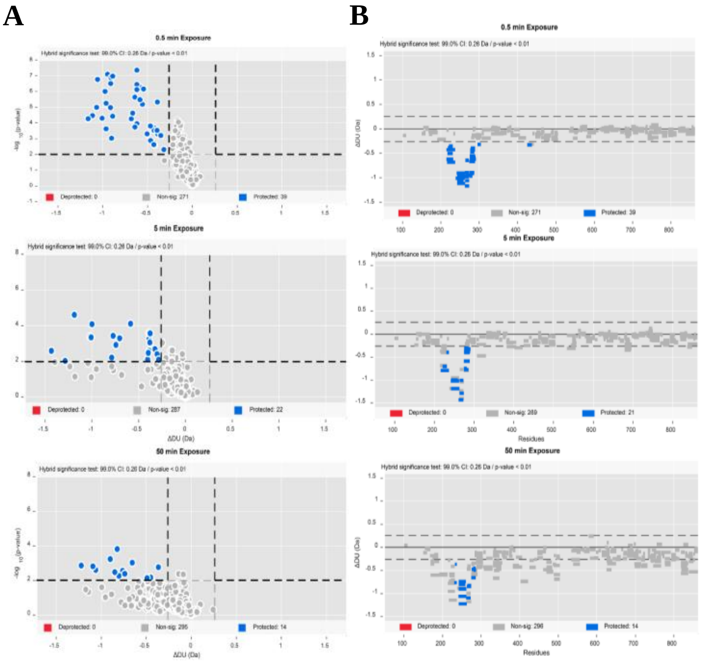


**Fig. S9. HDX-MS statistical analysis.** (**A**) Volcano plots for the identification of significant differences using α=0.01 (see Methods for details). Dots represent differential HDX differences, with blue indicating significant protection and light grey indicating non-significant differences. (**B**) Woodsplot representation with statistically significant peptides at each labelling time point. Peptides are represented as rectangles, with blue indicating significant protection and light gray indicating no differences. The x-axis denotes protein residues, while the y-axis shows the magnitude of the differences.

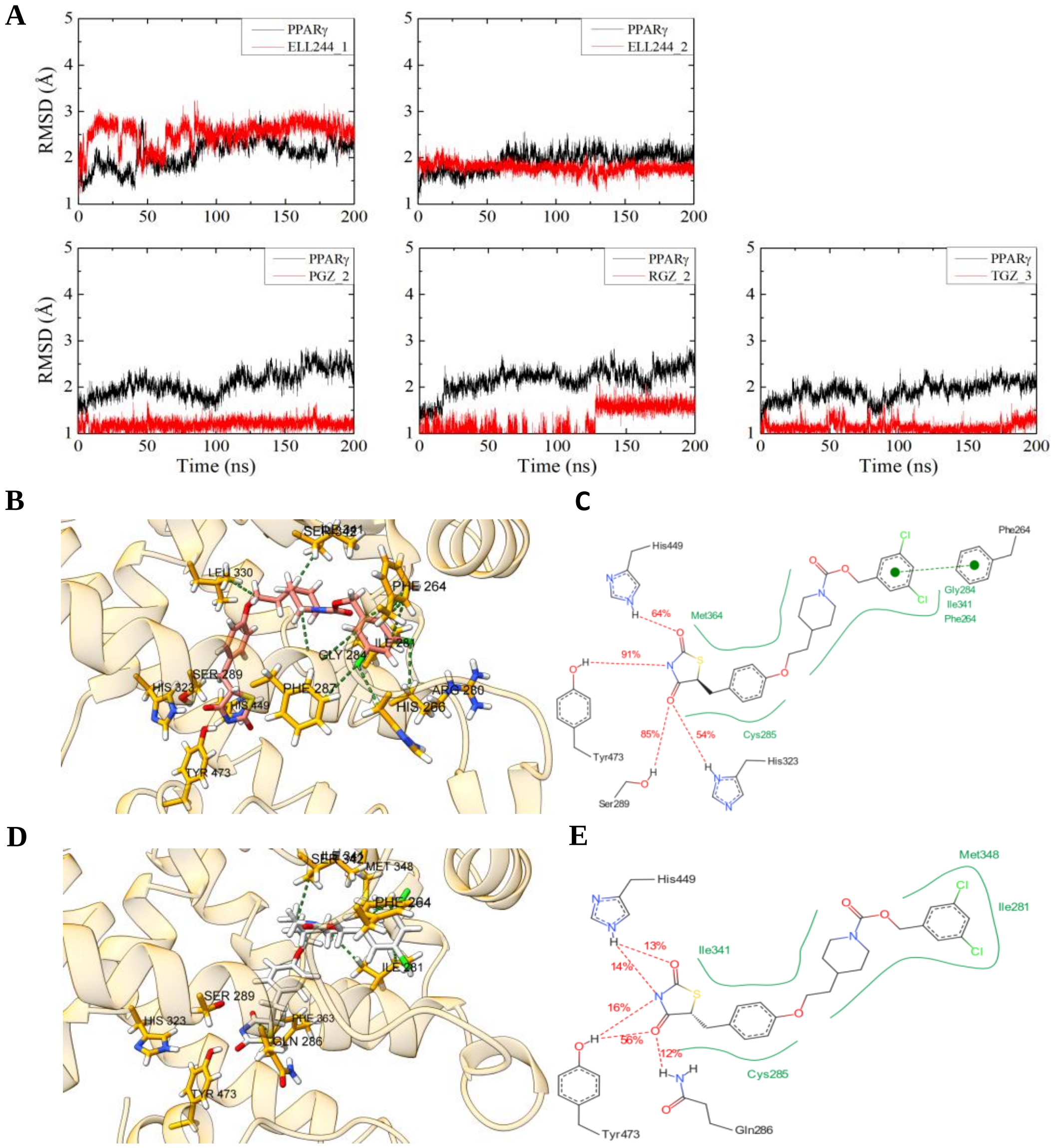


**Fig. S10. RMSD and MD simulation analysis of EL244 isomers binding to PPARγ.** (**A**) RMSD plots of the PPARγ protein systems investigated in their complex bound states with EL244 and TGZ. (**B**) Three- and (**C**) two- dimensional MD simulation representations of the EL244_2-PPARγ complex centroid conformation. (**D**) Three- and (**E**) two- dimensional MD simulation representations of the EL244_1-PPARγ complex centroid conformation.

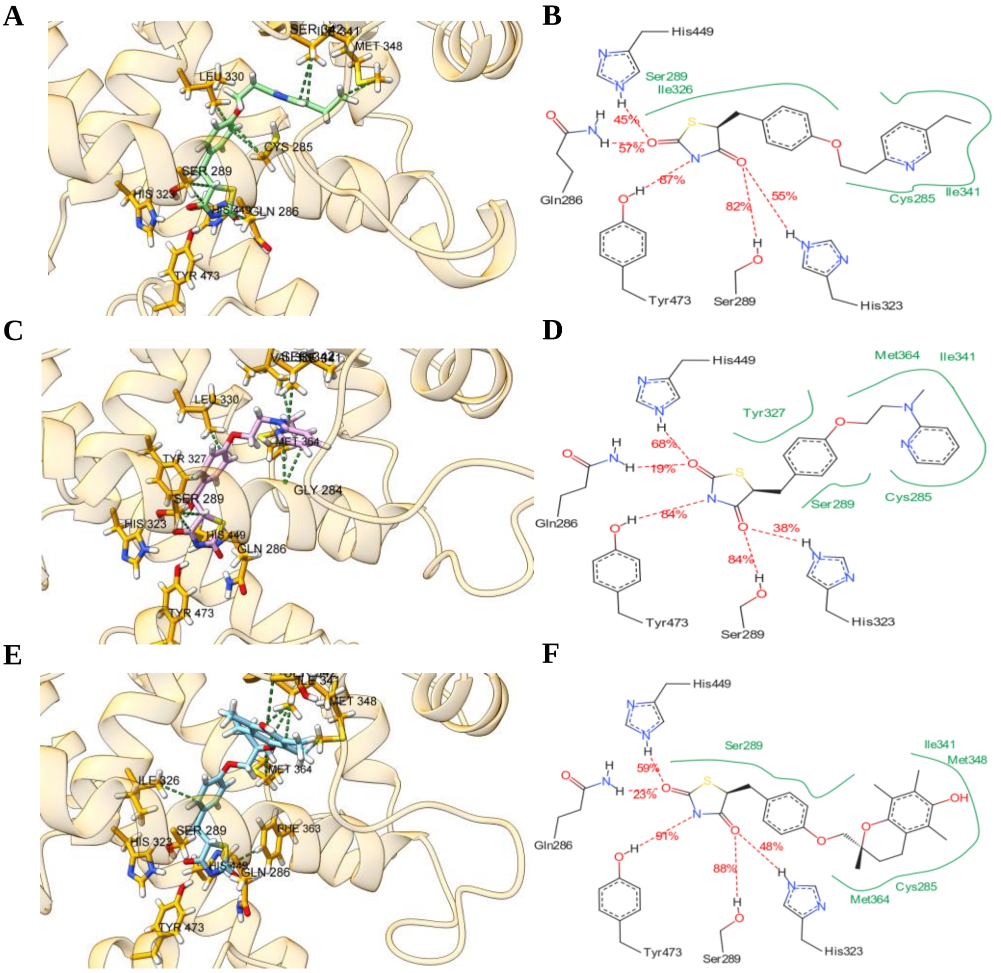


**Fig. S11. MD simulation analysis of TGZ isomers binding to PPARγ.** (**A**) Three- and (**B**) two- dimensional MD simulation representations of the PGZ_2-PPARγ complex centroid conformation. (**C**) Three- and (**D**) two- dimensional MD simulation representations of the RGZ_2-PPARγ complex centroid conformation. (**E**) Three- and (**F**) two- dimensional MD simulation representations of the TGZ_3-PPARγ complex centroid conformation.

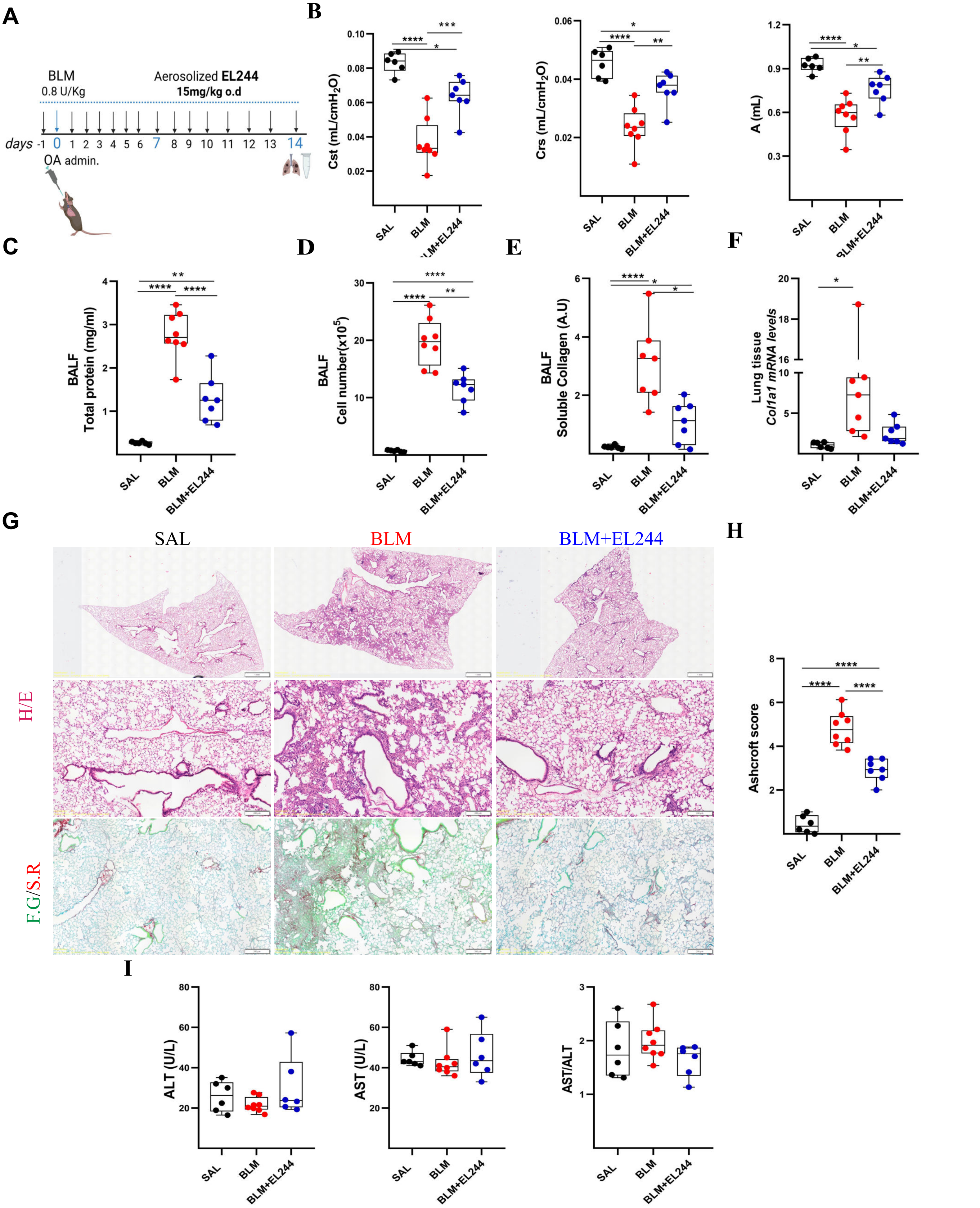


**Fig. S12. Inhaled prophylactic EL244 administration attenuates BLM-induced pulmonary fibrosis.** (**A**) Schematic representation of drug administration. (B) Respiratory mechanics were assessed using the FlexiVent system. (**C**) Total protein concentration in BALFs, as determined with the Bradford assay. (**D**) Inflammatory cell numbers in BALFs, as counted with a haematocytometer. (**E**) Soluble collagen levels in the BALFs were detected with the direct red assay; statistical significance was assessed with the Welch ANOVA test. (**F**) *Col1a1* mRNA expression was interrogated with Q-RT-PCR; values were normalised to the expression of *B2m* and presented as fold change over control. (**G**) Representative images of lung sections from mice of the indicated treatment groups, stained with H&E and Fast Green/Sirius Red (F.G/S.R; green/red). (**H**) Quantification of fibrosis severity in H&E-stained lung sections with the Ashcroft score. (**I**) Serum concentrations of alanine aminotransferase (ALT), aspartate aminotransferase (AST) and the AST/ALT ratio.Following normality testing, statistical significance was assessed with Kruskal-Wallis and post-hoc Dunn's test (I/AST) or one-way ANOVA and post-hoc Tukey's test (B, C) or Welch ANOVA and post-hoc Games-Howell test (D, E, F, H, I/ALT, ALT/AST). Scale bars 1,200 mm.
