## Supplementary Tables S1-S6 for "An aerosolised dual-action Autotaxin inhibitor-PPARγ agonist for the treatment of pulmonary fibrosis"

**Table S1. Top-ranked candidate ATX inhibitors.** Binding affinities are in kcal $mol^{-1}$ units. % inhibition at 100 μM.

| a/a | Compound (Prestw #) | Docking Score | % inhibition (IC50 μM) | a/a | Compound (Prestw #) | Docking Score | % inhibition (IC50 μM) |
|---|---|---|---|---|---|---|---|
| 1 | 1503 | -36.38 | 63.53 (14.73) | 26 | 627 | -25.43 | -4.13 (>100) |
| 2 | 425 | -31.95 | 99.21 (1.60) | 27 | 470 | -25.38 | 70.90 (35.09) |
| 3 | 400 | -31.06 | 78.24 (2.75) | 28 | 1217 | -25.26 | 26.23 (>100) |
| 4 | 1726 | -30.35 | 66.92 (53.05) | 29 | 814 | -25.20 | 9.41 (>100) |
| 5 | 437 | -30.16 | 5.32 (>100) | 30 | 340 | -25.15 | 22.27 (>100) |
| 6 | 1743 | -30.10 | 59.50 (57.06) | 31 | 539 | -25.06 | 8.74 (>100) |
| 7 | 656 | -29.16 | 13.38 (>100) | 32 | 1239 | -25.04 | 3.37 (>100) |
| 8 | 1190 | -28.88 | 26.69 (>100) | 33 | 1350 | -24.79 | 69.33 (62.76) |
| 9 | 327 | -28.73 | 13.87 (>100) | 34 | 1467 | -24.70 | 99.17 (0.53) |
| 10 | 991 | -28.72 | 74.27 (>39.38) | 35 | 1796 | -24.44 | 37.06 (>100) |
| 11 | 862 | -28.69 | 88.64 (3.51) | 36 | 482 | -24.34 | 31.18 (>100) |
| 12 | 316 | -28.50 | 48.82 (~100) | 37 | 131 | -24.19 | 5.40 (>100) |
| 13 | 1290 | -27.96 | 46.70 (~100) | 38 | 1737 | -24.11 | 26.64 (>100) |
| 14 | 939 | -27.72 | -5.03 (>100) | 39 | 819 | -24.00 | 35.57 (>100) |
| 15 | 473 | -27.19 | 18.19 (>100) | 40 | 389 | -23.73 | 41.89 (>100) |
| 16 | 1174 | -27.05 | 33.56 (>100) | 41 | 104 | -23.54 | 6.88 (>100) |
| 17 | 150 | -26.96 | 26.79 (>100) | 42 | 1270 | -23.34 | 6.72 (>100) |
| 18 | 143 | -26.19 | 31.78 (>72.5) | 43 | 1798 | -23.31 | 19.95 (>100) |
| 19 | 1188 | -26.06 | 29.81 (>100) | 44 | 587 | -22.97 | 74.79 (>51.92) |
| 20 | 1770 | -26.01 | 43.57 (>100) | 45 | 1793 | -22.91 | -7.94 (>100) |
| 21 | 869 | -25.81 | >100 | 46 | 1189 | -22.64 | 80.58 (>51.52) |
| 22 | 1494 | -25.79 | 23.20 (>100) | 47 | 285 | -22.55 | 46.28 (~100) |
| 23 | 149 | -25.62 | -7.37 (>100) | 48 | 1292 | -22.46 | 5.90 (>100) |
| 24 | 1794 | -25.55 | 19.33 (>100) | 49 | 378 | -22.20 | 13.22 (>100) |
| 25 | 1364 | -25.53 | 65.18 (>12.97) | | | | |

**Table S2. Per-residue MM–GBSA energy decomposition in the ATX complexes with tested compounds.** Only selected residues are shown. Uncertainties denote standard error of the mean (units in kcal mol-1) and are included in parentheses.

| Label | Residue | van der Waals | | Electrostatic | | Polar Solvation | | Total energy | |
|---|---|---|---|---|---|---|---|---|---|
| ATX-5_1 | Tyr82 | -1.82 | (0.02) | -0.64 | (0.01) | 1.32 | (0.01) | -1.43 | (0.02) |
| | Phe210 | -1.82 | (0.01) | -0.04 | (0.01) | 0.22 | (0.00) | -1.85 | (0.01) |
| | Leu213 | -0.91 | (0.01) | 0.92 | (0.00) | -0.86 | (0.00) | -0.91 | (0.01) |
| | Tyr214 | -1.87 | (0.01) | 0.46 | (0.01) | -0.03 | (0.00) | -1.62 | (0.01) |
| | Phe249 | -2.46 | (0.02) | 0.01 | (0.01) | 0.57 | (0.01) | -2.19 | (0.01) |
| | Trp254 | -3.86 | (0.02) | 0.33 | (0.01) | 0.45 | (0.01) | -3.57 | (0.02) |
| | Pro258 | -0.92 | (0.00) | -1.30 | (0.01) | 1.03 | (0.01) | -1.31 | (0.01) |
| | Phe273 | -1.81 | (0.01) | -1.12 | (0.01) | 1.49 | (0.01) | -1.68 | (0.01) |
| | Phe274 | -1.89 | (0.01) | -0.18 | (0.01) | 0.32 | (0.01) | -2.02 | (0.01) |
| ATX-5_2 | Phe210 | -2.82 | (0.01) | 0.24 | (0.01) | 0.07 | (0.01) | -2.82 | (0.01) |
| | Leu213 | -2.04 | (0.01) | 1.54 | (0.00) | -1.68 | (0.00) | -2.39 | (0.01) |
| | Ala217 | -1.15 | (0.01) | 0.79 | (0.01) | -0.77 | (0.00) | -1.23 | (0.01) |
| | Lys248 | -1.22 | (0.01) | 2.77 | (0.02) | -3.08 | (0.02) | -1.71 | (0.01) |
| | Phe249 | -1.88 | (0.01) | 0.16 | (0.00) | 0.08 | (0.00) | -1.85 | (0.01) |
| | Pro258 | -0.58 | (0.00) | -2.69 | (0.01) | 1.60 | (0.00) | -1.68 | (0.01) |
| | Leu259 | -0.84 | (0.00) | -3.81 | (0.01) | 2.56 | (0.01) | -2.12 | (0.01) |
| | Trp260 | -1.02 | (0.00) | -2.52 | (0.01) | 1.78 | (0.01) | -1.84 | (0.01) |
| | Phe273 | -2.51 | (0.01) | -1.70 | (0.01) | 2.56 | (0.01) | -1.82 | (0.01) |
| | Phe274 | -4.99 | (0.01) | -0.50 | (0.01) | 1.79 | (0.01) | -4.29 | (0.01) |
| | Trp275 | -1.18 | (0.01) | -4.53 | (0.02) | 4.20 | (0.01) | -1.59 | (0.01) |
| | Tyr306 | -2.33 | (0.01) | -1.88 | (0.02) | 1.12 | (0.01) | -3.31 | (0.02) |
| ATX-EL244_1 | Thr209 | -1.27 | (0.00) | -0.72 | (0.01) | 1.02 | (0.01) | -1.14 | (0.00) |
| | Phe210 | -1.61 | (0.00) | -0.39 | (0.00) | 0.69 | (0.00) | -1.46 | (0.00) |
| | Leu213 | -2.53 | (0.00) | -0.90 | (0.00) | 0.83 | (0.00) | -2.86 | (0.01) |
| | Leu243 | -0.98 | (0.00) | -0.47 | (0.00) | 0.57 | (0.00) | -1.14 | (0.00) |
| | Phe273 | -1.62 | (0.00) | -0.37 | (0.00) | 0.70 | (0.00) | -1.44 | (0.00) |
| | Phe274 | -2.60 | (0.01) | -2.49 | (0.01) | 1.61 | (0.00) | -3.74 | (0.01) |
| | Trp275 | -1.29 | (0.00) | -1.32 | (0.01) | 0.74 | (0.00) | -1.97 | (0.00) |
| | Tyr306 | -3.66 | (0.01) | -1.10 | (0.00) | 1.59 | (0.00) | -3.46 | (0.01) |
| ATX-EL244_2 | Phe210 | -1.46 | (0.01) | 0.21 | (0.01) | 0.16 | (0.01) | -1.31 | (0.00) |
| | Leu213 | -1.13 | (0.01) | -0.11 | (0.01) | 0.15 | (0.01) | -1.24 | (0.01) |
| | Phe249 | -1.92 | (0.01) | -0.09 | (0.01) | 0.63 | (0.01) | -1.59 | (0.01) |
| | Trp254 | -3.60 | (0.01) | -0.18 | (0.01) | 1.08 | (0.01) | -3.08 | (0.01) |
| | Trp260 | -1.02 | (0.01) | -1.11 | (0.01) | 1.33 | (0.01) | -0.94 | (0.01) |
| | Ile261 | -0.58 | (0.01) | -0.48 | (0.00) | 0.51 | (0.00) | -0.67 | (0.01) |
| | Phe274 | -3.79 | (0.01) | -3.43 | (0.03) | 3.35 | (0.02) | -4.53 | (0.02) |
| | Tyr306 | -1.92 | (0.01) | 0.85 | (0.01) | -0.52 | (0.01) | -1.82 | (0.01) |
| ATX-TGZ_1 | Lys208 | -1.18 | (0.01) | -31.52 | (0.12) | 29.56 | (0.11) | -3.24 | (0.02) |
| | Thr209 | -1.64 | (0.01) | -7.50 | (0.04) | 5.59 | (0.03) | -3.74 | (0.02) |
| | Phe210 | -2.48 | (0.01) | -4.77 | (0.03) | 3.39 | (0.01) | -4.14 | (0.01) |
| | Leu213 | -2.77 | (0.01) | -3.35 | (0.01) | 2.70 | (0.01) | -3.75 | (0.01) |
| | Leu243 | -0.92 | (0.01) | -0.06 | (0.01) | 0.16 | (0.01) | -1.00 | (0.01) |
| | Phe273 | -1.50 | (0.01) | 0.10 | (0.01) | 0.44 | (0.00) | -1.12 | (0.01) |
| | Phe274 | -2.59 | (0.01) | -0.78 | (0.00) | 1.16 | (0.00) | -2.46 | (0.01) |
| | Tyr306 | -1.91 | (0.01) | -0.45 | (0.00) | 0.99 | (0.00) | -1.62 | (0.01) |
| | Leu78 | -1.07 | (0.00) | -0.60 | (0.00) | 0.67 | (0.00) | -1.18 | (0.00) |

| | | | | | | | | | |
|---|---|---|---|---|---|---|---|---|---|
| **ATX-TGZ_2** | Phe210 | -2.18 | (0.01) | -0.45 | (0.00) | 0.78 | (0.00) | -2.12 | (0.01) |
| | Tyr214 | -1.59 | (0.01) | 0.21 | (0.00) | 0.05 | (0.00) | -1.45 | (0.01) |
| | Lys248 | -1.61 | (0.01) | -16.12 | (0.02) | 16.30 | (0.02) | -1.60 | (0.01) |
| | Phe249 | -2.33 | (0.01) | 1.22 | (0.01) | -0.55 | (0.01) | -1.94 | (0.01) |
| | His251 | -0.49 | (0.01) | -4.95 | (0.04) | 4.47 | (0.03) | -1.08 | (0.01) |
| | Trp254 | -2.27 | (0.01) | -0.58 | (0.01) | 1.33 | (0.01) | -1.81 | (0.01) |
| | Pro258 | -1.08 | (0.00) | -0.67 | (0.00) | 0.59 | (0.00) | -1.26 | (0.01) |
| | Trp260 | -2.30 | (0.01) | -0.08 | (0.01) | 0.59 | (0.01) | -2.10 | (0.01) |
| | Phe274 | -3.23 | (0.01) | 0.21 | (0.01) | 0.51 | (0.00) | -3.00 | (0.01) |
| **ATX-TGZ_3** | Leu78 | -1.04 | (0.01) | -0.01 | (0.00) | 0.11 | (0.00) | -1.09 | (0.01) |
| | Phe210 | -1.69 | (0.01) | -0.05 | (0.00) | 0.34 | (0.00) | -1.69 | (0.01) |
| | Leu243 | -2.29 | (0.01) | 1.42 | (0.01) | -0.97 | (0.01) | -2.08 | (0.01) |
| | Arg244 | -1.68 | (0.01) | -10.80 | (0.02) | 11.25 | (0.03) | -1.52 | (0.01) |
| | Lys248 | -2.54 | (0.01) | -23.77 | (0.05) | 23.95 | (0.04) | -2.66 | (0.01) |
| | Phe249 | -1.86 | (0.01) | 0.85 | (0.02) | -0.51 | (0.01) | -1.73 | (0.01) |
| | Trp254 | -1.90 | (0.01) | 0.16 | (0.01) | 0.40 | (0.01) | -1.54 | (0.01) |
| | Phe274 | -1.84 | (0.01) | -0.53 | (0.01) | 0.88 | (0.01) | -1.71 | (0.01) |
| **ATX-TGZ_4** | Phe210 | -1.90 | (0.01) | -0.50 | (0.01) | 0.85 | (0.01) | -1.86 | (0.01) |
| | Leu213 | -2.40 | (0.01) | -2.55 | (0.01) | 1.81 | (0.01) | -3.44 | (0.01) |
| | Tyr214 | -1.10 | (0.00) | -0.38 | (0.00) | 0.48 | (0.00) | -1.06 | (0.00) |
| | Leu243 | -1.19 | (0.01) | 1.53 | (0.02) | -1.29 | (0.02) | -1.10 | (0.01) |
| | Arg244 | -0.99 | (0.01) | -29.39 | (0.15) | 27.64 | (0.12) | -2.92 | (0.03) |
| | Lys248 | -0.41 | (0.01) | -32.00 | (0.15) | 30.49 | (0.13) | -2.05 | (0.03) |
| | Phe273 | -1.41 | (0.01) | 0.01 | (0.00) | 0.42 | (0.00) | -1.12 | (0.01) |
| | Phe274 | -3.24 | (0.01) | -1.92 | (0.01) | 1.90 | (0.00) | -3.58 | (0.01) |
| | Tyr306 | -2.04 | (0.01) | 0.12 | (0.00) | 0.24 | (0.00) | -1.91 | (0.01) |

**Table S3. Structures and total binding energies ($\Delta G_{bind}$) with ATX and PPAR$_\gamma$ complexes of tested compounds**, as calculated with the MM–GBSA method. Standard error of the mean is shown in parentheses. The suggested (by MD simulations) type of inhibition of the compounds against ATX is also included.

| Compound | Label | Structure | $\Delta G_{bind}$ (kcal mol$^{-1}$) / ATX | $\Delta G_{bind}$ (kcal mol– 1) / PPARγ | ATX inhibitor type |
|---|---|---|---|---|---|
| Troglitazone (TGZ) | TGZ_1 | 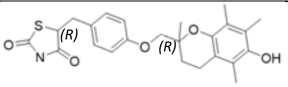 | -54.21 (0.52) | | Type II |
| | TGZ_2 | 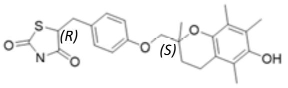 | -40.51 (0.29) | | |
| | TGZ_3 | 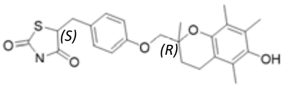 | -35.61 (0.41) | -58.01 (0.21) | |
| | TGZ_4 | 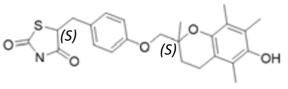 | -53.59 (0.43) | | Type II |
| 5 | 5_1 | 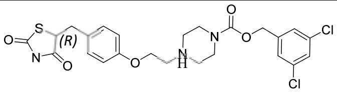 | -40.84 (0.23) | -63.40 (0.17) | Type III |
| | 5_2 | 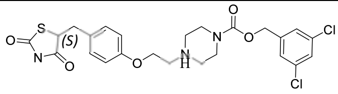 | -60.86 (0.22) | -60.17 (0.20) | Type IV |
| EL244 | EL244_1 | 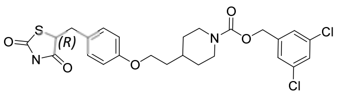 | -76.62 (0.23) | -66.23 (0.32) | Type I |
| | EL244_2 | 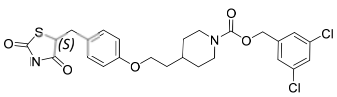 | -46.01 (0.26) | -71.01 (0.16) | Type IV |
| Rosiglitazone (RGZ) | RGZ_1 | 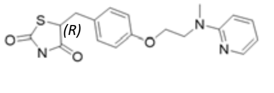 | | | |
| | RGZ_2 | 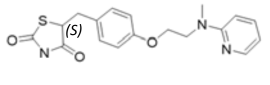 | | -47.23 (0.16) | |

| | | | | |
|---|---|---|---|---|
| **Pioglitazone (PGZ)** | PGZ_1 | 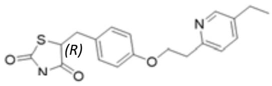 | | |
| | PGZ_2 | 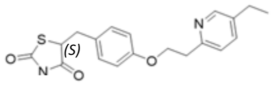 | | -49.76 (0.18) |

**Table S4. Critical HDX-MS experimental information.**

| **Data Set** | **ATX** | **ATX + EL244** |
|---|---|---|
| HDX reaction details | 10mM PBS, 150mM NaCl, pD=7.4 at 25°C | 10mM PBS, 150mM NaCl, pD=7.4 at 25°C |
| HDX time course (s) | 30, 300, 3000 | 30, 300, 3000 |
| HDX control samples | none | none |
| Deuterium recovery (mean) | 70%* | |
| # of Peptides | 310 | 310 |
| Sequence coverage | 84.54% | 84.54% |
| Average peptide length / Redundancy | 12.48/5.62 | 12.48/5.62 |
| Replicates (biological or technical) | 4 (Technical) | 4 (Technical) |
| Repeatability | 0.0482 (average SD) | 0.0629 (average SD) |
| Significant differences in HDX (delta HDX > X D) | 0.26Da | 0.26Da |

** Average deuterium recovery calculated using RRPYIL, DRVYIHPF and RPKPQQFFGLM-NH2 as model peptides.*

**Table S5. Calculated occurrence of water bridges observed in the ATX complexes examined.** Only compounds where water bridges occur higher than 25% of the simulation time are shown.

| Compound | ATX residues involved | Occurrence (%) |
|---|---|---|
| EL244_1 | Asp311 | 46 |
| EL244_2 | Trp275 | 53 |
| TGZ_1 | – | – |
| TGZ_2 | Glu67 | 60 |
| | Trp260 | 59 |
| | Arg74 | 27 |
| TGZ_3 | Thr272 | 37 |
| TGZ_4 | – | – |

**Table S6. Per-residue MM–GBSA energy decomposition in the PPAR$_{\gamma}$ complexes examined.** Only selected residues are shown. Uncertainties denote standard error of the mean (units in kcal mol$^{-1}$) and are included in parentheses.

| Label | Residue | van der Waals | | Electrostatic | | Polar Solvation | | Total energy | |
|---|---|---|---|---|---|---|---|---|---|
| **PPAR*γ*-5_1** | Cys285 | -1.61 | (0.01) | -0.18 | (0.01) | 0.57 | (0.01) | -1.38 | (0.01) |
| | Gln286 | -2.02 | (0.01) | -7.91 | (0.02) | 7.71 | (0.02) | -2.46 | (0.01) |
| | Arg288 | -1.28 | (0.00) | 2.10 | (0.01) | -2.10 | (0.01) | -1.49 | (0.00) |
| | Ser289 | -2.07 | (0.01) | -1.04 | (0.01) | 1.52 | (0.01) | -1.78 | (0.01) |
| | His323 | -2.28 | (0.01) | -7.97 | (0.02) | 7.96 | (0.01) | -2.56 | (0.01) |
| | Ile326 | -2.87 | (0.01) | -0.04 | (0.01) | 0.02 | (0.01) | -3.19 | (0.01) |
| | Tyr327 | -1.75 | (0.01) | -1.21 | (0.01) | 1.22 | (0.01) | -1.89 | (0.01) |
| | Leu330 | -1.42 | (0.00) | 0.36 | (0.00) | -0.37 | (0.00) | -1.58 | (0.00) |
| | Phe363 | -1.76 | (0.01) | -2.15 | (0.02) | 2.26 | (0.02) | -1.79 | (0.01) |
| | Met364 | -1.51 | (0.01) | -0.39 | (0.01) | 0.39 | (0.01) | -1.64 | (0.01) |
| | Leu465 | -1.47 | (0.00) | -3.73 | (0.01) | 2.85 | (0.00) | -2.42 | (0.01) |
| | His466 | -1.08 | (0.01) | -0.52 | (0.01) | -0.43 | (0.01) | -2.11 | (0.01) |
| | Leu469 | -1.38 | (0.00) | 0.11 | (0.01) | 0.35 | (0.00) | -1.00 | (0.01) |
| | Gln470 | -1.22 | (0.00) | -1.84 | (0.01) | 1.73 | (0.01) | -1.37 | (0.00) |
| **PPAR*γ*-5_2** | Cys285 | -3.12 | (0.01) | -2.66 | (0.01) | 3.02 | (0.01) | -3.21 | (0.01) |
| | Gln286 | -1.42 | (0.00) | -1.46 | (0.03) | 1.35 | (0.02) | -1.58 | (0.01) |
| | Arg288 | -1.96 | (0.01) | 10.74 | (0.02) | -10.19 | (0.02) | -1.69 | (0.01) |
| | Ser289 | -0.46 | (0.01) | -7.34 | (0.02) | 6.49 | (0.01) | -1.37 | (0.01) |
| | His323 | -0.61 | (0.01) | -6.91 | (0.01) | 4.94 | (0.01) | -2.63 | (0.01) |
| | Ile326 | -1.52 | (0.00) | -1.79 | (0.00) | 1.81 | (0.00) | -1.74 | (0.01) |
| | Leu330 | -1.83 | (0.01) | -0.59 | (0.00) | 0.64 | (0.00) | -2.08 | (0.01) |
| | Val339 | -1.27 | (0.01) | 0.75 | (0.00) | -0.67 | (0.00) | -1.30 | (0.01) |
| | Ile341 | -1.71 | (0.00) | 1.02 | (0.01) | -0.51 | (0.00) | -1.42 | (0.00) |
| | Phe363 | -1.11 | (0.01) | -0.36 | (0.00) | 0.52 | (0.00) | -1.03 | (0.00) |
| | Met364 | -2.32 | (0.01) | -0.78 | (0.00) | 1.10 | (0.00) | -2.20 | (0.01) |
| | Lys367 | -0.96 | (0.00) | -5.72 | (0.01) | 5.70 | (0.01) | -1.03 | (0.00) |
| | His449 | -0.66 | (0.01) | -6.18 | (0.02) | 4.24 | (0.01) | -2.65 | (0.01) |
| | Tyr473 | -0.21 | (0.01) | -6.35 | (0.01) | 5.66 | (0.01) | -0.95 | (0.01) |
| **PPAR*γ*-EL244_1** | Arg280 | -1.09 | (0.00) | -10.7 | (0.01) | 10.38 | (0.01) | -1.49 | (0.00) |
| | Ile281 | -2.83 | (0.01) | 0.38 | (0.00) | -0.06 | (0.00) | -2.75 | (0.01) |
| | Phe282 | -2.62 | (0.01) | 0.61 | (0.01) | -0.25 | (0.00) | -2.38 | (0.01) |
| | Cys285 | -2.74 | (0.01) | -0.65 | (0.01) | 1.35 | (0.01) | -2.53 | (0.01) |
| | Gln286 | -1.17 | (0.01) | -5.91 | (0.02) | 4.40 | (0.01) | -2.78 | (0.01) |
| | Ile341 | -2.21 | (0.01) | -0.72 | (0.00) | 0.77 | (0.00) | -2.45 | (0.01) |
| | Met348 | -1.37 | (0.01) | -0.12 | (0.00) | 0.22 | (0.00) | -1.38 | (0.01) |
| | Phe363 | -1.31 | (0.01) | 0.27 | (0.01) | -0.12 | (0.01) | -1.26 | (0.01) |
| | Met364 | -1.29 | (0.00) | -0.08 | (0.00) | 0.14 | (0.00) | -1.36 | (0.00) |
| | Lys367 | -1.06 | (0.01) | -32.5 | (0.11) | 30.05 | (0.09) | -3.68 | (0.03) |
| | His449 | -0.44 | (0.01) | -5.66 | (0.04) | 3.92 | (0.03) | -2.26 | (0.02) |
| | Tyr473 | -0.05 | (0.01) | -5.46 | (0.04) | 5.01 | (0.03) | -0.55 | (0.01) |
| | Phe264 | -2.67 | (0.00) | -0.60 | (0.00) | 0.81 | (0.00) | -2.70 | (0.00) |
| | Arg280 | -1.42 | (0.00) | -10.6 | (0.01) | 10.79 | (0.01) | -1.40 | (0.00) |
| | Ile281 | -2.38 | (0.01) | 0.40 | (0.00) | -0.25 | (0.00) | -2.39 | (0.01) |
| | Gly284 | -1.41 | (0.01) | -0.39 | (0.00) | 0.60 | (0.00) | -1.42 | (0.01) |
| | Cys285 | -3.30 | (0.01) | -2.33 | (0.01) | 2.16 | (0.01) | -3.77 | (0.01) |
| | Gln286 | -1.63 | (0.00) | -2.13 | (0.03) | 1.90 | (0.02) | -1.91 | (0.01) |

| | | | | | | | | | |
|---|---|---|---|---|---|---|---|---|---|
| **PPAR*γ*-EL244_2** | Arg288 | -1.12 | (0.01) | -12.8 | (0.01) | 12.91 | (0.01) | -1.29 | (0.01) |
| | Ser289 | -0.30 | (0.01) | -7.63 | (0.02) | 6.85 | (0.01) | -1.15 | (0.01) |
| | His323 | -0.39 | (0.01) | -7.37 | (0.02) | 5.05 | (0.01) | -2.74 | (0.01) |
| | Leu330 | -1.15 | (0.00) | -0.73 | (0.00) | 0.77 | (0.00) | -1.29 | (0.00) |
| | Ile341 | -1.91 | (0.01) | -0.11 | (0.00) | 0.28 | (0.00) | -2.01 | (0.01) |
| | Met348 | -1.10 | (0.00) | -0.66 | (0.01) | 0.55 | (0.00) | -1.28 | (0.00) |
| | Met364 | -1.16 | (0.00) | 0.10 | (0.01) | 0.07 | (0.01) | -1.11 | (0.00) |
| | His449 | -0.92 | (0.01) | -7.10 | (0.01) | 5.02 | (0.01) | -3.06 | (0.01) |
| | Tyr473 | -0.11 | (0.01) | -7.28 | (0.01) | 6.29 | (0.01) | -1.14 | (0.01) |
| **PPAR*γ*-PGZ_2** | Cys285 | -3.49 | (0.01) | -1.70 | (0.01) | 1.93 | (0.01) | -3.59 | (0.01) |
| | Gln286 | -1.76 | (0.01) | -5.55 | (0.03) | 4.12 | (0.02) | -3.24 | (0.01) |
| | Arg288 | -1.34 | (0.00) | -15.2 | (0.01) | 15.42 | (0.01) | -1.43 | (0.00) |
| | Ser289 | -0.46 | (0.01) | -7.84 | (0.02) | 7.05 | (0.01) | -1.32 | (0.01) |
| | His323 | -0.37 | (0.01) | -7.77 | (0.02) | 5.26 | (0.01) | -2.92 | (0.01) |
| | Leu330 | -1.30 | (0.00) | -0.73 | (0.00) | 0.76 | (0.00) | -1.46 | (0.00) |
| | Ile341 | -1.82 | (0.01) | -0.98 | (0.00) | 0.94 | (0.00) | -2.11 | (0.01) |
| | His449 | -0.62 | (0.01) | -6.38 | (0.01) | 4.72 | (0.01) | -2.31 | (0.00) |
| | Tyr473 | -0.12 | (0.01) | -7.09 | (0.01) | 6.22 | (0.01) | -1.03 | (0.01) |
| **PPAR*γ*-RGZ_2** | Gly284 | -0.86 | (0.00) | -0.15 | (0.01) | -0.03 | (0.00) | -1.16 | (0.00) |
| | Cys285 | -3.56 | (0.01) | -2.09 | (0.01) | 2.18 | (0.01) | -3.79 | (0.01) |
| | Gln286 | -1.59 | (0.00) | -2.83 | (0.04) | 2.30 | (0.02) | -2.16 | (0.02) |
| | Arg288 | -1.33 | (0.00) | -15.2 | (0.01) | 15.25 | (0.01) | -1.54 | (0.01) |
| | Ser289 | -0.32 | (0.01) | -7.71 | (0.01) | 6.91 | (0.01) | -1.18 | (0.01) |
| | His323 | -0.48 | (0.01) | -7.18 | (0.01) | 5.06 | (0.01) | -2.62 | (0.01) |
| | Ile326 | -1.09 | (0.00) | -0.82 | (0.00) | 0.85 | (0.00) | -1.19 | (0.00) |
| | Ile341 | -1.61 | (0.01) | -0.85 | (0.00) | 0.82 | (0.00) | -1.86 | (0.01) |
| | His449 | -0.60 | (0.01) | -7.16 | (0.01) | 4.94 | (0.01) | -2.86 | (0.01) |
| | Tyr473 | -0.05 | (0.01) | -7.32 | (0.01) | 6.41 | (0.01) | -1.01 | (0.01) |
| **PPAR*γ*-TGZ_3** | Gly284 | -1.10 | (0.01) | -0.16 | (0.01) | 0.21 | (0.01) | -1.20 | (0.01) |
| | Cys285 | -3.54 | (0.01) | -1.89 | (0.01) | 1.89 | (0.01) | -3.86 | (0.01) |
| | Gln286 | -1.62 | (0.01) | -3.38 | (0.04) | 2.76 | (0.02) | -2.28 | (0.02) |
| | Arg288 | -2.03 | (0.01) | -13.8 | (0.01) | 14.17 | (0.01) | -2.08 | (0.01) |
| | Ser289 | -0.38 | (0.01) | -7.48 | (0.01) | 6.81 | (0.01) | -1.12 | (0.01) |
| | His323 | -0.47 | (0.01) | -7.10 | (0.01) | 4.97 | (0.01) | -2.62 | (0.01) |
| | Ile326 | -1.07 | (0.00) | -0.81 | (0.00) | 0.85 | (0.00) | -1.16 | (0.00) |
| | Leu330 | -1.17 | (0.00) | -0.68 | (0.00) | 0.72 | (0.00) | -1.31 | (0.00) |
| | Ile341 | -2.61 | (0.01) | -0.89 | (0.00) | 0.98 | (0.00) | -2.77 | (0.01) |
| | His449 | -0.61 | (0.01) | -7.03 | (0.01) | 4.87 | (0.01) | -2.81 | (0.01) |
| | Tyr473 | -0.10 | (0.01) | -7.40 | (0.01) | 6.38 | (0.01) | -1.16 | (0.01) |
